# Function-guided design of active enzymes

**DOI:** 10.64898/2026.06.27.735025

**Authors:** Mingyang Hu, Lunjie Wu, Yi Yang, Feiran Li, Linchao Zhu

## Abstract

Designing enzymes from functional descriptions remains challenging because catalytic activity is governed by sequence–structure–function relationships. Here we present EnzymeArt, a function-conditioned enzyme-design framework centred on a generative sequence model. EnzymeArt couples function-conditioned sequence generation with structure-guided refinement, annotation checks and substrate-aware computational prioritization to select candidates for synthesis and biochemical testing. Across alcohol dehydrogenase (ADH), malate dehydrogenase (MDH) and triacylglycerol lipase design campaigns, 57 of 60 synthesized designs showed crude-lysate activity above matched background controls. Purified representatives further showed quantitative steady-state catalytic activity. The best designed ADH reached ***k*_cat_ = 223.7 s*^−^*^1^** and exceeded a wild-type reference under matched conditions, an MDH reached ***k*_cat_ = 267.57 s*^−^*^1^** despite having only 33% sequence identity to its closest BLASTP hit, and a designed lipase hydrolysed both short– and long-chain triglycerides with apparent activity modestly above that of a commercial lipase reference. Together, these results establish a route for converting functional descriptions into experimentally validated enzyme designs with quantitative steady-state kinetic activity.

## 1 Introduction

Designing enzymes from functional specifications is a long-standing goal in protein engineering. Yet catalytic activity requires more than assigning an enzyme name or recovering a plausible fold. Productive catalysis depends on the joint compatibility of sequence, fold, active-site chemistry, substrate recognition and the surrounding protein environment. Although generative models can now produce protein sequences and structures with high predicted confidence, converting such plausibility into experimentally measurable catalysis remains difficult, particularly when the design objective is specified at the level of function rather than an explicit active-site geometry [1, 2].

Previous experimental successes in de novo enzyme design have largely relied on workflows based on theozymes [3], in which an active-site geometry is first specified and then transplanted into compatible protein scaffolds. Early landmark studies demonstrated that such strategies can yield enzymes with measurable activity [4, 5]. However, these activities are typically orders of magnitude lower than those of natural enzymes [6, 7], reflecting the difficulty of reconciling atomic-level geometry with globally coherent protein folds [8, 9].

In parallel, structure-based generative models have substantially advanced de novo protein design. Diffusion-based frameworks, such as RFdiffusion [10] and Chroma [11], have expanded the accessible design space by enabling backbone-level motif scaffolding [12]. However, these approaches typically separate backbone generation from side-chain packing [13], making it difficult to explicitly optimize strict atom-level side-chain geometries required by theozymes during the generative process. Moreover, efforts to combine atom-level active-site enumeration with backbone scaffolding [14] remain computationally intensive and scale poorly with active-site complexity. As a result, typical workflows [15] treat backbone generation and sequence optimization as distinct stages (e.g., using ProteinMPNN [16]), a separation that often compounds errors and yields candidates that are well-folded yet inactive [17]. Similarly, symbolic approaches such as Pinal [18], which generate discrete structural tokens prior to sequence decoding, rely on intermediate representations that can amplify upstream uncertainties.

Sequence-based generative strategies offer an appealing alternative but face challenges intrinsic to protein sequences [19]. Unlike natural language, protein sequences exhibit strong bidirectional dependencies; information from both termini shapes the local and global structural context. Autoregressive approaches such as ZymC-TRL [20] and ProGen2 [21] capture only unidirectional context [22–24], whereas masked language models provide bidirectional context and partially alleviate this limitation [25–28]. More recently, multimodal models [29–32] such as ESM3 [33] have integrated sequence, structure, and function representations to enable controllable protein generation.

However, such functional annotations do not by themselves specify substrate-dependent catalytic performance or provide a direct criterion for selecting designs with high measured turnover. This makes candidate prioritization a central bottleneck: generative models can produce many sequences or structures that appear plausible by confidence, structural or annotation-based metrics, but these metrics do not necessarily identify which candidates should be synthesized or which are most likely to show favourable substrate-specific kinetics. Thus, translating functional prompts into experimentally testable enzyme designs remains insufficiently coupled to quantitative kinetic objectives.

ZymCTRL, for example, demonstrated that conditioning sequence generation on EC numbers can yield enzymes with experimentally detectable activity [20]. However, EC-level conditioning alone does not specify substrate-dependent turnover, nor does it provide a quantitative criterion for prioritizing generated sequences for biochemical testing. More generally, existing frameworks provide limited guidance for translating functional prompts into experimentally testable candidates and for ranking those candidates toward steady-state kinetic outcomes [34, 35].

Here we introduce EnzymeArt, an integrated framework for designing enzymes from functional descriptions. EnzymeArt combines function-conditioned sequence generation with sequence and structure refinement, domain-annotation filtering and substrate-aware kinetic ranking. We applied this workflow to alcohol dehydrogenases, malate dehydrogenases and triacylglycerol lipases, spanning nicotinamide-dependent redox chemistry and lipid hydrolysis.

Across these three enzyme-design campaigns, 57 of 60 synthesized candidates showed crude-lysate activity above matched background controls. We then purified representative candidates for steady-state kinetic characterization. These included an alcohol dehydrogenase that exceeded a wild-type reference under matched conditions, a malate dehydrogenase with only 33% sequence identity to its closest BLASTP hit, and a lipase whose apparent activity was modestly higher than that of a commercial benchmark. These results suggest that functional prompting can support experimentally grounded enzyme design when sequence generation is coupled to substrate-aware selection and biochemical validation.

## 2 Results

### 2.1 Function-guided enzyme design with EnzymeArt

EnzymeArt is a function-guided generative framework for enzyme design. Starting from a functional description and a fully or partially masked sequence, EnzymeArt generates enzyme sequences, refines them through sequence–structure optimization, and prioritizes candidates using complementary computational criteria before experimental testing. The framework links high-level functional specifications to molecular-level assessment criteria, including predicted fold confidence, sequence plausibility, domain organization and substrate-conditioned kinetic ranking.

EnzymeArt is organized into three stages of design, optimization and evaluation (Fig. 1A). In the design stage, the generator uses a Transformer backbone to model protein sequence patterns and incorporates the functional description through a sequence–function interaction module (SFIM; Fig. 1B; Methods 4.5, 4.2.3). This conditioning allows functional information to modulate residue selection during iterative decoding, producing an initial pool of candidate enzyme sequences.

**Fig. 1.**
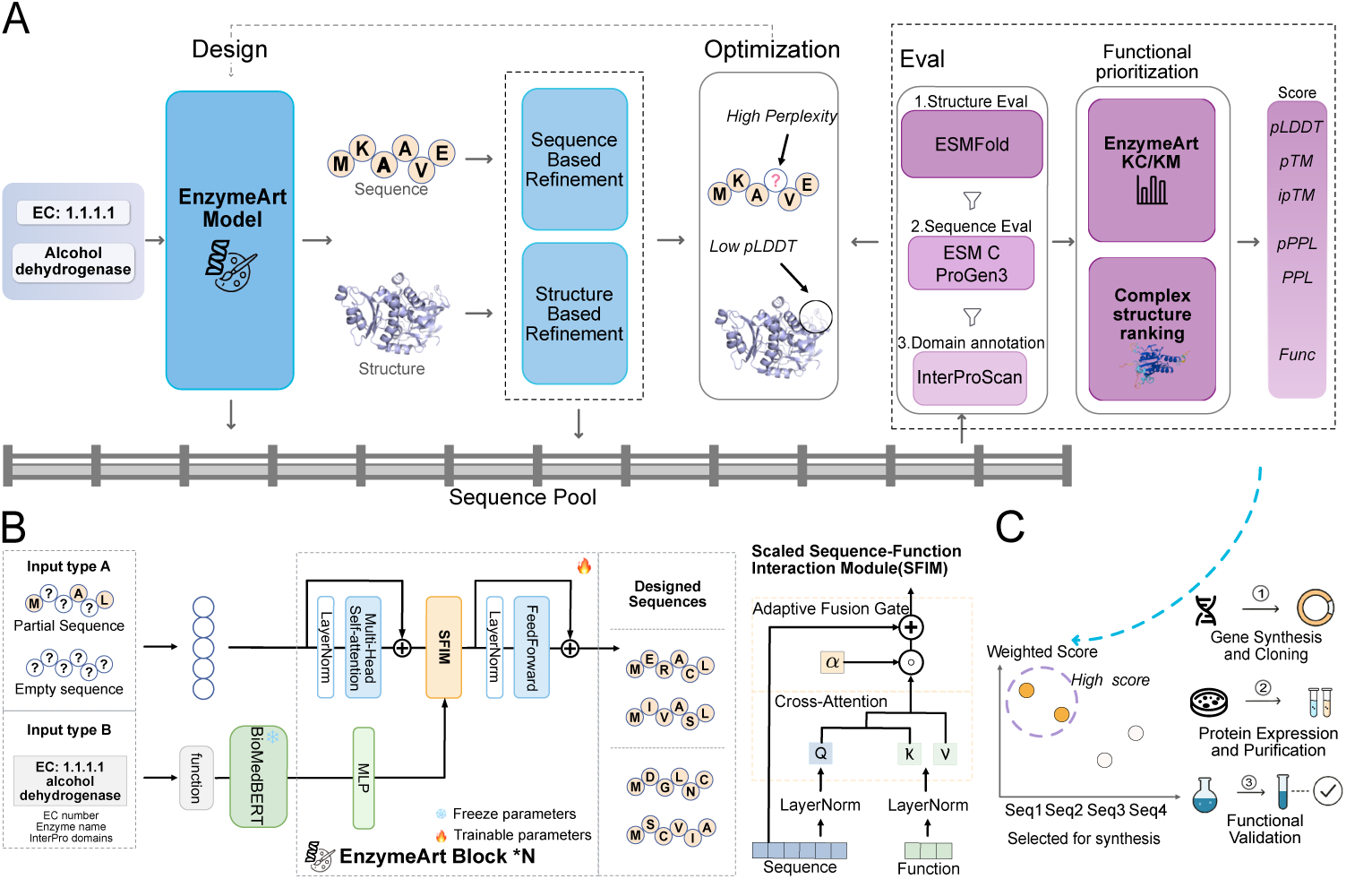
The EnzymeArt framework for function-guided enzyme design. **A**, The generative pipeline begins with the EnzymeArt model, which translates functional prompts (e.g., EC number and enzyme name) into diverse sequence proposals. These sequences are subsequently redesigned using sequence– and structure-based criteria. All sequences are passed through a computational pipeline for scoring based on kinetic prediction, predicted structure, sequence likelihood, and functional evaluation. **B**, Architecture of the EnzymeArt model, which integrates multiple input modalities to guide sequence generation. The SFIM module incorporates functional features. **C**, Top-ranked candidates from the task-adaptive prioritization pipeline are selected for experimental validation.

In the optimization stage, candidate sequences are refined through iterative structure– and sequence-based optimization (Methods 4.5.6). Predicted structures are used to evaluate structural confidence and provide structural inputs for ProteinMPNN inverse-folding redesign, whereas ESM C residue-level confidence identifies low-certainty sequence positions for targeted remasking and EnzymeArt-guided redesign. Low-confidence positions are selectively redesigned while higher-confidence regions are retained, improving predicted structural confidence and sequence plausibility over refinement cycles.

In the evaluation stage, refined sequences are ranked through an integrated assessment of catalytic potential, structural plausibility, sequence quality and functional consistency (Fig. 1C). EnzymeArt-KC provides substrate-conditioned, rank-normalized scores for candidate prioritization, with higher scores indicating higher predicted relative turnover. Additional structural, sequence and annotation-based analyses assess fold confidence, sequence plausibility, sequence novelty and functional-domain consistency. Together, these stages narrow large generated pools to prioritized enzyme designs for synthesis and experimental validation.

### 2.2 Computational assessment of EnzymeArt-generated enzyme candidates

We adopt a two-stage training strategy for enzyme generation to satisfy both structural requirements and functional constraints. In the first stage, EnzymeArt-base is pretrained on large collections of natural protein sequences, enabling the model to learn sequence patterns. We train EnzymeArt-base on a multi-modal dataset (Methods 4.3.1). In the second stage, we fine-tune the model on enzyme-focused datasets, allowing it to capture catalytic features and active-site motifs. We refer to this fine-tuned model as the EnzymeArt model, which we use as the conditional generator for all function-guided enzyme design experiments.

We evaluate the unconditional generation capability of EnzymeArt-base. In this task, the model generates complete protein sequences from masked inputs without any functional conditioning (Methods 4.5.2). We sample target sequence lengths from 100 to 1,000 amino acids (Methods 4.7.5). We use mean predicted LDDT (pLDDT) and predicted TM-score (pTM) as metrics to assess the structural quality of generated sequences. Across the entire length range, EnzymeArt-base generates sequences with high predicted structural confidence (pLDDT: 0.78 ± 0.04; pTM: 0.68 ± 0.08) (Fig. 2A). These values exceed those of recent generative models such as ESM3 and Pinal, indicating that EnzymeArt-base captures sequence features linked to protein structure. Maintaining coherent tertiary structures in long proteins requires the coordination of extensive long-range interactions [36, 37]. Across increasing sequence lengths (100–1,000 amino acids), most generative models exhibit a decline in predicted structural confidence, as reflected by lower pLDDT and pTM scores. In contrast, EnzymeArt-base maintains consistently higher confidence across this range and preserves global fold integrity even beyond 500 residues (pLDDT: 0.78; pTM: 0.63). These results indicate that EnzymeArt-base generates sequences with higher structure prediction confidence over a broad span of sequence lengths, providing a reliable structural starting point for downstream enzyme design.

**Fig. 2.**
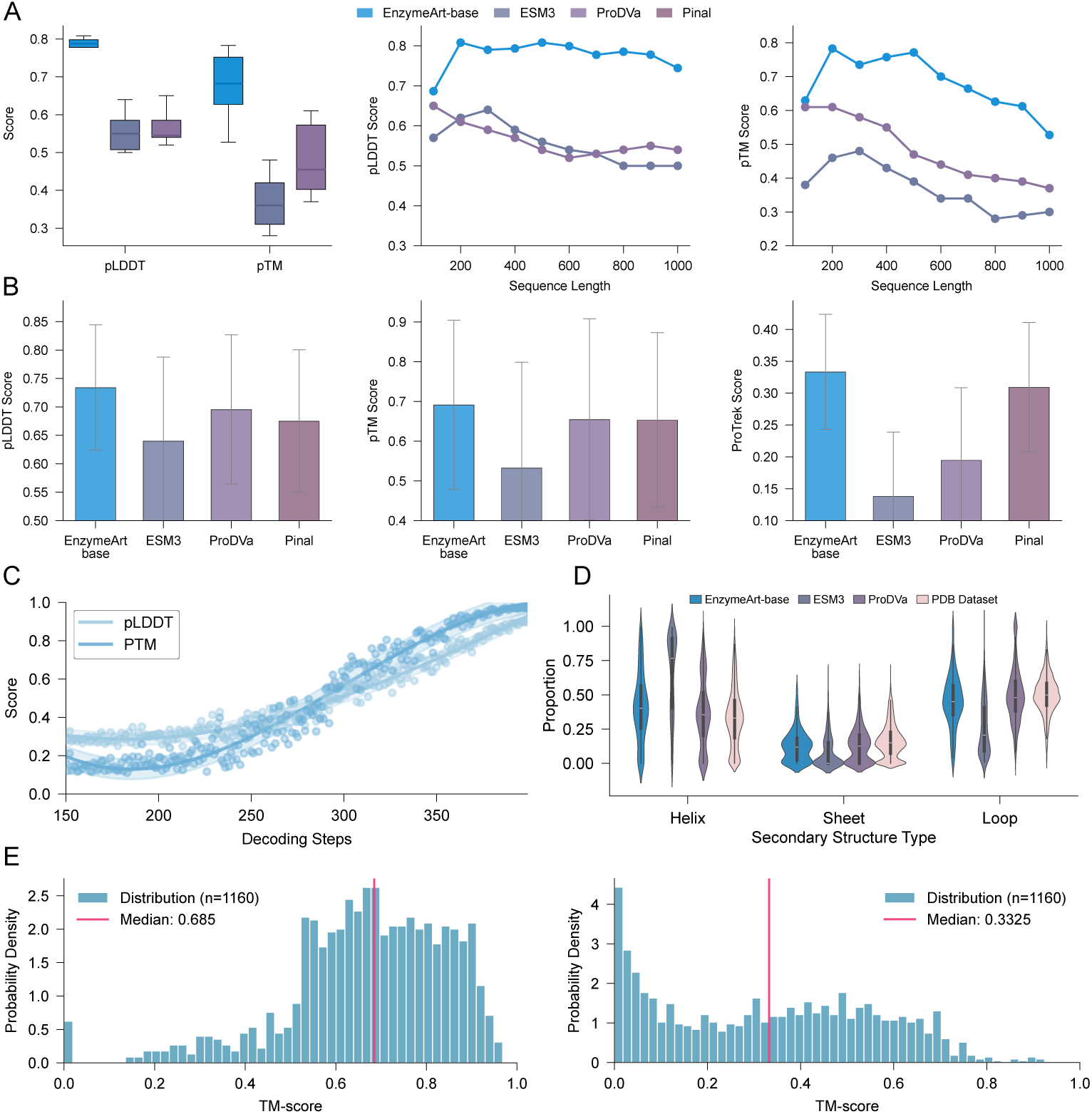
EnzymeArt-base is associated with higher predicted confidence in generative protein design. **A**, Unconditional generation. Predicted pLDDT and pTM scores for EnzymeArt-base are higher than those for ESM3, ProDVa, and Pinal across a range of sequence lengths. **B**, Conditional generation. In function-conditioned tasks, EnzymeArt-base attains higher values on the reported structural and functional metrics than the other methods considered. **C**, Generation trajectory. Both pLDDT and pTM scores progressively increase during the decoding process. **D**, Secondary Structure Distribution. The proportions of helices, sheets, and loops in the model’s designs closely recapitulate the distributions observed in natural proteins (PDB dataset), indicating high structural realism. **E**, Diversity and Novelty. TM-score analysis indicates that generated proteins are consistent with novel and diverse structures.

Next, we evaluate the function-conditioned generation capability of EnzymeArt-base. In this task, the model generates complete protein sequences given 1,000 functional prompts and fully masked sequence inputs (Methods 4.7.7). The resulting designs show high predicted structural confidence (pLDDT: 0.73 ± 0.11; pTM: 0.69 ± 0.21). We evaluate the alignment between each generated sequence and its intended catalytic role using the ProTrek score, which quantifies how well sequence features reflect functional descriptions (Methods 4.7.8). EnzymeArt-base achieves a mean ProTrek score of 0.33 ± 0.09, compared with 0.14 ± 0.10 for an ESM-3 baseline evaluated on the same benchmark (Fig. 2B). With increasing decoding steps, the model’s structural metrics can be further improved (Fig. 2C). The generated enzymes exhibit low sequence similarity among designs and low structural similarity to PDB references (mean pairwise TM-score: 0.23 ± 0.21; Methods 4.7.6), suggesting that the generated sequences exhibit diversity rather than simple duplication of reference structures (Fig. 2E).

We next examine the EnzymeArt model, which builds on EnzymeArt-base to generate enzymes directly from functional prompts (Methods 4.7.7). To assess generalization across enzyme classes, we evaluate subclass-wise structural confidence following the procedure in Methods 4.7.9. EnzymeArt produces consistently high-confidence structures in major enzyme families such as oxidoreductases (EC 1) and hydrolases (EC 3), and also maintains coherent folds in sparsely represented subclasses (EC 6 and EC 7) (Fig. 3A). In function-specified design tasks, EnzymeArt achieves high structural confidence and strong functional alignment (Fig. 3B), based on 1,000 randomly sampled functional prompts from the dataset. On average, EnzymeArt achieves pLDDT and pTM scores of 0.77 and 0.81, together with a ProTrek score of 0.28. Pinal produces structures with moderate predicted structural confidence (pLDDT: 0.74; pTM: 0.74) but shows lower functional correspondence (ProTrek 0.23). ESM3 exhibits similar patterns (pLDDT: 0.67; pTM: 0.59; ProTrek: 0.13). Fine-tuning ESM3 on an enzyme-specific dataset leads to a small increase in functional correspondence for ESM3-EC (ProTrek: 0.16), but with further reductions in pLDDT and pTM (0.63 and 0.51, respectively).

**Fig. 3.**
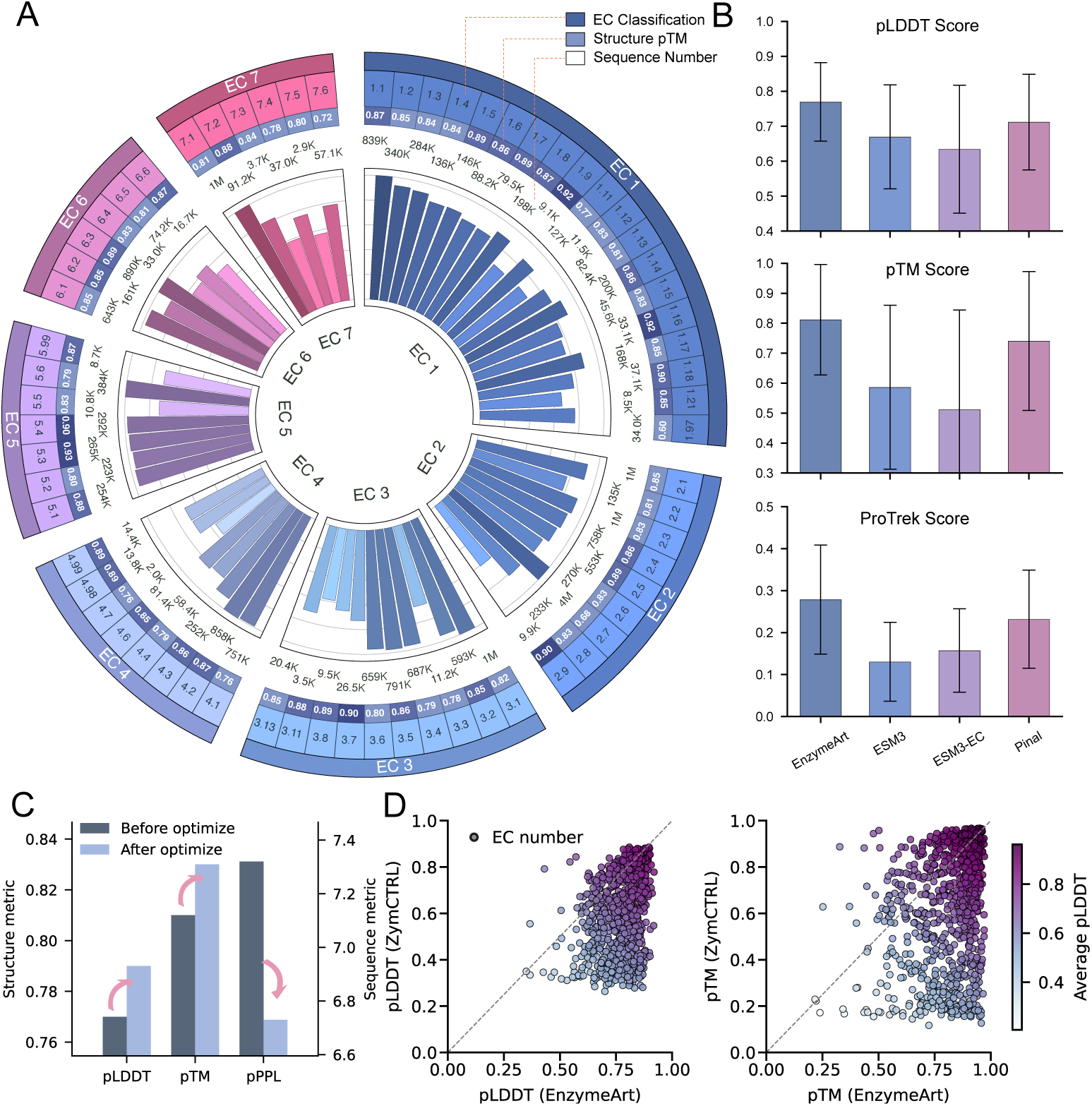
EnzymeArt model generates high-quality designs across the enzyme functional space. **A**, Performance overview showing consistently high average pTM scores for designs across major EC subclassifications. **B**, Benchmark comparison against other models indicates that, on this dataset, the EnzymeArt model yields higher predicted structural accuracy (pLDDT, pTM) and higher ProTrek scores. **C**, During the optimization process, ESM C and ESM-IF models further improve predicted structural and sequence plausibility. **D**, Across four-level EC specifications, EnzymeArt sustains high predicted structural confidence and shows higher pLDDT and pTM values than ZymCTRL on this benchmark, suggesting consistent performance across the enzyme classes evaluated.

The refinement stage modestly increases structure prediction confidence scores, with mean pLDDT changing from 0.77 to 0.79 and pTM from 0.81 to 0.83, while improving sequence plausibility, as reflected by a reduction in pPPL (pseudo-perplexity) from 7.32 to 6.73 (Fig. 3C).

Finally, we compare EnzymeArt with a specialized enzyme-generation model, ZymCTRL (Fig. 3D). To facilitate comparison, we evaluate both models across 897 enzyme classes in EC 1 and EC 2. EnzymeArt occupies higher-confidence regions in the pLDDT–pTM space, indicating that its generations form more confidently predicted structures across enzyme families. By contrast, ZymCTRL exhibits a broader spread with lower pTM values (Methods 4.7.10).

### 2.3 Catalytic-geometry controllability analysis

Catalytic activity arises from the precise spatial arrangement of active-site residues, a geometry that is highly conserved across divergent folds [38–40]. We therefore assess whether, given a functional description and a known catalytic site, EnzymeArt can generate backbones that preserve the required catalytic geometry. Structural confidence is measured by the pTM, and catalytic geometry by the root-mean-square deviation of active-site heavy atoms (cRMSD).

As a controlled motif-prompting test, we supply the ADH functional prompt together with a zinc-binding residue specification and evaluate whether the generated sequences retain the expected local catalytic geometry. The design recovers the overall architecture expected for ADHs and shows high global confidence (pTM: 0.96). At the catalytic site, the surrounding side-chain geometry closely matches a natural reference (cRMSD: 0.31 Å over heavy atoms in the annotated residues; Fig. 4A). The designs show Rosetta energy scores within a comparable range to natural scaffolds, providing an auxiliary modeled-energy assessment of conformational compatibility with the target active-site geometry (Fig. 4B; Methods 4.7.11). We do not constrain the backbone outside the zinc site, suggesting that a minimal functional or motif prompt in this controlled setting can steer sequence choices toward an ADH-compatible active-site environment (Methods 4.8.4).

**Fig. 4.**
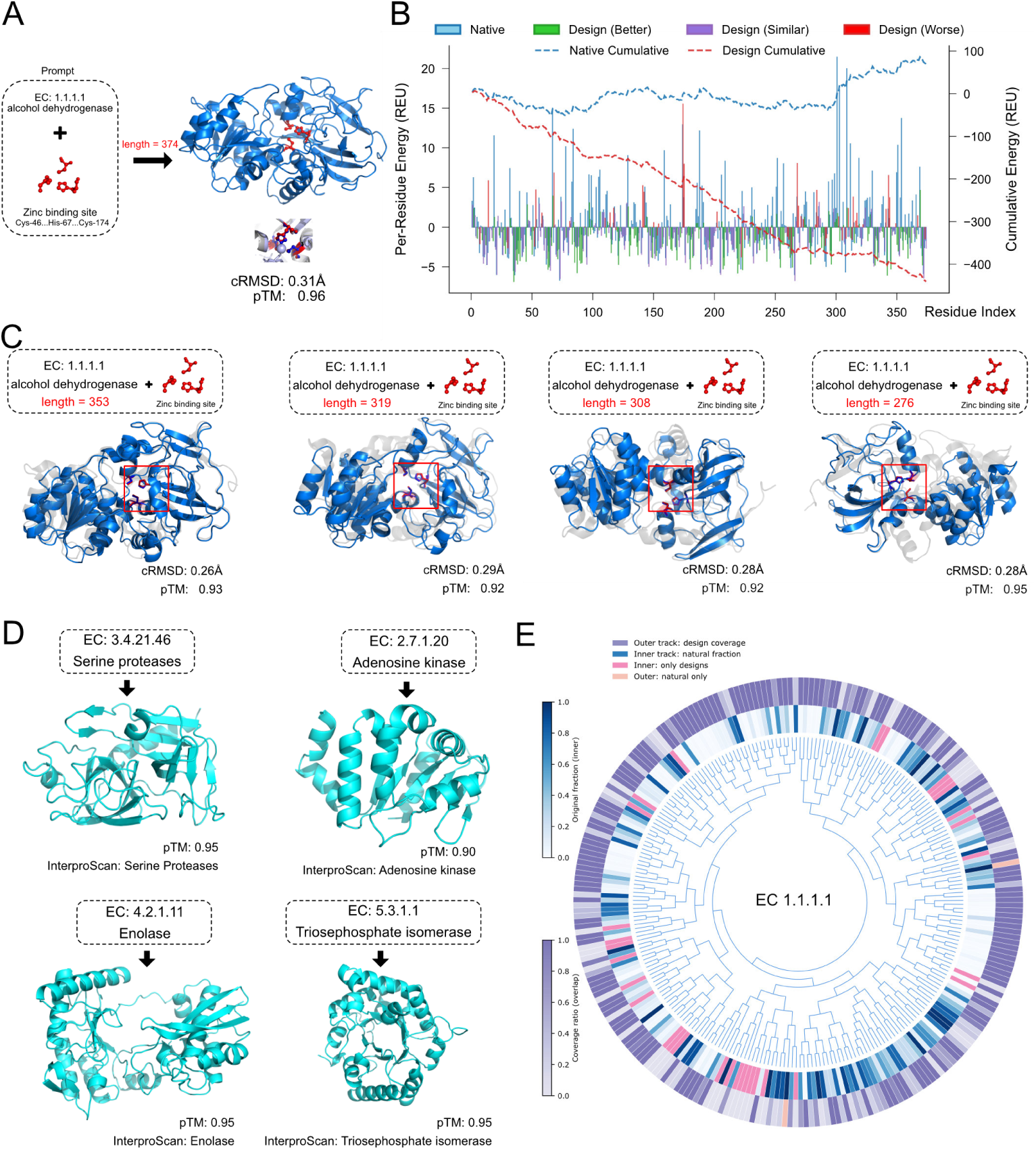
EnzymeArt generates enzymes with high predicted structural confidence, functional coherence, and evolutionary diversity. **A**, High-resolution design of an alcohol dehydrogenase (EC 1.1.1.1). The designed structure (blue) achieves active site accuracy (cRMSD = 0.31 Å on catalytic residues Cys46, His67, Cys171) and high global confidence (pTM = 0.96) compared with the native structure (grey). **B**, Rosetta energy analysis provides an auxiliary modeled conformational-energy comparison between the designed ADH and a natural reference structure. **C**, Generation of length-variable ADH designs (276–353 residues) while preserving the specified catalytic geometry. All variants maintain active site fidelity (cRMSD = 0.26–0.29 Å) and high overall structural confidence (pTM = 0.92–0.95). **D**, Representative designs for diverse EC classes, all exhibiting high pTM scores and correctly identified functional domains via InterProScan. **E**, Phylogeny-aligned circular representation illustrating that EnzymeArt-designed alcohol dehydrogenases (outer) recapitulate nearly all natural EC 1.1.1.1 clusters (99.1% coverage) and introduce 31 novel sequence clusters that expand beyond known natural diversity.

To evaluate how well EnzymeArt handles variable sequence lengths when the functional site is fixed, we prompt the EnzymeArt model to generate ADH designs spanning 276–353 amino acids under a fixed zinc-binding active-site specification. Across this range, the designs retain the same catalytic geometry (cRMSD: 0.26–0.29 Å) and high global confidence (pTM: 0.92–0.95) (Fig. 4C). This ability to accommodate length variation within a family suggests that the model can generalize across diverse realizations of a given enzyme family.

EnzymeArt extends beyond a single sequence family to generate enzymes from multiple classes. We prompt the model with specifications for serine protease (EC 3.4.21.46), adenosine kinase (EC 2.7.1.20), enolase (EC 4.2.1.11), and triosephosphate isomerase (EC 5.3.1.1) (Methods 4.7.9). In each case, the designs achieve high pTM values and recover the expected domains (Fig. 4D).

To assess how EnzymeArt explores and extends natural sequence space, we use alcohol dehydrogenases (ADHs; EC 1.1.1.1) as a representative family (Methods 4.8.3). We collect all UniProtKB entries annotated with EC 1.1.1.1 and remove sequences that lack explicit functional annotation. We embed natural and EnzymeArt-designed sequences using ESM2, project them using UMAP, and cluster them with k-means (k = 256) followed by hierarchical linkage to construct a circular sequence-space map (Fig. 4E). Natural sequences occupy 225 clusters, whereas EnzymeArt designs span 254 clusters, with 223 clusters shared. Thus, 99.1% of natural sequence groups are represented among EnzymeArt designs. EnzymeArt-designed sequences exclusively occupy 31 clusters, 20 of which show high predicted pTM score. The continuity between the inner (natural) and outer (designed) rings indicates that EnzymeArt reproduces the organization of natural sequences while extending into previously unoccupied regions of viable sequence space. These results demonstrate that the model maintains catalytic geometry while expanding the sequence space associated with alcohol dehydrogenase diversity.

Together, these results show that EnzymeArt preserves catalytic geometry, supports controlled diversity in sequence and fold, and extends the explored sequence manifold of enzymes.

### 2.4 EnzymeArt-KC Prioritizes Designs for Experimental Testing

EnzymeArt generates large candidate pools whose catalytic performance varies widely across sequences. To prioritize designs for experimental validation, we introduce EnzymeArt-KC/KM, a substrate-conditioned scoring model that estimates catalytic turnover and affinity and provides a quantitative basis for selecting top candidates (Methods 4.6.4).

The model is trained on 27,179 enzyme–substrate kinetic measurements spanning 12,707 enzymes, with 10% of the data randomly held out as a test set. Protein sequences are encoded with a Transformer, and substrates are represented by a text encoder. The sequence and substrate encoders produce embeddings that are merged into a joint representation. The joint representation is passed to two independent ensembles of multilayer perceptrons. Each ensemble aggregates the outputs of multiple predictors to produce a final estimate, with one ensemble predicting k_cat_ and the other predicting K_M_ (Fig. 5A). Because experimental k_cat_ values span several orders of magnitude [41], EnzymeArt-KC outputs rank-based normalized scores between 0 and 1 instead of raw estimates (Methods 4.6.2). For each substrate, the model aims to place sequences in the correct order of activity rather than to reproduce absolute kinetic values. This formulation matches the practical goal of screening, where only the relative ordering of candidates is needed for choosing variants to test in the laboratory.

**Fig. 5.**
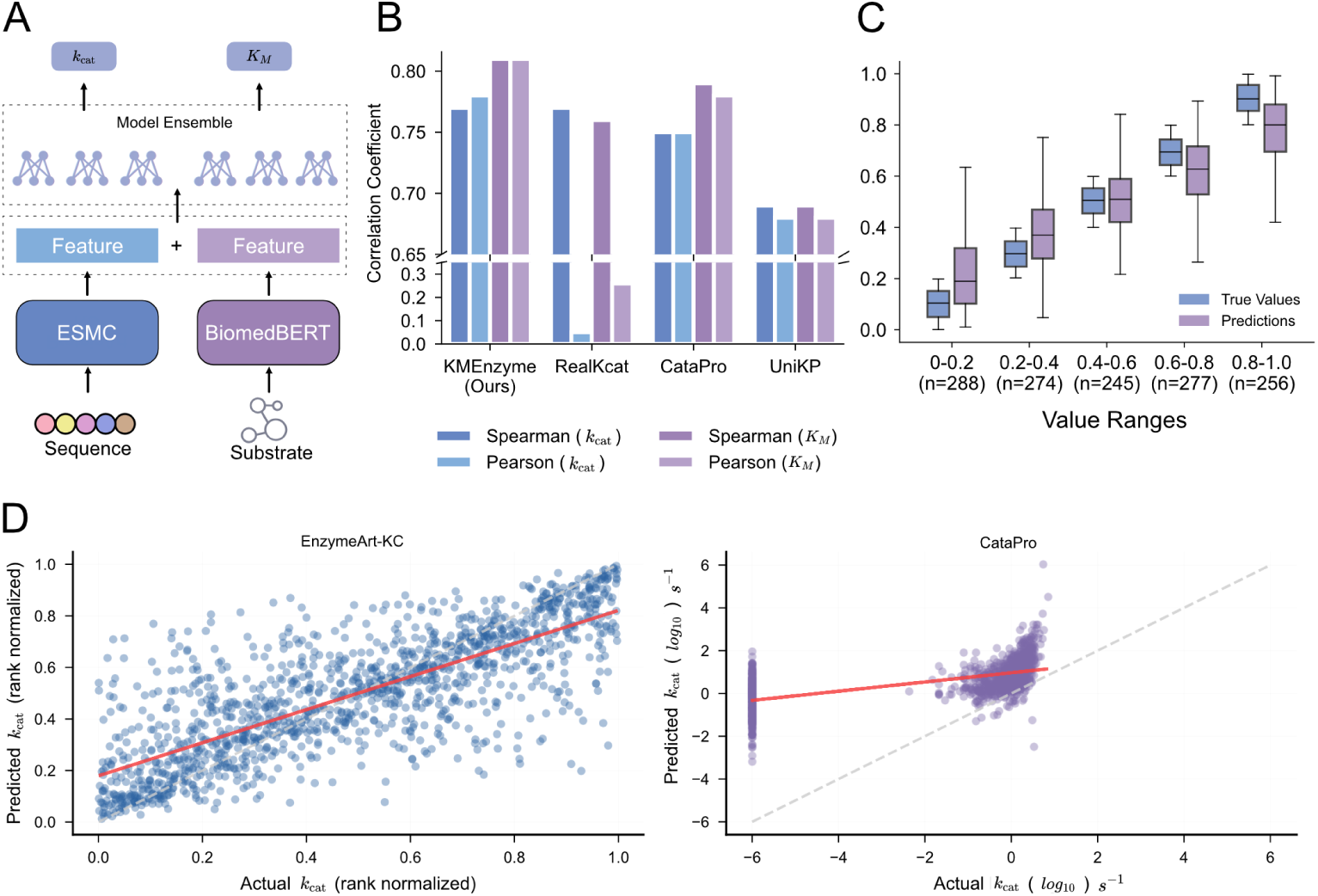
EnzymeArt predicts enzyme kinetic parameters using a rank-based strategy. **A**, The EnzymeArt-KC/KM model employs a dual-encoder architecture that jointly embeds protein sequences and substrates to predict both *k*_cat_ and *K*_M_. **B**, Benchmarking against the methods considered shows that EnzymeArt-KC/KM achieves higher Spearman and Pearson correlations for both *k*_cat_ and *K*_M_ on this dataset. **C**, Model predictions follow the true value trend across ranges, demonstrating interval discrimination. **D**, Comparison of prediction performance on rank-normalized *k*cat values (left) versus absolute log_10_-transformed values (right), illustrating the behavior of the rank-based approach.

On the held-out test set, EnzymeArt-KC/KM achieves higher correlations than baseline models in predicting both k_cat_ and K_M_ (Fig. 5B). For k_cat_, predicted and experimental values show a strong rank correlation (Spearman’s ρ = 0.77) and a high linear correlation on the transformed scale (Pearson’s r = 0.77). K_M_ shows a similar pattern, as higher affinity corresponds to lower K_M_ values. Predicted catalytic rankings closely mirror the distribution of true ranking (Fig. 5C). Within the highest k_cat_ region, the model exhibits a slight conservative bias, modestly underestimating the largest k_cat_ values. This behavior may bias downstream experimental choices toward more conservative candidates.

To focus on prioritization rather than absolute values, we evaluate predictions and targets within their original scales (Fig. 5D). Against a rank-normalized target, EnzymeArt-KC shows a strong monotonic relationship (Spearman’s ρ = 0.77) with a coefficient of determination of R^2^ = 0.60. In contrast, CataPro directly regresses log_10_(k_cat_). This log transformation compresses values across different kinetic regimes into a narrower numerical range, which may reduce the separability of activity differences and contribute to the broader spread of predictions on the log scale (R^2^ = 0.56). Because r and R^2^ depend on how we transform the response variable, we treat them as descriptive metrics. Under this comparison, EnzymeArt-KC provides more consistent prioritization for both k_cat_ and K_M_.

Within the EnzymeArt pipeline, EnzymeArt-KC scores every generated enzyme–substrate pair and filters candidates before structure-based evaluation. In practice, we retain only the top-ranked fraction of sequences for downstream modeling and experimental assays, which substantially reduces the search space while preserving high-activity designs. This selection allows structure prediction and analysis to focus on a small subset of kinetically promising candidates.

### 2.5 Functional prompting generates experimentally active enzymes

Computational prioritization identifies designs with favourable predicted properties, but catalytic activity must be established experimentally. We therefore used biochemical assays to test whether EnzymeArt could convert functional descriptions into active enzymes.

We selected three enzyme settings that differed in substrate type, reaction chemistry and assay readout. Alcohol dehydrogenase (ADH) provided a tractable but stringent redox design problem, in which generated candidates could be screened for ethanol oxidation. Malate dehydrogenase (MDH) tested whether the same strategy could produce malate-dependent turnover in a distinct dehydrogenase family. Triacylglycerol lipase extended the validation beyond nicotinamide-dependent redox chemistry to ester hydrolysis, with activity quantified by glycerol release from triglyceride substrates.

Across these three synthesis panels, we first performed repeated crude-lysate activity assays under the corresponding substrate condition: ethanol for ADH, malate for MDH, and triglyceride substrates for lipase. Each design was tested in three independent crude-lysate experiments with matched assay-background controls and empty-vector controls. A design was classified as active only when activity was reproducibly detected above the corresponding controls. Using this criterion, 57 of 60 tested designs showed measurable activity, corresponding to a 95% crude-lysate hit rate across ADH, MDH and lipase. We then used purified-enzyme kinetics to ask what forms of catalytic performance were recovered in each enzyme setting.

We began with ADH, a well-characterized but still demanding redox enzyme family. Ethanol oxidation requires cofactor-dependent hydride transfer together with productive positioning of the alcohol substrate. From a functional text prompt specifying ADH, EnzymeArt produced a diverse pool of candidate sequences (Fig. 6A). Substrate-aware kinetic prediction, together with structural, sequence and domain-level filters, prioritized 20 non-redundant designs for synthesis.

**Fig. 6.**
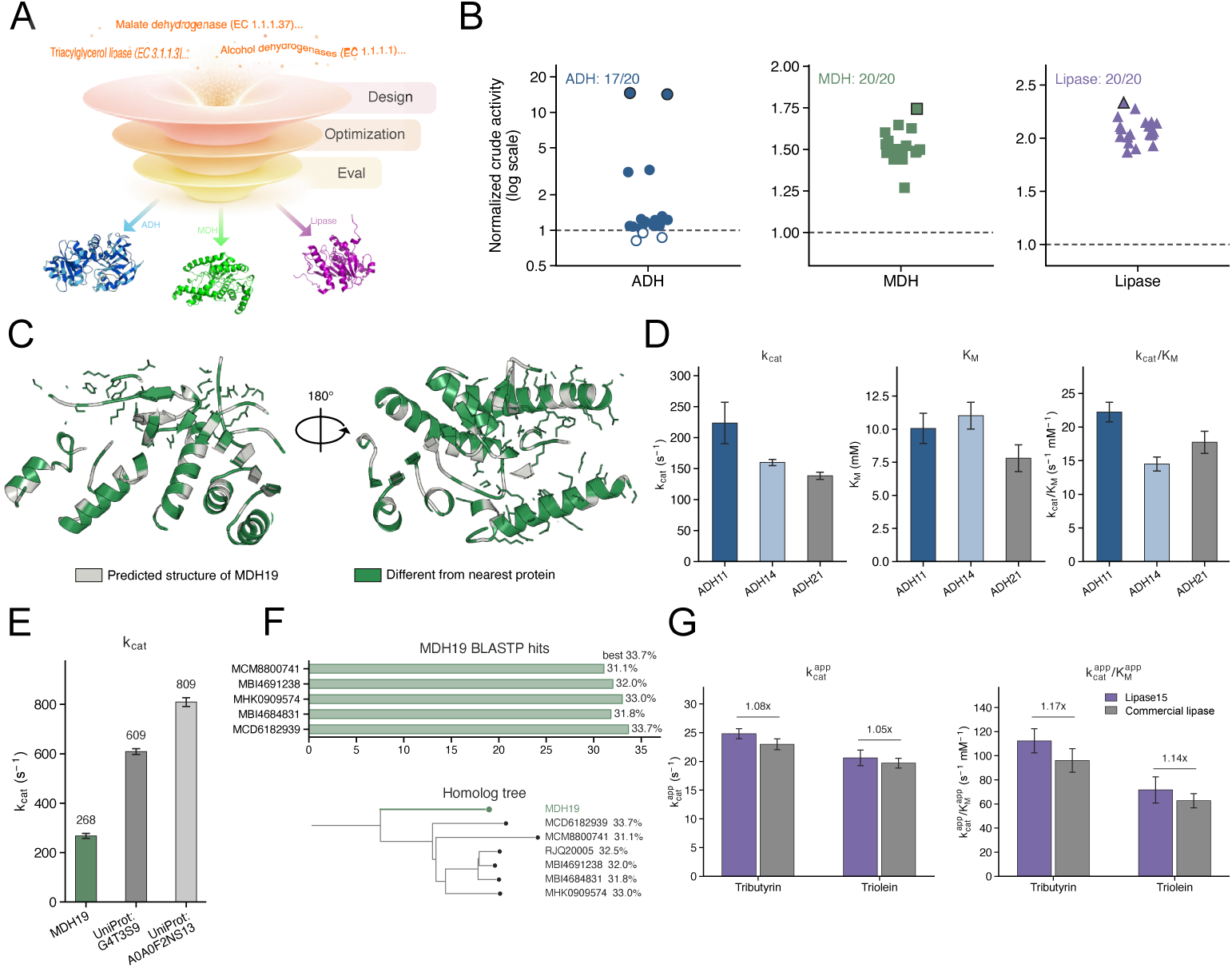
Experimental validation of designed enzymes. **A**, Workflow for function-guided enzyme generation, optimization and evaluation. **B**, Crude-lysate activity of ADH, MDH and lipase designs relative to matched controls. Each point represents the mean of three independent crude-lysate assays. Dashed lines indicate matched-control activity. Open symbols mark ADH designs not significantly above background. **C**, Two cutout views of the predicted structure of MDH19. Residues differing from its closest natural homolog are shown in green. **D**, Purified ADH kinetics for ADH11 and ADH14 compared with the natural reference ADH21/P00330 under matched ethanol-oxidation assay conditions. **E**, Purified MDH turnover for MDH19 and two natural MDH controls under matched malate-conversion assay conditions. **F**, BLASTP sequence-divergence analysis and homolog tree for MDH19. **G**, Purified lipase apparent kinetics for Lipase15 and a commercial lipase control on tributyrin and triolein. Bars show substrate-specific 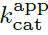 and 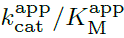.

In three independent crude-lysate assays, 17 of the 20 ADH designs showed significant ethanol-dependent activity relative to the empty-vector control (Fig. 6B). The strongest designs, including ADH11 and ADH14, consistently produced activity well above assay background and retained activity across multiple alcohol substrates (Supplementary Fig. 14). After purification, both designs showed ethanol-dependent saturation kinetics. ADH11 reached k_cat_ = 223.7 s^−1^ and k_cat_/K_M_ = 22.24 s^−1^mM^−1^, exceeding ADH21 from *Saccharomyces cerevisiae* (UniProt P00330) under the same assay conditions (Fig. 6D). ADH14 also retained substantial ethanol oxidation activity, with k_cat_ = 160.0 s^−1^ and k_cat_/K_M_ = 14.52 s^−1^mM^−1^. Together, these results show that functional prompting, followed by substrate-aware prioritization, enriched a compact synthesis set for active ADH designs and yielded a purified enzyme that outperformed a canonical natural benchmark under matched assay conditions.

We next asked whether this strategy could produce activity in a distinct dehydrogenase setting. MDH also uses nicotinamide-dependent redox chemistry, but acts on malate rather than alcohol substrates and requires a different substrate-recognition environment. From an MDH functional prompt, EnzymeArt generated candidates that were prioritized by generation quality, predicted structural confidence, domain-level consistency, malate-conditioned catalytic-property prediction and sequence novelty (Fig. 6A). In three independent crude-lysate assays, all 20 tested MDH designs showed reproducible malate-dependent activity above the corresponding controls (Fig. 6B).

Among these designs, MDH19 was selected for purified characterization because it combined reproducible crude-lysate activity with high sequence divergence. Across detected BLASTP hits, MDH19 had a maximum sequence identity of 33% (Fig. 6C, F). Purified MDH19 showed malate-dependent saturation kinetics fit by the Michaelis–Menten model. Despite this sequence divergence, MDH19 reached k_cat_ = 267.57 s^−1^, although its apparent K_M_ and catalytic efficiency remained below those of the natural controls (Fig. 6E, Table 8). The MDH experiment therefore tested a different aspect of the design problem. Here, the key result was not only panel-level activity, but the recovery of a sequence-divergent enzyme that retained high catalytic turnover after purification. MDH19 suggests that EnzymeArt can produce active designs beyond close sequence recapitulation of detected natural homologues.

We then moved away from nicotinamide-dependent redox chemistry. Triacylglycerol lipases hydrolyse ester bonds in hydrophobic triglyceride substrates and do not use a redox cofactor. This changed both the chemistry and the experimental readout, with activity quantified by glycerol formation rather than NAD(H)-linked absorbance. Prompting EnzymeArt with triacylglycerol lipase function produced candidates that were screened for triglyceride hydrolysis (Fig. 6A). In three independent crude-lysate assays, all 20 lipase designs showed reproducible triglyceride-hydrolysis activity above the corresponding controls. The designs hydrolysed tributyrin, a short-chain triglyceride, and also showed activity on triolein, a long-chain triglyceride. This cross-substrate activity indicates that the generated lipase panel supported triglyceride hydrolysis across substrates with different acyl-chain lengths.

Lipase15 was selected for purified characterization because it combined reproducible crude-lysate activity with favourable expression and sequence-divergence properties. Across detected BLASTP hits, Lipase15 had a maximum sequence identity of 58%. Purified Lipase15 hydrolysed both tributyrin and triolein. Under matched assay conditions, the designed lipase exhibited 1.17-fold and 1.14-fold higher apparent catalytic efficiency toward these substrates, respectively, compared with the commercial lipase control (Fig. 6G; Table 9). The lipase campaign therefore provided the strongest test of generality across reaction chemistry. The recovery of a purified design that hydrolysed both short– and long-chain triglycerides, with apparent activity modestly above an external commercial benchmark, shows that the workflow can extend from redox enzymes to triglyceride hydrolysis.

Taken together, these assays show that functional prompts can be converted into proteins with experimentally measurable catalytic activity, rather than only plausible annotations or predicted folds. The crude-lysate screens established panel-level enrichment across three enzyme settings, with 57 of 60 designs active across ADH, MDH and lipase assays. Purified-enzyme characterization then revealed three complementary forms of experimental success: an ADH design surpassing a canonical natural benchmark, an MDH design retaining substantial turnover despite low sequence identity to detected homologues, and a lipase design active on both short– and long-chain triglycerides. These results connect function-conditioned generation to measured catalytic activity across the tested redox and hydrolytic settings.

## 3 Discussion

Designing enzymes requires identifying sequences that not only fold into well-folded structures but also support precise active-site geometries. While modern generative models can now produce sequences that differ substantially from those explored by natural evolution, the central challenge lies in determining which among these sequences are viable. Natural evolution proceeds through incremental mutational steps that typically preserve function across intermediates, thereby constraining the regions of sequence space that remain accessible. Generative models, by contrast, can propose sequences that lie far outside these historically traversed paths, expanding the search space, but at the cost of introducing a vast fraction of sequences whose stability or activity is uncertain. Consequently, generation alone is insufficient: making enzyme design practical requires a framework for evaluating and selecting designs that are both structurally consistent and catalytically competent.

In EnzymeArt, we integrate functional conditioning, structure-aware refinement, and task-adaptive computational prioritization into a unified workflow. This framework does not restrict the generative capacity of the model, but instead prioritizes sequences that preserve catalytic-site geometry and exhibit favourable predicted turnover, enabling more reliable selection of experimentally testable candidates. This perspective raises a broader question about the organization of functional sequence space.

Generative enzyme design highlights a broader conceptual point: the space of sequences compatible with catalysis is likely far larger than the space evolution has realized. Evolutionary history reflects contingent paths, sequences that are discovered under specific ecological, energetic, and historical constraints, rather than the full extent of what is biophysically or catalytically possible. In this framing, natural enzymes represent one set of workable solutions, not the solution set itself. A generative model trained on evolutionary data learns the statistical structure of this solution manifold, including regions that evolution has approached only indirectly or not at all.

However, existence in this manifold does not guarantee accessibility. Many sequence realizations compatible with catalysis may lie in regions that evolution could not fea-sibly reach through stepwise mutations without transient loss of function. Generative models remove this continuity constraint; they can propose sequences that lie across evolutionary “gaps”. This capacity is powerful, but it also reveals the central challenge: expansion of the reachable landscape sharply increases uncertainty. As model capacity grows, the generative frontier moves outward into sequence regions where evolutionary data provide weaker guidance.

Viewed in this light, sequence generation must be coupled to principles of catalytic and structural coherence. In EnzymeArt, catalytic-site geometry functions as an organizing axis that persists across sequence divergence: although scaffolds may vary, the physical requirements of catalysis impose conserved spatial relationships among key residues. Similarly, kinetic rank scoring identifies sequences whose inferred energetic environments are compatible with catalytic turnover in a given substrate context. Rather than constraining the model’s creativity, these signals structure the expanded search space, distinguishing plausible designs from degenerate or non-functional ones.

This framework suggests a conceptual reframing of enzyme design. Instead of asking whether generative models can produce catalytically active sequences, a question now affirmatively answered, we must ask how to reliably identify the subset of generated sequences that inhabit the functional manifold. EnzymeArt represents one instantiation of such a framework: a model that explores broadly, coupled to biophysically grounded filters that select narrowly. The success of experimentally validated designs suggests that functional solutions are not arbitrarily distributed in sequence space, but can be enriched by structured constraints such as catalytic geometry, fold confidence, and kinetic ranking.

The added MDH and lipase validation cases are important because the original ADH task could, in principle, benefit from the dense representation of alcohol dehydrogenases in natural sequence databases. The MDH and lipase results help address this concern. In particular, MDH19 shares only 33% sequence identity with its nearest BLASTP hit, yet the purified enzyme shows malate-dependent Michaelis–Menten turnover. This provides an experimentally active lower-nearest-identity case, rather than simple recovery of a close natural homologue. Lipase15 further extends the validation to a hydrolase function and hydrolyses both short– and long-chain triglyceride substrates, with apparent activity comparable to or above the matched commercial lipase control under the same assay conditions. Together with the sequence-similarity analyses for the experimentally tested designs, these results help address the concern that the observed activity reflects only close homologue recovery within a densely represented ADH family.

At the same time, our results should be interpreted within the scope of the present study. The enzymes examined here belong to established natural enzyme classes, and experimental validation remains limited to a modest number of candidates per task. We therefore view EnzymeArt not as a demonstration of unrestricted enzyme design, but as evidence that function-guided generation, when coupled to structure-and kinetics-aware prioritization, can identify active and sequence-diverse candidates within known catalytic families.

More generally, these results support the view that functional enzyme sequence space may be broader and more navigable than what is directly accessible through stepwise evolutionary trajectories alone. Generative models expand what can be proposed, whereas coherent selection frameworks help determine what can be realized experimentally. Together, they outline a path toward more intent-driven discovery in protein design.

## 4 Methods

### 4.1 Overview of the EnzymeArt workflow

EnzymeArt is organized as a staged workflow for converting functional specifications into experimentally testable enzyme sequences. First, EnzymeArt-base is pretrained on a broad protein and function corpus to learn transferable sequence and functional priors. Second, the model is fine-tuned on enzyme records to obtain EnzymeArt, which generates sequences from functional descriptions such as enzyme names, EC numbers and InterPro domain terms. Third, generated candidates are refined and prioritized using structure prediction, sequence plausibility metrics, functional domain annotation and, where applicable, substrate-conditioned kinetic prediction.

The Methods section follows this workflow. We first describe the model architecture, training data and optimization procedure. We then describe function-conditioned generation, sequence refinement, kinetic prediction and computational evaluation. Finally, we give the design campaigns and experimental validation protocols for alcohol dehydrogenase (ADH), malate dehydrogenase (MDH) and triacylglycerol lipase.

### 4.2 Model architecture

#### 4.2.1 Tokenization and functional representations

*Protein sequence representation.* Protein sequences are tokenized using a fixed vocabulary of 33 tokens, comprising the 20 canonical amino acids, four non-standard residues (B, U, Z, O), and special symbols including <cls>, <pad>, <mask>, <unk>, and <eos>. Beginning-of-sequence and end-of-sequence tokens are prepended and appended during preprocessing.

*Functional descriptions.* Functional descriptions are tokenized using a WordPiece-style subword vocabulary of 28,895 tokens. This vocabulary covers alphanumeric characters, punctuation, multilingual units, and domain-relevant subwords (e.g., “enzym”, “substr”, “activ”). Both EC numbers and enzyme names are represented as sequences of these subword tokens, allowing consistent handling of structured identifiers and free-text descriptions within a unified functional embedding space.

#### 4.2.2 Model inputs and forward computation

As described above, EnzymeArt models accept multiple input tracks, each of which can be independently enabled or masked. For clarity, we denote the inputs to the model as

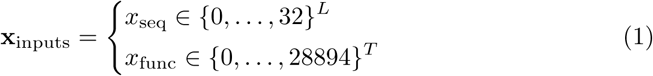

where x_seq_ denotes the protein sequence tokens drawn from a 33-token vocabulary, and x_func_ denotes the functional description tokens (EC numbers or enzyme names) drawn from a 28,895-token subword vocabulary.

For EnzymeArt-base and EnzymeArt, the protein sequence and functional description are processed by two separate encoders. The protein tokens are first mapped to a continuous representation using a SequenceEncoder, which provides contex-tualized embeddings of the residue sequence. In parallel, the functional description is encoded using a FunctionEncoder, and the resulting embedding is projected through a linear transformation to match the hidden dimensionality of the model.

The sequence embedding serves as the initial model state and is propagated through a stack of Transformer layers. Each layer first applies self-attention to capture dependencies between residues. The resulting sequence representation is then updated by the Sequence—Function Interaction Module (SFIM), which incorporates functional information by attending to the projected functional embedding. A feed-forward network further updates the combined representation before it is passed to the next layer.

After all layers are applied, the final hidden representation is decoded by a language modeling head, which produces logits for each residue position and defines the masked language modeling objective. This computation defines the forward pass used for both EnzymeArt-base pretraining and EnzymeArt sequence generation from functional descriptions.

**Algorithm 1.**
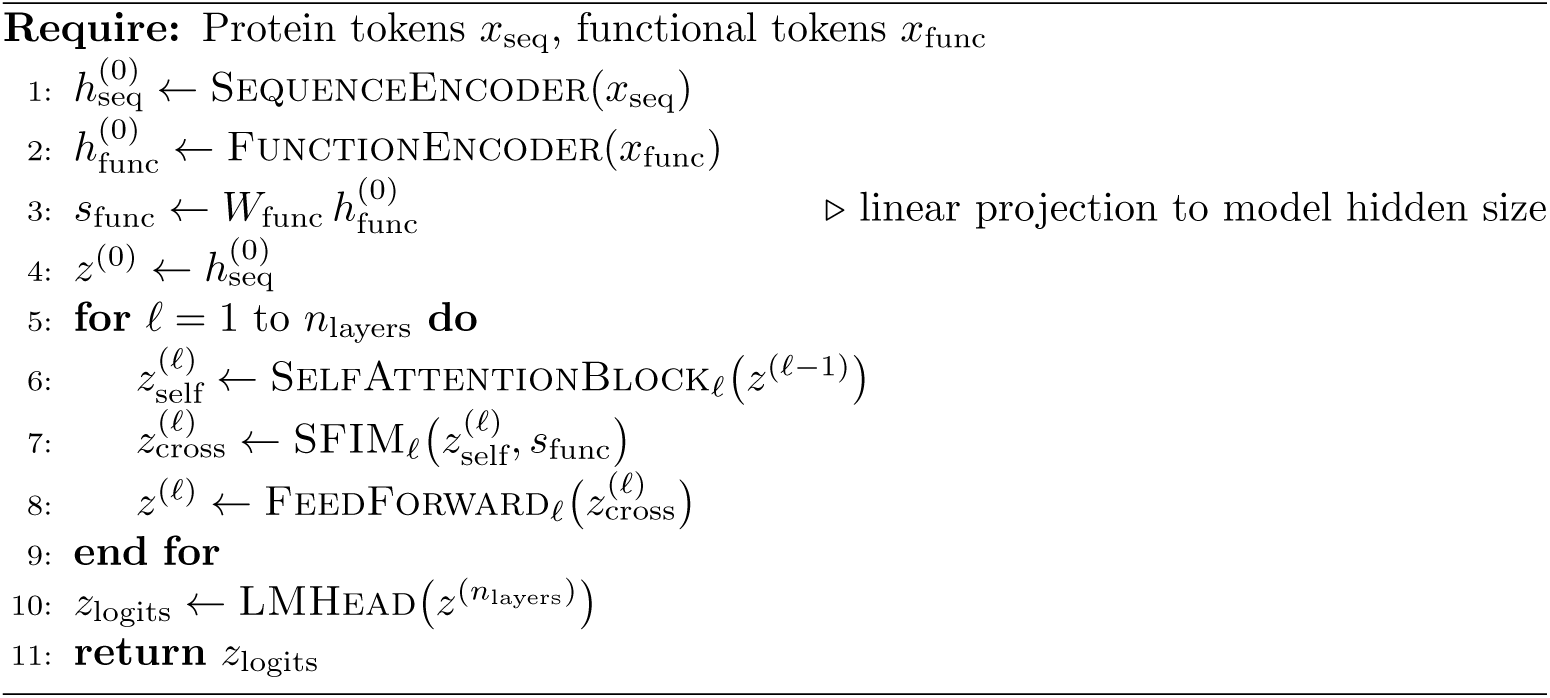
enzymeart_base_forward.

#### 4.2.3 Scaled sequence–function interaction module (SFIM)

To incorporate functional context throughout the network depth, EnzymeArt models employ a Scaled Sequence–Function Interaction Module (SFIM) in every Transformer layer. SFIM operates on the intermediate sequence representation and a shared functional embedding derived from the function encoder. At layer ℓ, the sequence state 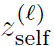 is first normalized, as is the projected functional embedding s_func_. SFIM then performs multi-head cross-attention, using the normalized sequence states as queries and the normalized functional embedding as keys and values. The resulting attention output captures position-specific interactions between the sequence and the functional description. This signal is modulated by a learned scalar gate and added back to the sequence representation through a residual connection, yielding an updated state conditioned on the intended catalytic function. Repeating this operation across layers enables the functional description to guide residue-level features in a distributed and hierarchical manner.

The learned scalar gate α*_ℓ_* stabilizes training by controlling the strength of functional conditioning, preventing early layers from being dominated by functional context while allowing deeper layers to integrate high-level semantics.

**Algorithm 2.**
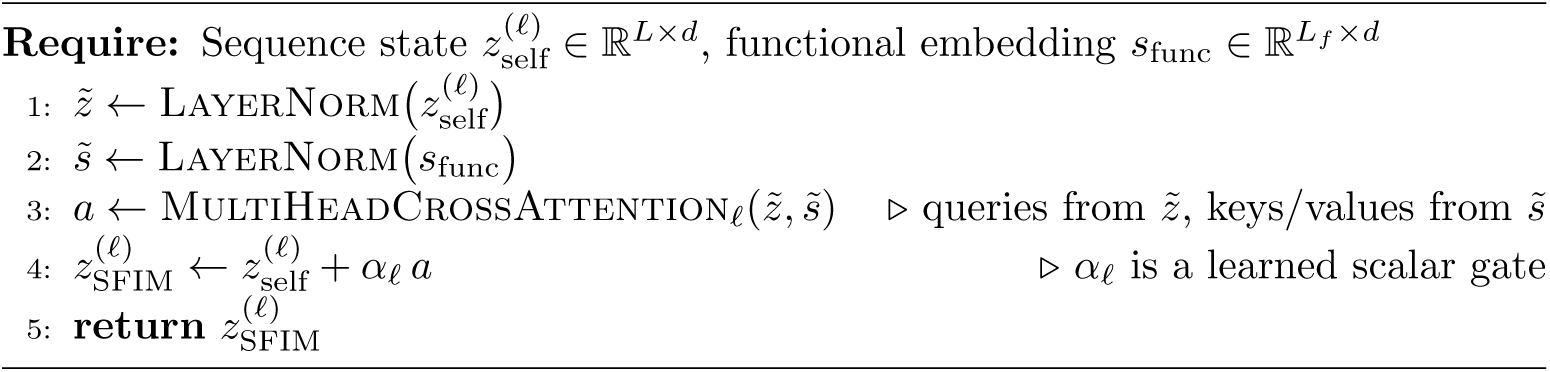
SFIM layer*_ℓ_*.

#### 4.2.4 Foundation model for protein and enzyme sequence generation

EnzymeArt-base is a general-purpose foundation model trained to learn how protein sequences relate to biological functions in full-length proteins. EnzymeArt-base and the enzyme-specialized EnzymeArt model use the same Transformer architecture, tokenization scheme and training objective, and differ only in their training data and optimization schedule.

The model aims to capture the organizing principles of protein evolution and biochemical function within a unified probabilistic framework. Unlike protein language models trained on datasets containing fragmented genomic sequences [42], or structure-first diffusion models such as RFdiffusion and Chroma [10, 11], EnzymeArt-base learns directly from full-length proteins with functional annotations. This enables the model to infer both local residue dependencies and global biochemical intent, supporting coherent generation of functionally meaningful proteins. EnzymeArt uses the same architecture and is derived from EnzymeArt-base by fine-tuning on a dataset specifically curated for enzyme design (Section 4.4.3).

**Table 1.** Training configurations of EnzymeArt-base models.

| Params | Layer | $d_{\text{model}}$ | $d_{\text{head}}$ | Context length | Learning rate | Warm-up steps |
| --- | --- | --- | --- | --- | --- | --- |
| 332M | 30 | 640 | 20 | 1024 | $4 \times 10^{-4}$ | 2000 |
| 1B | 33 | 1280 | 20 | 1024 | $4 \times 10^{-4}$ | 2000 |

**Table 2.** Training compute and token budget for EnzymeArt-base pretraining.

| Params | Batch size (tokens) | Num steps | Total tokens | FLOPs |
| --- | --- | --- | --- | --- |
| 332M | $1 \times 10^6$ | $1.3 \times 10^6$ | $1.3 \times 10^{12}$ | $2.6 \times 10^{21}$ |
| 1B | $1 \times 10^6$ | $5.0 \times 10^5$ | $5.0 \times 10^{11}$ | $3.0 \times 10^{21}$ |

### 4.3 Training data for pretraining and fine-tuning

#### 4.3.1 EnzymeArt-base-MultiModal dataset

The foundation model is trained on the EnzymeArt-base-MultiModal dataset, a large-scale corpus integrating protein sequences with functional domain annotations. Protein sequences are obtained from AFDB50 [43], a version of the AlphaFold Protein Structure Database clustered at 50% sequence identity [44], ensuring broad sequence diversity while minimizing redundancy.

Functional annotations are obtained using the UniProtKB mapping file protein2ipr.dat.gz (available from the InterPro FTP site). This file provides residue-level associations between UniProt accessions and InterPro domain signatures. Mapping is performed by joining AFDB50 sequences to their corresponding UniProtKB identifiers [45].

To maintain both diversity and functional interpretability, the dataset is divided into two subsets: one containing proteins with curated functional annotations (“With Function”) and another containing unannotated proteins (“Without Function”). The annotated subset provides explicit supervision for learning sequence–function relationships, while the unannotated portion allows the model to generalize across unlabeled structural patterns.

#### 4.3.2 EnzymeArt-MultiModal dataset

To specialize the foundation model for enzyme design, we construct EnzymeArt-MultiModal, a curated corpus of ∼24 million enzyme sequences with functional annotations. Protein sequences are sourced from UniProtKB by selecting all entries with assigned EC numbers (EC:*), ensuring that the dataset contains proteins with experimentally verified or computationally inferred enzymatic activities.

InterPro annotations are obtained from InterPro release 104.0 [46] using the UniProtKB mapping file protein2ipr.dat.gz, which provides residue-level links between UniProt accessions and InterPro domain signatures. Each sequence is assigned domain descriptions through this mapping. Entries lacking InterPro annotations or showing inconsistent or ambiguous domain assignments are excluded, yielding a high-confidence set of enzyme sequences.

Each record in EnzymeArt-MultiModal includes the amino-acid sequence, EC number, enzyme name, and InterPro-derived domain descriptions. The resulting distribution spans the full enzymatic hierarchy and provides broad functional coverage for fine-tuning EnzymeArt-base into an enzyme-specialized generative model.

**Table 3.** Dataset composition of EnzymeArt-base-MultiModal and EnzymeArt-MultiModal.

|  | With Function | Without Function | Total |
| --- | --- | --- | --- |
| EnzymeArt-base-MultiModal | 29,763,593 | 23,902,267 | 53,665,860 |
| EnzymeArt-MultiModal | 24,778,979 | 0 | 24,778,979 |

This dataset hierarchy provides a natural progression from general protein modeling to enzyme specialization: EnzymeArt-base learns universal protein grammar from EnzymeArt-base-MultiModal, while EnzymeArt inherits this knowledge and adapts it to enzyme-specific data from EnzymeArt-MultiModal.

### 4.4 Training objective and optimization

#### 4.4.1 Masked language modeling objective

Both EnzymeArt-base and EnzymeArt are trained using a generative masked language modeling (MLM) objective to capture long-range dependencies and bidirectional residue interactions. The MLM objective allows the model to learn bidirectional dependencies without the need for explicit structure supervision, which is critical given the scale and diversity of the dataset.

For a protein sequence x with masked positions m, we optimize the weighted negative log-likelihood of recovering the correct amino acids:

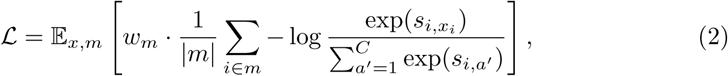

where s*_i,a_′* denotes the logits over the amino acid vocabulary C. The masking-ratio-dependent weight w*_m_* is defined as:

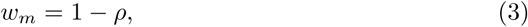

with ρ denoting the masking ratio. This weighting emphasizes sequences with richer contextual signal (lower ρ), while down-weighting heavily occluded examples.

Optimization employs the AdamW optimizer (β_1_ = 0.9, β_2_ = 0.98) with a two-phase learning-rate schedule: the learning rate is linearly increased to 4 × 10^−4^ during the first 2,000 warm-up steps and then decays to 4 × 10^−5^ following a cosine schedule over the subsequent training steps.

#### 4.4.2 Pretraining EnzymeArt-base

EnzymeArt-base is pretrained for 100,000 steps with masking ratios uniformly sampled from [0, 1], exposing the model to a wide spectrum of contextual completeness. Functional descriptions are stochastically dropped in 25% of samples, encouraging the model to integrate both intrinsic sequence constraints and auxiliary annotations without over-reliance on either.

An additional 100,000-step pretraining phase increases the functional dropout rate to 75%, encouraging the model to recover functional associations from minimal conditioning cues. This is intended to enhance robustness to sparse or incomplete annotations and may improve generalization to proteins outside the training distribution.

Increasing functional dropout from 25% to 75% encourages the model to rely more strongly on intrinsic sequence constraints while still incorporating functional context when available. Together, these training stages establish a representational backbone that jointly encodes residue-level dependencies and high-level functional semantics, supporting function-conditioned sequence generation.

#### 4.4.3 Fine-tuning EnzymeArt for functional specialization

To obtain the enzyme-specialized EnzymeArt model, we fine-tune EnzymeArt-base on the EnzymeArt-MultiModal dataset described above using the same MLM objective and SFIM-based architecture.

We fine-tune the model using the AdamW optimizer (β_1_ = 0.9, β_2_ = 0.98) with mixed-precision training. The learning rate is linearly increased from 0 to 1 × 10^−4^ during the first 2,000 warm-up steps, then decayed following a cosine schedule to 1 × 10^−5^ by step 500,000, after which it is held constant. Mini-batches contain 10,000 tokens per GPU, and gradient accumulation is applied to achieve an effective batch size of approximately 1 × 10^6^ tokens.

Conditioning inputs include amino-acid sequences together with multi-modal functional descriptions, which are encoded through separate embedding channels. These representations are fused by the SFIM module, allowing functional context to modulate token selection throughout the generation process. This setup enables the model to learn consistent mappings between functional descriptions and plausible enzyme sequences while preserving the evolutionary and fold-level organization captured during EnzymeArt-base pretraining.

During fine-tuning, all parameters of the Transformer backbone, embedding layers, and SFIM modules are updated jointly, without freezing any components.

#### 4.4.4 Implementation details

Training is performed in PyTorch on 8*NVIDIA A800 GPUs using BF16 mixed-precision computation. Sequences exceeding the maximum input length are cropped to 1,024 residues per sample, while shorter sequences are retained in full. We use the AdamW optimizer with (β_1_ = 0.9, β_2_ = 0.98, weight decay = 0.01) and cosine learning-rate decay with warm-up, as described above. Gradient accumulation is applied to achieve an effective batch size of approximately 10^6^ tokens. All experiments are run with fixed random seeds and synchronized data loaders to ensure reproducibility.

### 4.5 Sequence generation and refinement

EnzymeArt generates protein sequences using an iterative masked-refinement framework in which an initially uninformative sequence is progressively transformed into a coherent amino-acid chain [47]. EnzymeArt-Design extends this framework by conditioning on both evolutionary context and functional intent, enabling the model to synthesize amino-acid sequences consistent with a specified catalytic role. The model operates on an input pair (x, f), where x is a fully or partially masked protein sequence and f is a functional description such as an enzyme name or EC number.

#### 4.5.1 Iterative masked-refinement framework

Generation begins from a fully masked template where all residue positions are set to [MASK]. At each refinement cycle t, the model predicts amino-acid distributions conditioned on the current partial sequence x^(*t*)^ and the functional description f. Let ℓ*_i_*^(*t*)^(a) denote the pre-softmax logit for amino acid a at position i and refinement cycle t. These logits are converted to a categorical distribution

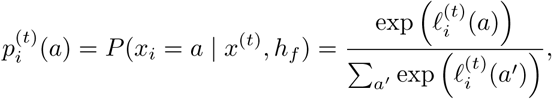

where h*_f_* is the functional embedding and x^(*t*)^ is the partially masked sequence at cycle t. Residue-level confidence is estimated as

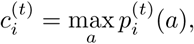

and provides a position-wise measure of how strongly the model prefers a single amino acid at that site.

A reparameterized update rule [48] is then applied to determine which positions are revised: low-confidence positions are re-masked and resampled, while higher-confidence residues are retained. This confidence-driven masking-and-refinement schedule enables globally coherent sequence formation by allowing high-confidence positions to consolidate while uncertain regions continue to evolve, avoiding premature fixation of suboptimal residue assignments. The refinement process proceeds for a fixed number of cycles, producing a complete amino-acid sequence without requiring structural templates or post-hoc filtering.

#### 4.5.2 Function-conditioned and unconditional generation

For function-conditioned generation, natural language prompts such as enzyme names or EC numbers are encoded using a biomedical language model and mapped to a continuous representation h*_f_* by a dedicated function encoder (Methods 4.2.3). SFIM injects this functional information into the generative backbone by modulating the logits ℓ*_i_*^(*t*)^(a) as a function of both the sequence context and h*_f_*, thereby shaping the distributions p*_i_*^(*t*)^(a) and the resulting refinement trajectory.

For unconditional sequence generation, we apply the same iterative refinement framework while withholding explicit functional information. The functional prompt is replaced with an [UNK] token and the input sequence remains fully masked. Across T = 500 refinement steps, editable positions are resampled using the same Gumbel–Softmax scheme and confidence-based masking policy, with the mode-collapse safeguard maintained. This configuration allows the model’s intrinsic priors over protein sequence space to be interrogated, enabling broad exploration of foldable and biophysically coherent sequences without requiring a template sequence.

**Algorithm 3.**
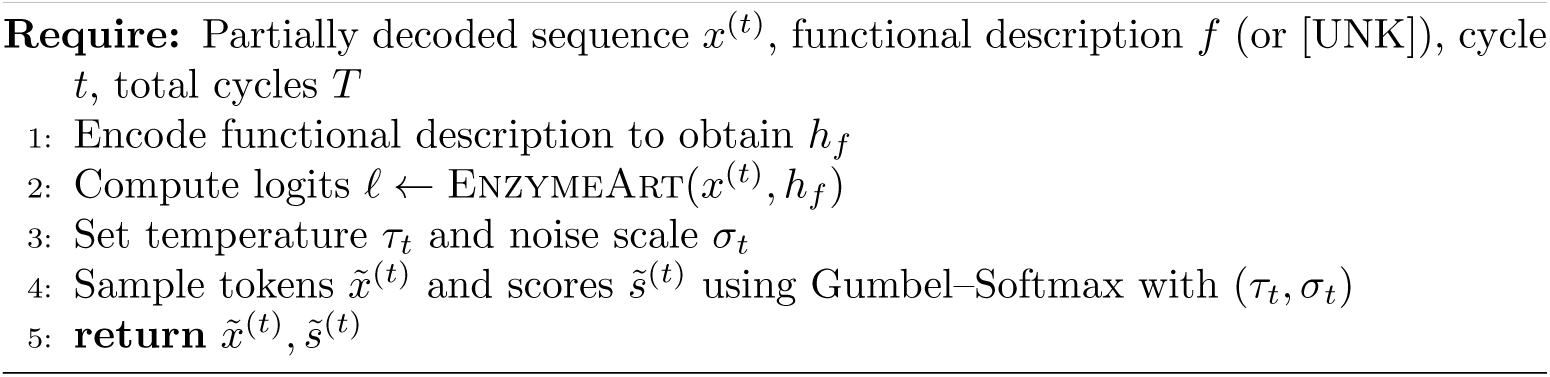
Token sampling at refinement cycle t.

#### 4.5.3 Refinement schedule

Unless otherwise stated, we perform iterative refinement for T = 500 cycles. At refinement cycle t, the fraction of positions eligible for re-masking is scheduled according to

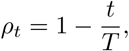

such that the lowest-confidence ⌊ρ*_t_* · L⌋ positions (where L is the sequence length) are re-masked and resampled, while higher-confidence positions remain fixed. Early cycles therefore allow broad exploration of sequence space, whereas later cycles perform increasingly conservative local adjustments.

For ADH design experiments, we draw resampled amino acids directly from the model’s categorical distribution, with special tokens disallowed. No nucleus filtering or additional diversity penalties are applied during the main sampling procedure; topp filtering is only invoked within the mode-collapse safeguard described below. No external structure prediction or functional annotation tools are used during generation. As such, the resulting sequences reflect the model’s internal statistical and functional priors, and sequence diversity arises intrinsically from the confidence-driven iterative refinement.

#### 4.5.4 Gumbel–softmax sampling for controlled diversity

To maintain stochasticity while preserving functional conditioning, editable positions are sampled using a Gumbel–Softmax perturbation of the categorical distribution:

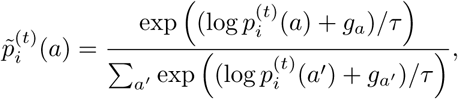

where g*_a_* ∼ Gumbel(0, 1) are i.i.d. noise samples and τ is a temperature parameter (optionally scheduled as τ*_t_* across refinement cycles). Amino-acid identities are then drawn from 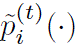 at each editable position. This reparameterization encourages exploration while retaining sensitivity to function-conditioned preferences encoded in p*_i_*^(*t*)^.

#### 4.5.5 Mode collapse prevention

To avoid trivial convergence toward homogeneous residue compositions, a simple safeguard is applied at each refinement cycle. If any amino-acid identity occupies more than a threshold fraction ρ = 0.25 of positions in the current sequence, all positions assigned to that residue are temporarily re-sampled once in that cycle using the same Gumbel–Softmax procedure (optionally with a mild top-p filter). Subsequent cycles then revert to the standard confidence-based masking and refinement policy. This mechanism maintains sequence-level diversity and reduces the risk of degenerate local optima induced by strong functional prompts [47], while still allowing the model to consolidate confident, functionally relevant motifs.

**Algorithm 4.**
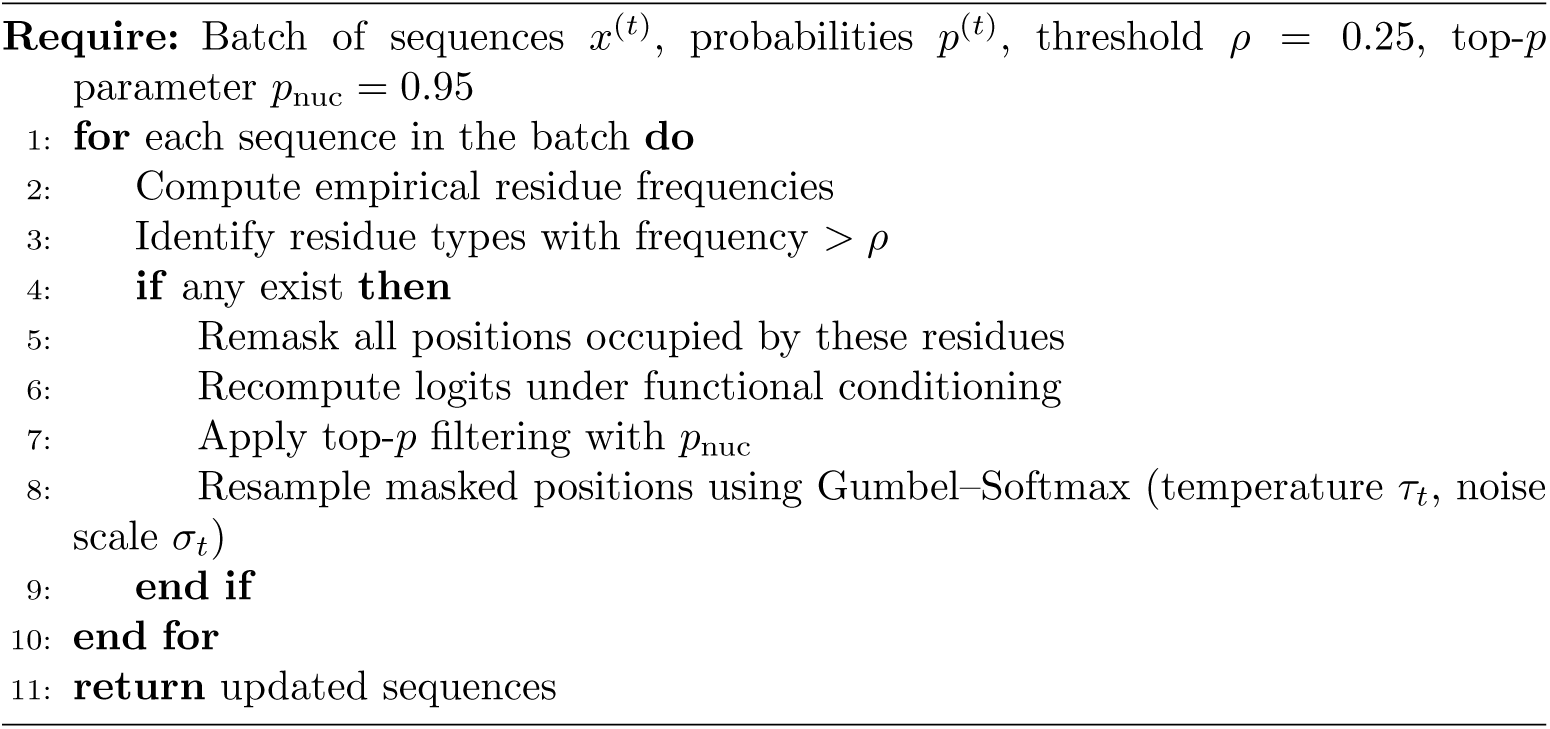
Conditional resampling for repetitive residues.

#### 4.5.6 Structure– and sequence-based optimization (EnzymeArt-Optimization)

To enhance the structural plausibility and sequence quality of designs generated by EnzymeArt, we employ a dual-stage optimization strategy operating at both the structural and sequence levels. Together, these procedures form an iterative “prediction–evaluation–redesign” loop.

#### 4.5.7 Structure-based optimization

Sequences produced by EnzymeArt are folded using ESMFold to obtain predicted three-dimensional structures. Because sequence-only generation may produce local steric clashes or globally suboptimal packing, we introduce an additional structural constraint by performing inverse folding with ProteinMPNN.

Given a predicted structure S derived from an initial sequence x, ProteinMPNN generates a new amino-acid sequence x^∗^ that maximizes compatibility with the fold:

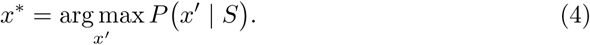

This structural refinement aligns the sequence with the geometric and physico-chemical constraints encoded in the predicted fold.

#### 4.5.8 Sequence-based optimization

To complement the structure-guided refinement, we apply ESM Cambrian (ESM C) to identify positions in the sequence that the model predicts with low certainty. For a sequence

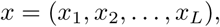

ESM C assigns a residue-level confidence score to each residue:

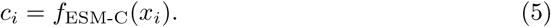

Residues with confidence values below a threshold τ are masked:

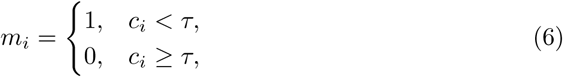

yielding a partially masked sequence x. This masked sequence is then fed back into EnzymeArt for targeted redesign:

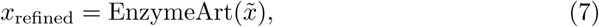

such that only low-confidence positions are modified while high-confidence regions are preserved.

*Integrated optimization pipeline.* By combining structure-level correction with confidence-guided sequence refinement, the optimization procedure converges toward sequences that are structurally coherent and compositionally robust. This dual-layer refinement is intended to improve the reliability and to increase the likelihood of obtaining biologically plausible enzyme candidates.

### 4.6 Kinetic prediction models

#### 4.6.1 EnzymeArt-KC: *k*_cat_ prediction

To enable substrate-conditioned prioritization of catalytic turnover from sequence information, we introduce EnzymeArt-KC, a predictive module trained to infer turnover number (k_cat_) from enzyme–substrate pairs. This component extends the EnzymeArt framework beyond generative design, providing a quantitative layer for function-guided candidate prioritization.

#### 4.6.2 Rank-based learning objective

Because experimentally determined k_cat_ values span several orders of magnitude and are often affected by measurement noise, we reformulate the regression objective into a rank-based learning problem. For each enzyme subclass, sequences are ranked according to their reported k_cat_ values, and the model is optimized to assign continuous scores monotonically correlated with these empirical ranks. This formulation prioritizes relative rather than absolute accuracy, yielding a noise-tolerant and scale-invariant measure of catalytic potential that is expected to generalize across heterogeneous experimental conditions.

#### 4.6.3 Representation of enzymes and substrates

To jointly capture sequence and substrate information, EnzymeArt-KC integrates pretrained biochemical language models for both molecular entities. Enzyme sequences are represented using embeddings from ESM C, which encode structural and evolutionary priors derived from large-scale protein corpora. Substrates are encoded using BioMedBERT, a transformer pretrained on biomedical text, where each input corresponds to the substrate’s natural language name (e.g., “ethanol,” “glucose-6-phosphate”). This representation allows the model to exploit semantic relations embedded in biochemical terminology, thereby placing chemically or functionally related substrates in proximal embedding regions even without explicit structural descriptions.

We freeze the pretrained encoders to preserve their biochemical priors. The resulting protein and substrate embeddings are concatenated and linearly projected into a shared latent space, providing a unified representation from which downstream layers can infer catalytic trends.

#### 4.6.4 Model architecture and optimization

The integrated embeddings are processed by a multi-layer perceptron (MLP) comprising three hidden layers with GELU activations and dropout regularization. Only the MLP parameters are trainable, ensuring computational efficiency and stable transfer from pretrained representations. Training employs the AdamW optimizer (learning rate 2 × 10^−5^, weight decay 0.01) for up to 500 epochs with early stopping based on validation rank correlation.

To improve robustness and reduce sensitivity to stochastic fluctuations during training, we employ a repeated random sub-sampling validation scheme. Specifically, the dataset is randomly partitioned into training and validation subsets multiple times, yielding 60 independently trained models under different data splits and initialization seeds. During inference, predictions from these models are averaged to form an ensemble output. This ensemble strategy helps mitigate variance, improves generalization consistency, and increases robustness across diverse enzyme subclasses.

**Fig. 7.**
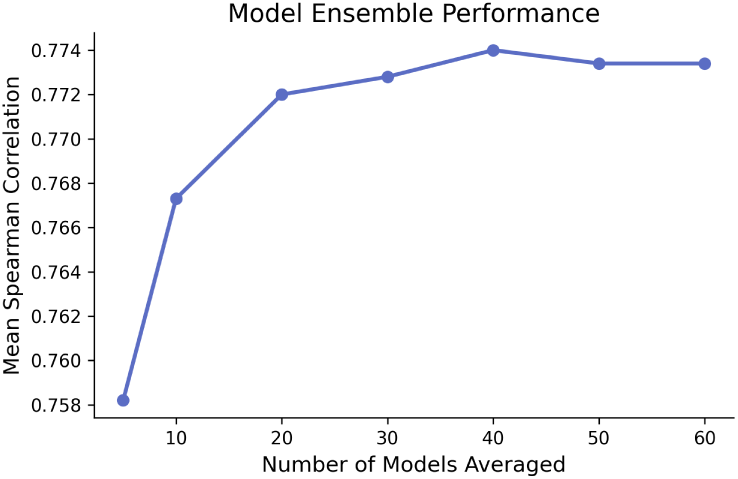
Effect of ensemble size on model performance. Increasing the number of models averaged improves the mean Spearman correlation, with performance saturating around 40 models.

#### 4.6.5 Rank-normalized kinetic labels and training objective

To remove scale dependence and stabilize optimization, all kinetic measurements are converted into rank-normalized labels. Given the rank of each kinetic value k*_i_*, computed in ascending order over N samples, we define

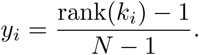

This transformation maps the smallest kinetic value to y*_i_* = 0 and the largest to y*_i_* = 1, while preserving the monotonic ordering of all intermediate measurements. The resulting labels provide a well-behaved target distribution for regression and make the learning objective invariant to the absolute scale of heterogeneous kinetic assays.

For each enzyme–substrate pair 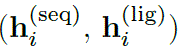, the model predicts

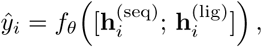

where f*_θ_* is a multi-layer perceptron applied to the joint embedding. We optimize the model by minimizing the mean squared error (MSE) between predicted and rank-normalized labels:

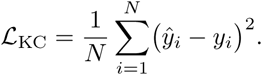

During training, we monitor validation MSE and the Spearman rank correlation between predictions ŷ*_i_* and targets y*_i_*; model checkpoints are selected based on the highest validation Spearman correlation.

#### 4.6.6 EnzymeArt-KM: KM prediction

EnzymeArt-KM extends the EnzymeArt-KC framework to predict Michaelis constants (K*_M_*) from the same enzyme–substrate representations. We reuse the frozen ESM C encoder for protein sequences and the BioMedBERT encoder for substrate names, and concatenate the resulting embeddings into a joint representation. This representation is passed through a three-layer MLP with GELU activations and dropout, identical in architecture to EnzymeArt-KC.

Because smaller K*_M_* values indicate higher substrate affinity, we preprocess the training labels so that the model still learns a “higher-is-better” kinetic score. Specifically, experimental K*_M_* measurements are first transformed to log K*_M_*, and then converted into rank-normalized affinity targets over the training set: samples with smaller K*_M_* (lower log K*_M_*) are assigned larger target values, whereas samples with larger K*_M_* receive smaller targets. EnzymeArt-KM is trained to regress these rank-normalized affinity scores using mean squared error, with the same AdamW optimizer settings and early-stopping strategy as for EnzymeArt-KC. This preprocessing ensures that, although K*_M_* itself is minimized, the learned output follows the same convention as EnzymeArt-KC, where higher predicted scores correspond to more favourable kinetic properties.

#### 4.6.7 RealKcat baseline implementation

To ensure a fair comparison with existing k_cat_ prediction approaches, we reimplemented the RealKcat prediction pipeline and trained our model on the same training split as RealKcat, followed by evaluation on the identical held-out test set. The full workflow consists of four stages: data preparation, feature extraction, model training, and performance assessment.

*Data preparation and preprocessing.* The dataset comprises enzyme–substrate pairs, each containing an amino acid sequence, a substrate SMILES string, and an experimentally determined k_cat_ value.

- **Data loading and filtering**: Raw entries are loaded from pickle files. Samples with missing fields or SMILES strings exceeding 512 characters are removed to ensure compatibility with the tokenizer.
- **Label discretization**: Continuous k_cat_ values are discretized into eight logarithmic bins using thresholds

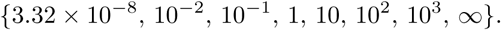
- **Dataset splitting**: Following the RealKcat setup, the training set is randomly partitioned into a 9:1 train–validation split. A separate test set (identical to RealKcat’s) is held out for final comparison.

*Feature extraction.* We adopt the same feature-generation strategy as RealKcat, using pretrained language models to encode both the enzyme and substrate representations.

- **Enzyme embedding**: Protein sequences are embedded using the esm2 t33 650M UR50D model. Representations from the final transformer layer are averaged across sequence positions to obtain a fixed-size vector.
- **Substrate embedding**: Substrate SMILES strings are encoded using seyonec/PubChem10M SMILES BPE 450k (ChemBERTa). Final-layer hidden states are mean-pooled to produce the substrate vector.
- **Feature fusion**: Enzyme and substrate embeddings are concatenated to form the input feature vector X ∈ R*^D^*.
- **Caching**: To avoid recomputation, extracted embeddings are cached using a hash-based key derived from the dataset configuration.

*Model training.* The task is framed as an eight-class classification problem, mirroring RealKcat’s discretization scheme.

- **Standardization**: Features are standardized using Z-score normalization computed from the training split. The same statistics are applied to the validation and test sets to avoid leakage.
- **Label remapping**: Class labels are remapped to a contiguous index range [0, C−1] based on the labels present in the training data.
- **Classifier**: We use XGBoost with the multi:softmax objective, following the RealKcat configuration.
- **Class balancing**: Sample weights follow scikit-learn’s balanced strategy to compensate for skewed class distributions.
- **Hyperparameters**: Key settings include

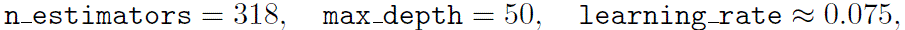

with GPU-accelerated tree method=’gpu hist’.
- **Early stopping**: Validation metrics (multi-class log loss, error rate, and AUC) are monitored with a patience of two rounds to prevent overfitting.

### 4.7 Computational evaluation and candidate prioritization (EnzymeArt-Eval)

Designed sequences undergo a multi-stage computational evaluation pipeline. Candidate pools are reduced through task-adaptive prioritization rather than a single universal scoring rule. When matched kinetic data are available, EnzymeArt-KC provides quantitative sequence–substrate ranking; in other settings, prioritization relies more strongly on structural confidence, annotation consistency, sequence-similarity or diversity filters, round-specific generation criteria, and active-site-focused structural assessment where relevant.

#### 4.7.1 Multi-stage screening of designed sequences

*Structural assessment.* Retained candidates are subsequently evaluated using the structure prediction protocol appropriate for each task. For each sequence, we record structural confidence metrics such as mean predicted local distance difference test (pLDDT), mean predicted aligned error (PAE), predicted TM score (pTM), ipTM, or related model confidence measures when available. Candidates that do not satisfy the structural criteria defined for each task are discarded.

#### 4.7.2 Sequence-level evaluation

Retained candidates are further analyzed using n-gram statistics, pseudo-perplexity (pPPL) from **ESM C**, and perplexity (PPL) from ProGen3. These metrics assess grammatical plausibility and global consistency with natural sequence distributions.

#### 4.7.3 Function evaluation

To assess whether EnzymeArt-generated sequences preserve domain architectures consistent with their intended functions, we perform domain annotation using InterProScan.

All generated sequences were analyzed using InterProScan with default parameters. For each sequence, we identified: (1) domain signatures and their InterPro (IPR) accessions, and (2) predicted catalytic or metal-binding motifs. We then evaluated the topological consistency of these annotations against known enzyme families. A design is considered functionally consistent if its domain architecture matches the canonical definition of the target class (e.g., Rossmann-like NAD-binding domains for dehydrogenases, α/β-hydrolase folds for hydrolases).

*Task-adaptive prioritization.* When matched kinetic data are available, each generated sequence paired with its target substrate is first evaluated by the EnzymeArt-KC kinetic predictor to estimate relative catalytic turnover rates (k_cat_), and only sequences exceeding a predefined percentile threshold are retained for further analysis.

*Final ranking.* Final experimental candidates are selected by integrating the evaluation signals appropriate to each task, including normalized kinetic predictions when available, structural confidence metrics, sequence-level plausibility, domain-level functional consistency, and, where relevant, local active-site structural features.

#### 4.7.4 Assessment of unconditional designs

To evaluate the intrinsic foldability and plausibility of unconditionally generated sequences, we employ a multi-tiered assessment strategy. Structural confidence is estimated using ESMFold, reporting both pLDDT and pTM as indicators of local and global structural confidence.

To assess sequence-level naturalness, we computed perplexity (PPL) using two pretrained protein language models, ESM C and ProGen3, which served as likelihood-based measures of how well designed sequences conform to the statistical distribution of natural proteins. Lower perplexity values correspond to higher biological plausibility.

#### 4.7.5 Sequence length sampling

To evaluate length-dependent behavior during unconditional generation, we adopted a controlled sequence-length sampling strategy. Target lengths are sampled from 100 to 1,000 amino acids in increments of 100.

#### 4.7.6 Structural similarity to known folds

To quantify the structural similarity of designed proteins to known natural folds, we compared each predicted structure against Protein Data Bank (PDB) library using Foldseek (GPU-accelerated version). All designed structures are first exported as individual PDB files and collected into a single directory. We then iterate over this directory and, for each design, launched easy-search against a pre-indexed PDB library using alignment-type 1 and a sensitivity parameter of 7.5 with GPU acceleration enabled. For downstream analyses, we used the top-hit TM-score as a summary measure of structural similarity between each designed protein and the closest known PDB structure, with lower TM-scores indicating more structurally novel designs.

#### 4.7.7 Functional prompt sampling and conditional design evaluation

To construct a representative benchmark for function-conditioned evaluation, we build a functional-prompt dataset by randomly sampling 1,000 enzyme sequences together with their annotated functional descriptions. Specifically, we select sequences from the curated enzyme subset of the EnzymeArt-MultiModal dataset. For each sampled sequence, the InterPro domain annotations are extracted as the functional prompt used during generation and evaluation. This sampling strategy provides broad coverage across diverse enzyme families while limiting class-specific bias. The resulting 1,000 sequence–function pairs serve as the functional evaluation set. In parallel, to evaluate EnzymeArt-base under matched conditions, we independently sample 1,000 sequences from the held-out test split of the EnzymeArt-base-MultiModal dataset and use their associated functional annotations as inputs to the model. This sampling strategy provides broad coverage across diverse enzyme families while limiting class-specific bias, and the resulting sets of 1,000 sequence–function pairs serve as the functional evaluation benchmarks for both EnzymeArt and EnzymeArt-base.

*Assessment of conditional designs and functional specificity.* For conditional designs, we evaluate functional adherence using complementary annotation-based and embedding-based analyses. First, we compare InterProScan domain assignments against the intended functional prompts to verify recovery of canonical catalytic motifs and domain architectures. Second, we quantify semantic alignment using ProTrek, a CLIP-style contrastive model that embeds protein sequences and functional text into a shared representation space.

#### 4.7.8 Function evaluation model fine-tuning (ProTrek)

To improve sensitivity to compact biochemical descriptions, we fine-tuned the ProTrek-650M dual-encoder model on a corpus of InterPro-annotated protein sequences. Each entry consisted of an amino-acid consensus sequence and one or more functional labels derived from InterPro annotations. Function strings are processed by removing identifier prefixes and concatenated into a single textual description. Sequences longer than 1,024 residues are randomly cropped per batch to avoid positional bias, whereas shorter sequences are retained in full.

Protein sequences are tokenized using the ESM-2 tokenizer (maximum length 1,024; padding and truncation enabled), and textual descriptions are tokenized using the PubMedBERT tokenizer. The model is initialized from the ProTrek-650M UniRef50 checkpoint, which aligns protein and text representations in a shared embedding space prior to task-specific adaptation.

During fine-tuning, paired protein–text embeddings are obtained via the model forward pass and optimized using the model’s built-in contrastive alignment objective, which encourages paired representations to occupy nearby regions in embedding space. Training is performed using AdamW (learning rate 1 × 10^−5^), mixed-precision computation (Bfloat16), gradient accumulation of 10 steps, and distributed data parallelism across 4 GPUs. Mini-batch sizes are 4 for training and 8 for validation. We run training for 2,500 optimization steps.

Model performance is evaluated using a held-out validation subset. For each pro-tein–text pair, cosine similarity between the respective embeddings is computed, and the mean similarity across the validation batch is reported. This measure served both as an indicator of functional alignment and the criterion for checkpoint selection.

*Cosine similarity.* To quantify the alignment between protein and textual representations, cosine similarity is computed between the respective encoder outputs. For each protein–text pair, embeddings are first obtained using the model’s inference routines, yielding fixed-dimensional vectors in the shared embedding space. Both vectors are ℓ_2_-normalized along the feature dimension, and similarity is evaluated as

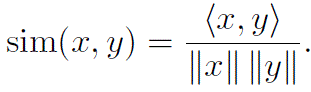

In practice, the similarity score for each pair is computed using the model’s cosine-similarity operator (F.cosine similarity) along the final embedding dimension. The resulting scalar is extracted and clipped to non-negative values to suppress spurious low-similarity noise. During validation, similarity is evaluated under mixed-precision inference (cuda autocast enabled), and the mean cosine similarity is reported as the primary metric of functional alignment.

#### 4.7.9 Subclass-wise structural confidence evaluation

We evaluated structural confidence at the EC 2-level (subclass) by aggregating all generated sequences per subclass and computing subclass-wise averages of pLDDT and pTM. The analysis covered oxidoreductases (EC 1.x), transferases (EC 2.x), hydrolases (EC 3.x), lyases (EC 4.x), isomerases (EC 5.x), ligases (EC 6.x), and translocases (EC 7.x). For each subclass, we report the total number of sequences contributing to the estimates (Sequence Count), the mean predicted local confidence (Avg pLDDT), and the mean predicted global fold integrity (Avg pTM).

All sequences are folded with the same structure prediction stack ESMFold under identical settings. We summarize model confidence using mean pLDDT and mean pTM. Subclass-wise means are computed directly from per-sequence predictions without additional weighting.

Subclass-level statistics are reported as (EC Subclass, Sequence Count, Avg pLDDT, Avg pTM) in Supplementary Table 16.

#### 4.7.10 Comparative analysis with ZymCTRL

We compare structural prediction confidence for designed sequences between EnzymeArt and ZymCTRL across a shared set of 897 enzyme entries. These entries covered two major functional classes: oxidoreductases (EC 1, 54.0%) and transferases (EC 2, 46.0%). This paired dataset ensured consistent evaluation of corresponding catalytic functions.

For each enzyme i, we collect paired confidence scores from both methods: pLDDT*_A_*(i) and pT M*_A_*(i) from EnzymeArt, and pLDDT*_B_*(i) and pT M*_B_*(i) from ZymCTRL.

For each pair, EnzymeArt scores are plotted on the x-axis and ZymCTRL scores on the y-axis. The diagonal line y = x indicates parity between methods. Color mapping represents the mean confidence across both predictions, allowing visual assessment of systematic bias and variance. All 897 paired entries contain complete and validated values within the 0–1 range, and no imputation is required.

EnzymeArt predictions exhibit higher mean confidence both locally and globally compared to ZymCTRL (Table 4).

**Table 4.** Summary statistics of structural confidence across 897 paired enzyme entries.

| Parameter | EnzymeArt | ZymCTRL |
| --- | --- | --- |
| Mean pLDDT | 0.769 | 0.550 |
| Mean pTM | 0.808 | 0.549 |
| Predictions per EC (mean ± SD) | 5.2 ± 1.0 | 3.8 ± 0.6 |

#### 4.7.11 Rosetta methods

To provide an auxiliary modeled-energy assessment of predicted structures, we compute all-atom Rosetta energies using PyRosetta (version 4).

*Initialization and structure preparation.* Energy calculations are performed using the standard full-atom (FA) Rosetta configuration. PyRosetta is initialized with default settings. Predicted or reference protein structures are loaded into a Rosetta Pose object directly from PDB files.

*Full-atom scoring function.* We use the canonical Rosetta full-atom scoring function, ref2015, initialized via the standard PyRosetta interface. This scoring function combines the standard Rosetta energetic components, including:

- Lennard–Jones attractive and repulsive energy;
- orientation-dependent hydrogen bonding;
- implicit solvation energy;
- backbone and side-chain torsional terms;
- electrostatic interactions;
- reference amino-acid energies.

*Energy calculation.* Total structure energies are obtained by scoring the pose using the ref2015 function. The resulting energy represents a weighted sum of all physical and statistical terms and provides a comparative measure of modeled conformational favor-ability within Rosetta’s energy landscape. No minimization or relaxation is applied unless otherwise stated.

#### 4.7.12 ESM3-EC baseline fine-tuning

We fine-tune the open-weight protein language model ESM3-small (ESM3 sm open v0) using parameter-efficient quantized low-rank adaptation (QLoRA). We initialize from the pretrained checkpoint, apply 4-bit quantization to the base weights, and inject LoRA adapters into all Linear layers within transformer blocks. The targeted linear layers are instantiated as 4-bit bitsandbytes modules using NF4 quantization. During training, we update only the low-rank LoRA parameters and keep all pretrained backbone weights frozen. Unless otherwise stated, we fix the LoRA hyperparameters to rank r = 8, scaling α = 16, and dropout p = 0.05. This procedure preserves the underlying model architecture and introduces only a small set of trainable adapter parameters for efficient adaptation.

Each training example comprises two aligned token tracks: (i) sequence tokens obtained by tokenizing the amino-acid sequence and (ii) auxiliary function tokens derived from the Enzyme Commission (EC) identifier in the sequence header. We parse EC identifiers using the regular expression EC[:=\s]*([0-9-]+(\.[0-9-]+)3). When no EC identifier is present, we assign a fallback label <no_ec>.

We deterministically map each EC label to a locality-sensitive hashing (LSH) token, represented as a comma-separated list of integers of length D (hereafter referred to as the hashing depth). We broadcast the same LSH token to all residue positions, such that each residue receives an identical EC-derived conditioning signal. We then encode the function track (including special tokens) to obtain a tensor of shape [L, D], where L matches the aligned sequence length (including two special positions). After padding within a batch, the sequence-token tensor has shape [B, L_max_] and the function-token tensor has shape [B, L_max_, D], aligned along the first two dimensions. The model consumes both tracks jointly during the forward pass.

The model is trained on the EnzymeArt-MultiModal dataset with a masked language modeling (MLM) objective over token sequences using cross-entropy loss. For each mini-batch, we sample a masking ratio uniformly from [0, 1] and apply it to the input tokens to obtain masked batch. Training uses a maximum sequence length of 512 tokens, a per-device batch size of 8, a peak learning rate of 1 × 10^−4^ with 2,000 warm-up steps, and a minimum learning rate of 1×10^−6^. The nominal training budget is 500,000 optimization steps.

#### 4.7.13 Baseline model evaluation

To ensure a fair comparison, all baseline models were reproduced using their official implementations and recommended inference configurations. For each model, we followed the prescribed conditioning interface, sampling parameters, and generation protocols without hyperparameter tuning unless explicitly required. All results reported in this study were generated using publicly released checkpoints.

*ESM-3.* ESM-3 was evaluated using the official inference pipeline and the 1.3B-parameter checkpoint, the largest publicly available model at release. Functional descriptions were first mapped to their corresponding InterPro (IPR) identifiers, which served as input to the model’s functional-conditioning interface. Generation followed the standard masked-refinement procedure, with the number of decoding cycles set equal to the target sequence length, as recommended by the authors. All sampling temperatures, masking schedules, logits post-processing steps, and optimization-free inference settings were left at their default values. One sequence was generated per functional prompt.

*Chroma.* Chroma was executed using the officially released functional-understanding (ProCap) module. The full functional description was supplied as free-form text, and conditioning was applied through the default ProCap caption encoder. Denoising, diffusion timestep schedules, and sampling temperatures followed the official settings. A single structure–sequence design was produced for each functional prompt without modification to the generation schedule.

*Pinal.* Pinal was evaluated using the officially released SaProt-T (760M) encoder paired with the T2Struc-1.2B structural decoder. Inference followed the default sampling configuration, including latent sampling variance, temperature, maximum decoding length, and iterative refinement settings. For each functional description, a single sequence was produced. No changes were made to any inference-time hyperparameters.

*ZymCTRL.* ZymCTRL was run using the official inference configuration, with the Enzyme Commission (EC) number provided as the sole conditioning input. Generation used nucleus sampling with stochastic decoding, top-k = 9, a repetition penalty of 1.2, a maximum sequence length of 1,024 tokens, sampling enabled (do sample = true), and four sequences per EC prompt. All other parameters, including end-of-sequence and padding token indices, were kept at default values.

*RFdiffusion + ProteinMPNN.* RFdiffusion was reproduced using the official unconditional design protocol. Backbone structures were generated at multiple target lengths using the default diffusion schedule, potentials, and sampling hyperparameters. For each specified length, 100 backbone designs were produced following the authors’ standard inference configuration.

ProteinMPNN was subsequently used to convert the RFdiffusion-generated backbones into protein sequences. The official model weights and recommended inference parameters were used, including one sequence per backbone, sampling temperature 0.1, random seed 42, and batch size 1.

We did not alter the masking procedure, side-chain treatment, or the MPNN sampling strategy from their default implementations.

Across all baselines, we follow three principles to ensure fair and reproducible comparisons. First, we use official checkpoints and inference pipelines for every model. Second, functional prompts are formatted to match each model’s native conditioning mechanism (IPR identifiers for ESM-3, free-text for Chroma and Pinal, EC numbers for ZymCTRL, and backbone-length specifications for RFdiffusion). Third, hyperparameters remain at their default values unless required otherwise by the model’s official instructions.

### 4.8 Design campaigns for experimental validation

#### 4.8.1 General workflow for function-conditioned enzyme design

Across the ADH, MDH and lipase design experiments, we follow a common stage-wise workflow for function-guided enzyme design. Each task begins with EnzymeArt-based sequence generation from fully masked sequence templates, using task-specific functional prompts and decoding settings. The resulting candidate pools are then reduced using task dependent initial screening, including kinetic ranking, generation score filtering, sequence similarity filtering or design criteria specific to each round, to remove clearly implausible candidates while preserving diversity for downstream analysis.

Surviving candidates are evaluated structurally using ESMFold or AlphaFold3 modeling as appropriate for the task. For each sequence, we record structural-confidence metrics including mean predicted local distance difference test (pLDDT), mean predicted aligned error (PAE), predicted TM-score (pTM), interface pTM (ipTM) or model ranking scores when available. Candidates with stronger structural confidence then undergo additional task-specific assessment of functional consistency, including homology search, InterProScan annotation, predicted catalytic properties or local structural inspection. Final experimental candidates are selected from these narrowed pools using criteria tailored to each enzyme class.

#### 4.8.2 Alcohol dehydrogenase design objective

We performed a dedicated set of analyses focused on EC 1.1.1.1 alcohol dehydrogenases (ADHs), including sequence landscape characterization, model-driven design and in silico screening. Experimental validation protocols are described separately below.

#### 4.8.3 EC 1.1.1.1 sequence landscape

*Dataset.* To characterize the natural sequence landscape of EC 1.1.1.1 alcohol dehydrogenases, we curate a reference set of proteins from UniProtKB. Entries are retrieved by querying UniProtKB for proteins annotated with EC 1.1.1.1, and we retain all sequences whose protein names explicitly contain the substring “1.1.1.1”. This procedure captures canonical alcohol dehydrogenases as well as multifunctional dehydrogenases with EC 1.1.1.1 among their assigned activities.

*Sequence embeddings.* To obtain a compact representation of each protein, we compute sequence embeddings using the ESM2-35M model (esm2 t12 35M UR50D). Sequences are first uppercased and cleaned by removing whitespace and stop symbols. We retain only sequences composed of the 20 standard amino acids; sequences containing non-standard characters are discarded. For numerical stability and consistent input length, sequences longer than 1,024 residues are truncated to the first 1,024 positions. We then feed each sequence through ESM2-35M and extract the representation of the special [CLS] token from the final layer as a fixed-length embedding.

*Design generation.* To investigate how EnzymeArt traverses the EC 1.1.1.1 sequence manifold, we generate sequences conditioned on natural protein names. We randomly sample natural EC 1.1.1.1 entries from the curated dataset and use their protein names as functional descriptions. Representative examples include:

- “alcohol dehydrogenase (EC 1.1.1.1)”
- “probable alcohol dehydrogenase AdhA (EC 1.1.1.1)”
- “S-(Hydroxymethyl)glutathione dehydrogenase/alcohol dehydrogenase (EC 1.1.1.1, EC 1.1.1.284)”
- “alcohol dehydrogenase (EC 1.1.1.1) (Alcohol dehydrogenase I)”

For each selected entry, we adopt the corresponding natural sequence length as the reference generation length. Conditioned on the protein name, EnzymeArt generates complete enzyme sequences at sampling temperatures ranging from 0.0 to 3.0, producing multiple candidates per prompt to explore diverse regions of sequence space. *UMAP visualization of the sequence landscape.* To visualize how EnzymeArt designs distribute relative to natural EC 1.1.1.1 proteins, we compare ESM2-35M [CLS] embeddings of natural and designed sequences using UMAP. Embeddings of natural proteins are first computed and stored as a reference. For EnzymeArt designs, we apply the same embedding protocol (cleaning, truncation, and ESM2-35M [CLS] extraction), and concatenate the resulting embeddings with the natural set into a single matrix.

We then apply UMAP (as implemented in umap-learn) with a cosine distance metric and a fixed random seed (random state = 42) to project the combined embeddings into two dimensions. Let N_nat_ denote the number of natural sequences and N_des_ the number of designed sequences. UMAP coordinates for the first N_nat_ points correspond to natural proteins, and the remaining N_des_ points correspond to EnzymeArt designs.

To characterize the density of the natural sequence manifold, we estimate a two-dimensional probability density over the UMAP coordinates of natural proteins using Gaussian kernel density estimation (KDE). We evaluate the KDE on a regular grid spanning the UMAP plane and render the result as grayscale filled contours, providing a smooth density background for the natural EC 1.1.1.1 landscape. Natural sequences are optionally plotted as faint grey points over this density map.

EnzymeArt designs are plotted as overlaid points in a contrasting purple color. This visualization highlights how generated sequences populate both high-density regions of the natural manifold and lower-density peripheral regions, illustrating the coverage and extrapolation behavior of EnzymeArt within the EC 1.1.1.1 sequence landscape (Fig. 8).

**Fig. 8.**
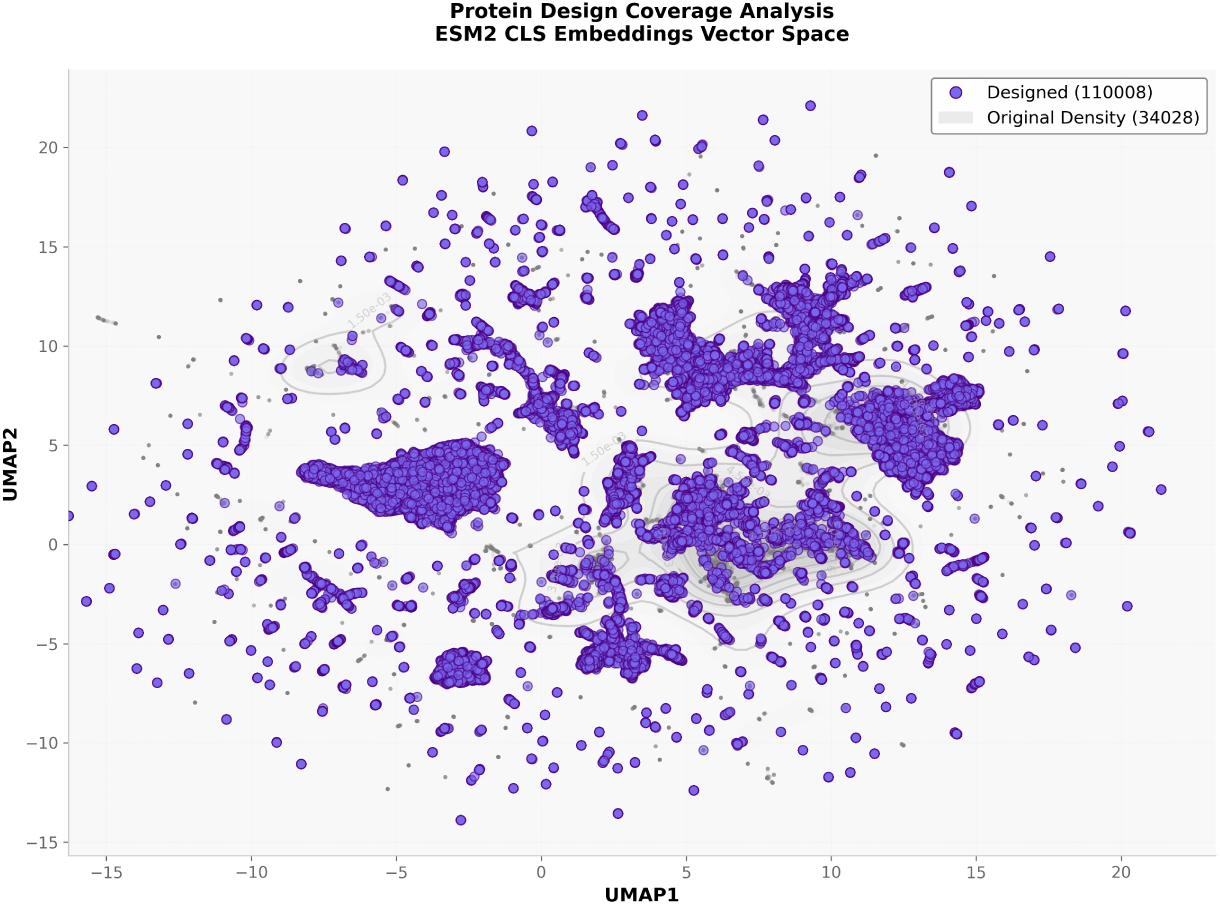
Embedding-space coverage of generated enzyme designs.

#### 4.8.4 ADH design prompts and decoding configuration

To evaluate the model’s ability to generate alcohol dehydrogenases under rich functional guidance, we condition EnzymeArt on a comprehensive prompt that integrates catalytic classification, family-level features and conserved structural domains for EC 1.1.1.1. Unless otherwise stated, ADH design experiments use a target length of 339 residues and T = 500 refinement cycles, with sampling temperature and Gumbel noise scale linearly decayed from their initial values to zero. The input specification is:

“alcohol dehydrogenase (EC 1.1.1.1), Alcohol dehydrogenase, zinc-type, conserved site, GroES-like superfamily, Alcohol dehydrogenase-like, C-terminal, Alcohol dehydrogenase-like, N-terminal, NAD(P)-binding domain superfamily”

To translate functional descriptions into protein sequences, we use a multi-stage decoding procedure that combines stochastic exploration with structured diffusion-style denoising. Generation starts from a fully masked sequence and proceeds for T decoding steps. At each step, the model predicts amino-acid distributions conditioned on a functional embedding obtained from a frozen biomedical language model. High-confidence predictions are progressively fixed, while low-confidence positions remain flexible. All hyperparameters used in this procedure are provided explicitly below.

*Stochastic token sampling via Gumbel–softmax.* At decoding step t ∈ {0, …, T − 1}, the model outputs amino-acid logits for each sequence position. Before sampling, all special tokens (<mask>, <pad>, <bos>, <eos>, and unused reserved indices in [24, 31]) are masked by assigning logits of −∞. Sampling is performed using Gumbel–softmax with:

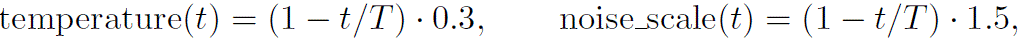

where the temperature controls the entropy of the categorical distribution and the noise scale determines the magnitude of injected Gumbel noise. The temperature therefore linearly decays from 0.3 at t = 0 (high exploration) to 0 at t = T (deterministic decoding), and the noise scale similarly decays from 1.5 to 0.

**Algorithm 5.**
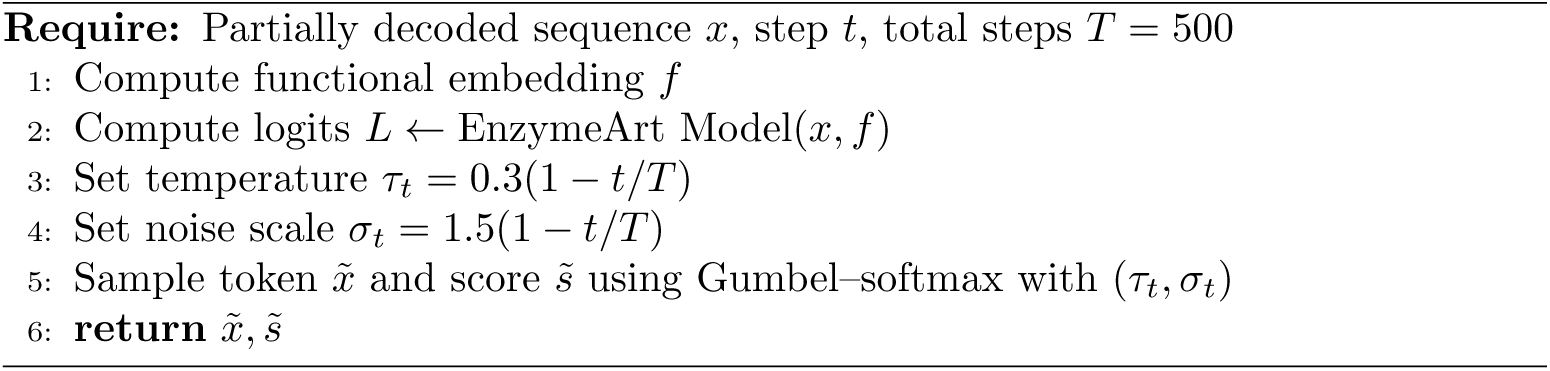
Stochastic token sampling at decoding step t.

*Frequency-aware conditional resampling.* To prevent degeneracy caused by over-repetition of a single amino acid, we use a conditional resampling step. After each sampling pass, any sequence in which a token occupies more than 15% of all non-special positions is marked for correction. All occurrences of these over-represented tokens are temporarily re-masked.

To introduce controlled stochasticity, resampling is then performed by adding Gumbel noise with temperature fixed to zero and a noise scale matched to the main sampler’s σ*_t_*, which preserves functional constraints while maintaining sufficient diversity.

*Progressive reparameterized denoising.* After sampling, we apply a structured denoising update that gradually fixes high-confidence positions. At decoding step t, a fraction of the valid positions is selected for denoising according to a monotonic schedule:

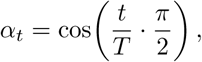

which decreases smoothly from α_0_ ≈ 1 to α*_T_* = 0. Thus, the number of positions updated at step t is:

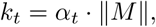

where M denotes the mask of non-special positions. Among these, the k*_t_* lowest-confidence positions (based on sampled scores 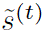) are treated as “noise” and remain masked; all remaining positions adopt the newly sampled tokens. This progressively reduces uncertainty while ensuring late-stage convergence.

**Algorithm 6.**
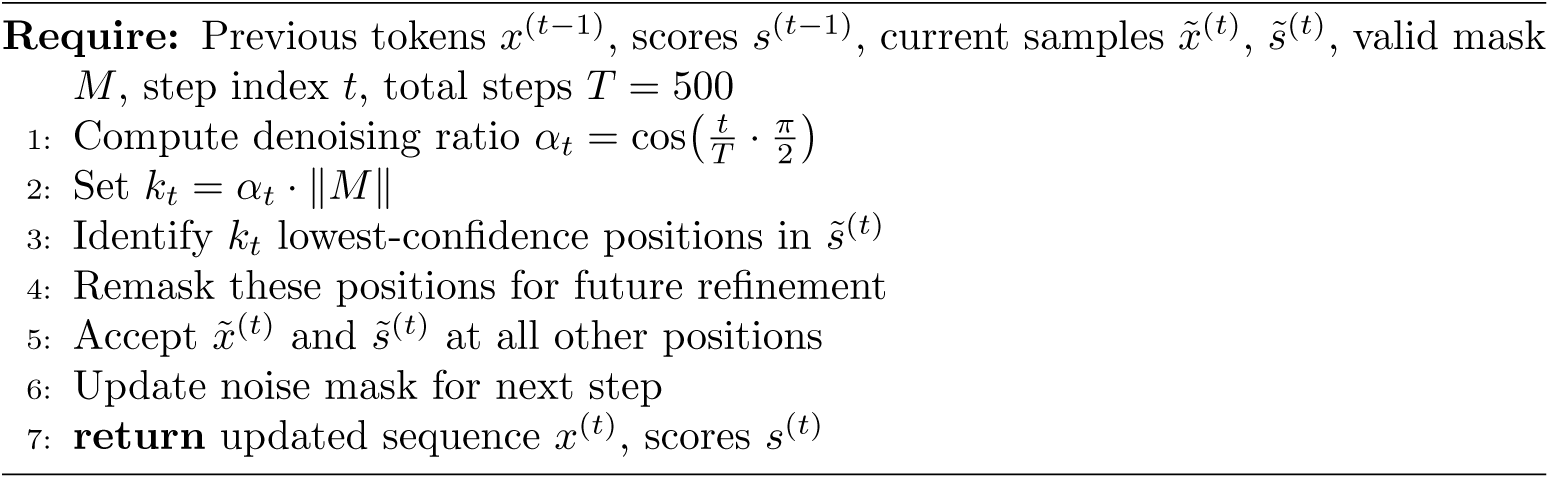
Reparameterized denoising step.

#### 4.8.5 In silico screening of designed ADHs

To characterize the biochemical plausibility of EnzymeArt-generated ADH sequences, we implement a multi-stage evaluation pipeline that integrates kinetic prediction, structure modeling, sequence-level naturalness assessment, and domain-level functional verification.

*Designs without post-generation optimization.* To assess the intrinsic reliability of the EnzymeArt model, we evaluated ADH designs produced without any structure– or sequence-level optimization. In this setting, sequences were sampled directly from the model conditioned only on their functional descriptions. This provides a direct test of whether EnzymeArt alone can generate biochemically plausible enzymes.

All screening evaluations are performed using the designed sequence and ethanol as the target substrate.

*Kinetic activity prediction.* To specialize EnzymeArt-KC for alcohol dehydrogenase (ADH, EC 1.1.1.1) activity prediction, we curate a refined subset of the RealKcat database. Specifically, we replace the original EC 1.1.1.1 entries with a newly compiled collection of 572 experimentally validated k_cat_ measurements for ADH enzymes. These records are aggregated from BRENDA, with each entry containing the enzyme sequence (UniProt accession), substrate name, and standardized turnover number (k_cat_, s^−1^).

To ensure data reliability, we remove inconsistent or duplicated measurements and average multiple reports for the same enzyme–substrate pair. The dataset covers both NAD^+^– and NADP^+^-dependent ADHs, spanning a dynamic range of approximately 10^−2^–10^3^ s^−1^. Fifty percent of the dataset is reserved as a held-out test set, while the remaining data are randomly partitioned into training and validation subsets in a 9:1 ratio. This configuration provides a consistent basis for evaluating model generalization within the dehydrogenase family. For each designed sequence x, we pair it with ethanol and evaluate the catalytic turnover rate using the EnzymeArt-KC. For each enzyme–substrate pair, the EnzymeArt-KC directly outputs predicted k_cat_ rank values, which are averaged to obtain the final activity score. Sequences with low predicted catalytic potential are discarded at this stage.

*Structure prediction with AlphaFold3.* Sequences that pass kinetic screening are structurally assessed using AlphaFold3. We provide the amino acid sequence together with ethanol as a ligand input. AlphaFold3 generates a predicted enzyme–substrate complex and associated structure confidence metrics including pLDDT, pTM, and interface pTM (ipTM). Designs with pLDDT < 0.85 or ipTM < 0.85 are removed from further analysis.

*Sequence-level naturalness analysis.* Remaining sequences are evaluated for grammatical consistency and compatibility with natural protein sequence statistics. We compute:

- pseudo-perplexity (pPPL) using ESM C to quantify residue-level predictability under a structural language model,
- perplexity (PPL) using ProGen3 to assess global sequence plausibility,

Designs exhibiting extreme pPPL or PPL values inconsistent with natural dehydrogenase families are filtered out.

*Functional domain and motif validation.* To ensure that the designed ADHs recapitulate the structural logic of natural enzymes, we annotated sequences using InterProScan with standard databases (including Pfam, PROSITE, and CATH-Gene3D). For each candidate sequence, we validated:

1. the presence of distinct ADH-family domains (e.g., Zinc-binding dehydrogenases);
2. sequence signatures associated with the NAD(P)-binding Rossmann fold;

A design is considered functionally consistent if all required domains are present and aligned with typical positions observed in zinc-dependent alcohol dehydrogenases.

*Final selection.* Only sequences that satisfy all criteria, including high predicted k_cat_, structural reliability in AlphaFold3, natural sequence statistics, and correct domain architecture, are retained as high-confidence ADH candidates for downstream experimental validation.

**Table 5.**
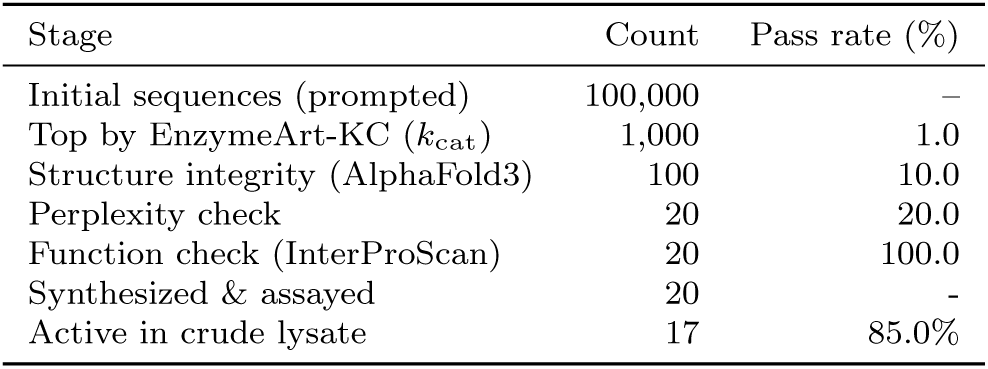
ADH candidate filtering and experimental validation pipeline.

| Stage | Count | Pass rate (%) |
| --- | --- | --- |
| Initial sequences (prompted) | 100,000 | – |
| Top by EnzymeArt-KC ($k_{\text{cat}}$) | 1,000 | 1.0 |
| Structure integrity (AlphaFold3) | 100 | 10.0 |
| Perplexity check | 20 | 20.0 |
| Function check (InterProScan) | 20 | 100.0 |
| Synthesized & assayed | 20 | – |
| Active in crude lysate | 17 | 85.0% |

#### 4.8.6 Representative prompts across enzyme families

#### 4.8.7 MDH design and candidate selection

The MDH campaign follows the same staged prioritization logic as the ADH campaign, with MDH-specific functional conditioning, structural filtering, catalytic-property prediction and novelty filtering before experimental selection.

**Table 6.** Target sequence lengths and functional prompts across representative enzyme classes.

| Enzyme Class / Family | Length (aa) | Functional Prompt |
| --- | --- | --- |
| Serine proteases | 220 | Complement factor D (EC 3.4.21.46); Adipsin; C3 convertase activator; Properdin factor D |
| Adenosine kinase | 166 | Adenosine kinase (AK) (EC 2.7.1.20); Adenosine 5'-phosphotransferase |
| Enolase | 348 | Enolase (EC 4.2.1.11) |
| Triosephosphate isomerase | 269 | Triosephosphate isomerase (TIM/TPI) (EC 5.3.1.1); Triose-phosphate isomerase |

*Generation and score-based pre-screening.* To evaluate whether EnzymeArt generalizes to a distinct dehydrogenase family, we apply the same overall design and screening strategy used for ADH to malate dehydrogenase (MDH; EC 1.1.1.37). The MDH campaign uses an MDH-specific functional prompt:

“Malate dehydrogenase (EC 1.1.1.37), Lactate/malate dehydrogenase, N-terminal, Lactate/malate dehydrogenase, C-terminal, NAD(P)-binding domain superfamily”

Sequence generation is performed with a target length of 320 amino acids.

The MDH design campaign generated an initial in silico pool of approximately 10^6^ candidate sequences. To reduce the cost of downstream structure prediction, we retained a high-confidence subset of approximately 10^5^ candidates according to the model-derived generation score. This pre-screening step was based only on the generation score and did not use structural information, sequence-similarity filtering or active-site annotation.

*Structure– and domain-based filtering.* The score-filtered MDH sequences are subjected to structure prediction using ESMFold. For each completed prediction, global structure-confidence metrics, including mean pLDDT, mean PAE and pTM, are recorded. Candidates with mean pLDDT greater than 0.80 are retained for downstream analysis.

To verify that the retained sequences preserve MDH-related domain organization, the structurally filtered candidates are further analysed using InterProScan. Candidates are prioritized if they contain annotations consistent with the lactate/malate dehydrogenase family and the expected NAD(P)-binding domain context. This step removes sequences with high predicted fold confidence but inconsistent or incomplete MDH-family domain annotations.

*Catalytic-property prediction and AlphaFold3-based complex assessment.* The domain-consistent MDH candidates are next evaluated using substrate-conditioned catalytic-property prediction with malate as the substrate. EnzymeArt-KC is used to predict k_cat_, and EnzymeArt-KM is used to predict K*_M_*. These predictions are used to estimate the catalytic potential of each candidate on malate, and candidates with stronger predicted malate-turnover properties are prioritized for further structural assessment. Shortlisted candidates are then subjected to AlphaFold3 complex modeling with malate as the substrate. The AlphaFold3 ranking score is used as the primary criterion for assessing the predicted compatibility between each designed MDH candidate and malate in the modelled complex. Candidates with low ranking scores or implausible enzyme–substrate complex models are deprioritized before final synthesis selection. *Similarity filtering and final candidate selection.* To assess sequence novelty, the remaining candidates are compared against known proteins using BLASTP. Designed candidates are retained only if their maximum detected sequence identity is below 50%, ensuring that the final design set contains sequences substantially diverged from known MDH-related proteins.

After screening by generation score, ESMFold-based structural filtering, InterProScan domain validation, AlphaFold3 structural assessment and BLASTP similarity filtering, 20 designed MDH candidates are selected for synthesis. Two natural MDH sequences are included as experimental controls, yielding 22 sequences for wet-lab testing. This screening strategy follows the same broad logic of staged computational prioritization used in the ADH campaign, while adapting the functional prompt, substrate-conditioned prediction and similarity-filtering criteria to the MDH design task.

#### 4.8.7 Lipase design and candidate selection

The lipase campaign applies the shared EnzymeArt design and prioritization workflow to a hydrolase target, using lipase-specific prompts, substrate-aware catalytic prediction and assay substrates relevant to triglyceride hydrolysis.

*Generation and score-based pre-screening.* To evaluate whether the EnzymeArt design workflow generalizes beyond nicotinamide-dependent dehydrogenases, we apply the same overall generation and screening strategy to triacylglycerol lipase design. The lipase campaign uses a lipase-specific functional prompt that combines the catalytic annotation Triacylglycerol lipase (EC 3.1.1.3) with family– and domain-level descriptions.

“Triacylglycerol lipase (EC 3.1.1.3), Triacylglycerol lipase family, PLAT/LH2 domain, Pancreatic lipase, Lipase, Lipase, LIPH-type, Alpha/Beta hydrolase fold, Lipase, N-terminal, PLAT/LH2 domain superfamily”

In the principal prompt-only setting, EnzymeArt generates fully masked sequences with a target length of 253 amino acids. The initial lipase design campaign generated an in silico pool of approximately (5 × 10^5^) candidate sequences. Stochastic decoding is performed for 500 denoising iterations, with the sampling temperature randomly selected between 0.8 and 2.0. The resulting lipase candidate pool is then subjected to the same staged computational prioritization framework used for the ADH and MDH design campaigns.

To reduce the cost of downstream structure prediction, the initial lipase design pool is first filtered according to the model-derived generation score. This score-based prescreening step is used only to retain high-confidence generated sequences for further evaluation. No structure prediction, sequence-similarity filtering, substrate-complex modeling or manually specified active-site constraint is used at this stage.

*Structure– and domain-based filtering.* The score-filtered lipase candidates are next subjected to structure prediction using ESMFold. For each completed prediction, global structural-confidence metrics, including mean pLDDT, mean PAE and pTM, are recorded. Candidates with stronger predicted structural confidence are retained for downstream analysis, whereas sequences with low-confidence or incomplete predicted folds are deprioritized.

To evaluate whether the retained structures are consistent with the intended lipase function, structurally filtered candidates are analysed using InterProScan. Candidates are prioritized if their annotations are consistent with lipase-related domain organization, including alpha/beta hydrolase fold annotations and lipase-associated N-terminal or PLAT/LH2-related domain context. This step removes candidates that show high predicted fold confidence but lack domain-level evidence supporting the intended triacylglycerol lipase function.

*Catalytic-property prediction and AlphaFold3-based substrate-complex assessment.* The lipase candidates that passed domain validation are then evaluated using catalytic property prediction conditioned on the target substrates. Because the experimental validation focuses on triglyceride hydrolysis, candidates are scored against the triglyceride substrates used in the downstream assays, including tributyrin and triolein. EnzymeArt-KC and EnzymeArt-KM were used to estimate relative catalytic turnover potential and apparent substrate affinity, respectively. UniKP predictions were further included as an independent substrate-aware kinetic assessment. Together, these scores were used to prioritize lipase candidates predicted to have stronger activity toward the tested substrates.

Shortlisted candidates are then evaluated by AlphaFold3 modeling of enzyme substrate complexes. These models are used to assess whether each candidate maintains a confident lipase fold in the presence of the target substrate. The AlphaFold3 ranking score and the overall plausibility of the predicted enzyme substrate complex are used as the main criteria at this stage. Candidates with low ranking scores, poorly resolved complex models or structures that appear incompatible with substrate binding are deprioritized before final synthesis selection. This step serves as a substrate aware structural assessment, rather than a manually specified active site or residue level constraint.

*Similarity filtering and final candidate selection.* To assess sequence novelty and avoid selecting close rediscoveries of known lipases, the remaining candidates are compared against known proteins using BLASTP. For each candidate, the maximum detected sequence identity among BLASTP hits is recorded. This analysis is used to characterize the relationship between the designed lipase candidates and known natural proteins and to support final candidate selection.

After prescreening by generation score, structural filtering with ESMFold, domain validation with InterProScan, substrate conditioned catalytic property prediction, AlphaFold3 substrate complex assessment and BLASTP similarity analysis, a compact set of lipase candidates is selected for synthesis and experimental testing. The workflow follows the same overall logic as the ADH and MDH campaigns, but is tailored to triacylglycerol lipase design through a lipase specific functional prompt, substrate aware prioritization and assay substrates relevant to triglyceride hydrolysis.

### 4.9 Experimental validation

For the experimentally tested ADH, MDH and lipase panels, candidates were generated from function-conditioned prompts without fixing, grafting or manually specifying catalytic residues, active-site motifs or metal-binding geometries during generation or final wet-lab candidate selection.

#### 4.9.1 Gene synthesis, cloning and expression

All designed enzyme sequences were codon-optimized for expression in *Escherichia coli* BL21(DE3). The optimized genes were synthesized and cloned into pET-28a(+) using Nde I/Xho I sites for ADH and MDH, and Nco I/Xho I sites for lipase. The cloning strategy placed an N-terminal His_6_ tag on ADH and MDH proteins and a C-terminal His_6_ tag on lipase proteins. Recombinant plasmids were transformed into chemically competent *E. coli* BL21(DE3) cells by heat shock. Full amino acid sequences are provided in Sections 7.10, 7.11 and 7.13, and in Table 23, 24, and 27.

Single colonies were inoculated into LB medium containing kanamycin (50 µg mL^−1^) and grown at 37 ^◦^C. Seed cultures were transferred and grown to OD_600_ = 0.6 and induced with 0.25 mM IPTG. Expression was carried out at 16 ^◦^C with shaking at 220 rpm for 12–16 h. Cells were harvested by centrifugation at 6000×g for 10 min at 4 ^◦^C. Cell pellets were lysed with B-PER reagent (Thermo Fisher, Cat. 78243), and debris was removed by centrifugation at 12000×g for 30 min at 4 ^◦^C.

#### 4.9.2 Protein purification

A HisTrap HP Ni^2+^-affinity column (Cytiva) was equilibrated with buffer A (100 mM potassium phosphate, 150 mM NaCl, pH 8.0) on an A KTA Avant FPLC system (Cytiva). Clarified lysates from the expression cultures were then loaded onto the column. The column was washed with 5% buffer B, corresponding to 25 mM imidazole, and bound proteins were eluted with 60% buffer B, corresponding to 300 mM imidazole. Buffer B contained 100 mM potassium phosphate, 150 mM NaCl and 500 mM imidazole at pH 8.0. Eluates were desalted into storage buffer (100 mM potassium phosphate, 100 mM NaCl, 1 mM DTT, pH 8.0) using PD-10 columns (Cytiva). Protein purity was assessed by SDS–PAGE, and protein concentration was determined by absorbance using a NanoDrop Lite Plus spectrophotometer (Thermo Fisher).

#### 4.9.3 Crude-lysate activity assays

*ADH and MDH crude-lysate assays.* Crude activities of ADH and MDH candidates were measured with a WST-8 coupled assay kit (Beyotime, Cat. S0241M). Reactions (100 µL) contained 90 µL working solution and 10 µL crude lysate. The working solution contained WST-8, NAD^+^ and substrate. ADH assays used 10 mM ethanol, and MDH assays used 2 mM L-malate. Absorbance at 450 nm was recorded every 1 min for 10 min at 37 ^◦^C using a Synergy H1 microplate reader. Crude-lysate screening signals are summarized in Supplementary Tables 12 and 13.

*Lipase crude-lysate assays.* Lipase hydrolysis reactions (500 µL) contained 100 mM potassium phosphate buffer (pH 8.0), 2 mM triglyceride substrate (tributyrin or triolein) and 50 µL crude lysate. Reactions were incubated at 37 ^◦^C with shaking at 200 rpm for 30 min and terminated at 100 ^◦^C for 10 min. Glycerol concentrations in the hydrolysis mixtures were measured using an Amplex Red assay (Beyotime, Cat. S0223M). One triglyceride molecule yields one glycerol molecule and three free fatty acids upon complete hydrolysis. For glycerol detection, 100 µL reactions contained 80 µL working solution and 20 µL hydrolysis mixture. After incubation at 37 ^◦^C for 30 min, absorbance at 570 nm was measured using a Synergy H1 microplate reader. Crude-lysate screening signals are summarized in Supplementary Table 14.

#### 4.9.4 Purified-enzyme steady-state kinetic measurements

*ADH kinetic measurements.* Purified ADH kinetics were measured in 100 µL reactions containing purified enzyme (0.05 µg µL^−1^ final concentration), 100 mM potassium phosphate buffer (pH 8.0), 2 mM NAD^+^ and ethanol at 0.1–40 mM. Reactions were run at 37 ^◦^C, and absorbance at 340 nm was recorded every 1 min for 10 min. Enzyme-free background controls were subtracted. Max reaction velocities were estimated by linear regression of the time-dependent absorbance traces and fit to the Michaelis–Menten equation in GraphPad Prism 9. Experiments were performed in triplicate. The fitted parameters are summarized in Table 7.

**Table 7.**
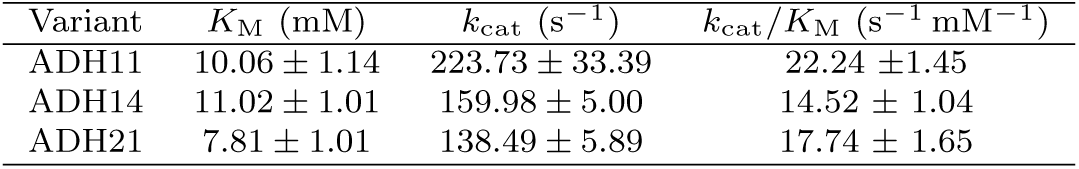
Kinetic parameters of purified ADH variants toward ethanol.

*MDH kinetic measurements.* Purified MDH kinetics were measured using the same reaction format, background subtraction and Michaelis–Menten fitting procedure as the ADH kinetic assays. Purified enzyme was used at 0.01 µg µL^−1^ final concentration, and L-malate was varied from 0.01 to 2 mM. MDH19 was assayed together with natural MDH controls corresponding to UniProt accessions G4T3S9 and A0A0F2NS13. The fitted parameters are summarized in Table 8.

**Table 8.** Kinetic parameters for MDH19 and natural MDH controls. MDH19 is the EnzymeArt design. MDH21 and MDH22 are natural MDH controls corresponding to UniProt accessions G4T3S9 and A0A0F2NS13, respectively.

| Enzyme | Source | Nearest-hit identity or ID | $K_M$ (μM) | $k_{cat}$ (s<sup>-1</sup>) | $k_{cat}/K_M$ (mM<sup>-1</sup> s<sup>-1</sup>) |
| --- | --- | --- | --- | --- | --- |
| MDH19 | EnzymeArt design | 33% BLASTP identity | 778.80±67.20 | 267.57±10.05 | 343.52±18.18 |
| MDH21 | Natural MDH control | UniProt: G4T3S9 | 121.53±9.12 | 608.97±11.76 | 5010.78±309.42 |
| MDH22 | Natural MDH control | UniProt: A0A0F2NS13 | 161.29±12.87 | 809.23±17.87 | 5017.20±321.34 |

All Michaelis–Menten fits gave $R^2 = 0.99$ .

*Lipase kinetic measurements.* Purified lipase kinetics were measured in 500 µL reactions containing purified enzyme (0.01 µg µL^−1^ final concentration), 100 mM potassium phosphate buffer (pH 8.0) and tributyrin or triolein at 0.01–1.8 mM. Reactions were incubated at 37 ^◦^C with shaking at 200 rpm for 10 min and terminated at 100 ^◦^C for 10 min. Glycerol was quantified using the same Amplex Red absorbance assay described for crude lipase measurements. Lipase-catalysed hydrolysis was quantified by endpoint measurement of glycerol formation using the commercial glycerol assay kit (Beyotime, Cat. S0223M). We used the 10-min endpoint consistently across all substrate concentrations and calculated the mean reaction velocity (V_mean_) from glycerol formed over the incubation interval. These endpoint-derived velocities were fit to the Michaelis–Menten equation in GraphPad Prism 9. Experiments were performed in triplicate. Because reaction velocities were estimated from fixed-time endpoint measurements rather than continuously monitored initial rates, the resulting lipase kinetic parameters are reported as apparent kinetic parameters, 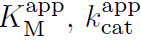 and 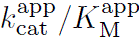.

**Table 9.** Apparent kinetic parameters for Lipase15 and the commercial lipase control. 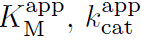 and 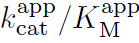 are endpoint-derived apparent parameters.

| Substrate | Enzyme | $K_{\text{M}}^{\text{app}}$<br>($\mu\text{M}$) | $k_{\text{cat}}^{\text{app}}$<br>($\text{s}^{-1}$) | $k_{\text{cat}}^{\text{app}}/K_{\text{M}}^{\text{app}}$<br>($\text{s}^{-1} \text{mM}^{-1}$) | $R^2$ |
| --- | --- | --- | --- | --- | --- |
| Tributyrin | Lipase15 | $220.79 \pm 25.56$ | $24.81 \pm 0.87$ | $112.38 \pm 10.03$ | 0.977 |
| Tributyrin | Commercial lipase | $239.30 \pm 31.64$ | $22.99 \pm 0.94$ | $96.08 \pm 9.72$ | 0.970 |
| Triolein | Lipase15 | $287.50 \pm 58.52$ | $20.61 \pm 1.35$ | $71.67 \pm 10.90$ | 0.939 |
| Triolein | Commercial lipase | $314.10 \pm 40.35$ | $19.69 \pm 0.84$ | $62.70 \pm 5.92$ | 0.975 |

$K_{\text{M}}^{\text{app}}$ denotes the apparent Michaelis constant estimated from endpoint assays; $k_{\text{cat}}^{\text{app}}$ denotes the apparent turnover number calculated over the 10-min hydrolysis period.

#### 4.9.5 Control enzymes and assay reagents

For ADH validation, ADH21 corresponds to the wild-type reference enzyme from *Saccharomyces cerevisiae* (UniProt P00330), a canonical yeast ADH1 benchmark with a curated BRENDA ethanol turnover number of 143 s^−1^ under pH 8.0 and 25 ^◦^C conditions. This reference therefore represents a well-characterized, catalytically competent wild-type ADH benchmark rather than a weak or poorly annotated comparator. For MDH validation, MDH21 and MDH22 correspond to UniProt accessions G4T3S9 and A0A0F2NS13, respectively. For lipase validation, crude-lysate assays included a natural lipase reference (UniProt P61872) and a commercial lipase preparation, and purified kinetic assays compared Lipase15 with the commercial lipase control under matched substrate and assay conditions.

**Table 10.** Control enzymes, assay reagents and external records used for experimental validation.

| 2441 | Item | Role in experiment | Identifier | Link or record |
| --- | --- | --- | --- | --- |
| 2442 | ADH21 | Natural ADH control | UniProt: P00330 | UniProt accession P00330 |
| 2443 | MDH21 | Natural MDH control | UniProt: G4T3S9 | UniProt accession G4T3S9 |
| 2444 | MDH22 | Natural MDH control | UniProt: A0A0F2NS13 | UniProt accession A0A0F2NS13 |
| 2445 | P61872 | Natural lipase reference | UniProt: P61872 | UniProt accession P61872 |
| 2446 | Commercial lipase | Lipase positive control | Macklin L812480 | https://www.macklin.cn/products/L812480 |
| 2447 | ADH activity assay kit | ADH activity measurement | Beyotime S0241M | https://www.beyotime.com/product/S0241M.htm |
| 2448 | Glycerol assay reagent | Lipase product assay | S0223M | https://www.beyotime.com/product/S0223M.htm |
| 2449 |  |  |  |  |
| 2450 |  |  |  |  |
| 2451 |  |  |  |  |
| 2452 |  |  |  |  |

#### 4.9.6 Relative comparison between Lipase15 and the commercial lipase control

Relative values in Table 11 were calculated as Lipase15 divided by the commercial lipase control for the same substrate.

**Table 11.** Relative kinetic comparison between Lipase15 and the commercial lipase control. Ratios are calculated within each substrate.

| 2463 | Substrate | Metric | Lipase15/control ratio | Difference |
| --- | --- | --- | --- | --- |
| 2464 | Tributyrin | $K_M^{app}$ | 0.92 | 7.7% lower for Lipase15 |
| 2465 | Tributyrin | $k_{cat}^{app}$ | 1.08 | 7.9% higher for Lipase15 |
| 2466 | Tributyrin | $k_{cat}^{app}/K_M^{app}$ | 1.17 | 16.9% higher for Lipase15 |
| 2467 | Triolein | $K_M^{app}$ | 0.92 | 8.5% lower for Lipase15 |
| 2468 | Triolein | $k_{cat}^{app}$ | 1.05 | 4.6% higher for Lipase15 |
| 2469 | Triolein | $k_{cat}^{app}/K_M^{app}$ | 1.14 | 14.3% higher for Lipase15 |

#### 4.9.7 Controls, background subtraction and statistical analysis

For crude-lysate screening, each design was tested in three independent experiments together with matched assay-background and empty-vector controls. No-substrate controls were included where applicable. A design was classified as active only when substrate-dependent activity was reproducibly detected above the corresponding matched background and empty-vector controls. Where statistical comparisons are reported, crude-lysate activities were compared with the empty-vector control using a two-sided Welch’s t-test. For purified-enzyme kinetic measurements, enzyme-free reactions were used for background subtraction before velocity estimation and Michaelis–Menten fitting.

### 4.10 Natural-sequence similarity analysis using MMseqs2

To compare the positions of ZymCTRL– and EnzymeArt-generated sequences in natural protein sequence space, we search both sequence sets against a local UniRef50 database with GPU-accelerated MMseqs2. We use UniRef50 as the reference database and convert it into an MMseqs2 database with mmseqs createdb, followed by conversion to a GPU-padded database with mmseqs makepaddedseqdb for efficient CUDA-based search over the full reference set. According to the MMseqs2 lookup table, the final database contains 60,315,044 sequences. We analyse two query sets, zymctrl common ec.fasta with 3,424 sequences and enzymeart common ec.fasta with 4,736 sequences. All searches are run on a single 8-GPU node, with each query set partitioned into four shards and each shard processed independently on one GPU. We run MMseqs2 search with –s 7.5, –-alignment-mode 3, –-max-seqs 300, and –-gpu 1, which balance sensitivity and throughput while retaining alignment-derived statistics for downstream analysis. We then extract the top-ranked hit for each query with mmseqs filterdb –-extract-lines 1. Throughout this section, “all-hit” denotes the complete set of retained hits under the –-max-seqs 300 limit rather than all possible matches in the database.

The MMseqs2 summary statistics are reported in Supplementary Table 15.

## 5 Data availability

EnzymeArt-base protein sequences used for training and evaluation were obtained from the AFDB50 (Foldseek) database. EnzymeArt protein sequences and enzyme names and EC numbers used for training and evaluation were obtained from the UniProtKB database (https://www.uniprot.org/). InterPro domain descriptions for all sequences were retrieved from the InterPro database v104.0 (https://www.ebi.ac.uk/interpro/). Enzyme–reaction pairings and catalytic classifications were derived from publicly accessible biochemical resources, including the RealKcat database (https://chowdhurylab.github.io/downloads.html) and the BRENDA database (https://www.brenda-enzymes.org/). Experimental structures from the Protein Data Bank (PDB; accession 1HSO) are used in this study.

## 6 Code availability

Source code for the EnzymeArt model, EnzymeArt-KC, EnzymeArt-KM, trained weights and inference scripts will be made available under an open-source license upon publication.

Neural networks were developed with PyTorch v.2.1.0 (https://github.com/pytorch/pytorch), PyTorch Lightning v.2.2.0 (https://github.com/Lightning-AI/pytorch-lightning), FlashAttention v.2.6.3 (https://github.com/Dao-AILab/flash-attention).

Data analysis used Python v.3.11 (https://www.python.org), pandas v.2.3.1 (https://github.com/pandas-dev/pandas), NumPy v.2.3.2 (https://github.com/numpy/numpy), Matplotlib v.3.10.5 (https://github.com/matplotlib/matplotlib), umap-learn v.0.5.9.post2 (https://github.com/lmcinnes/umap), InterPro (https://www.ebi.ac.uk/interpro), foldseek Release 10 (https://github.com/steineggerlab/foldseek) was used for computing TM-scores. Structure visualizations were created in PyMOL v.3.1.6.1 (https://github.com/schrodinger/pymol-open-source).

## 7 Supplementary

### 7.1 Crude-lysate activity screening tables

**Table 12.**
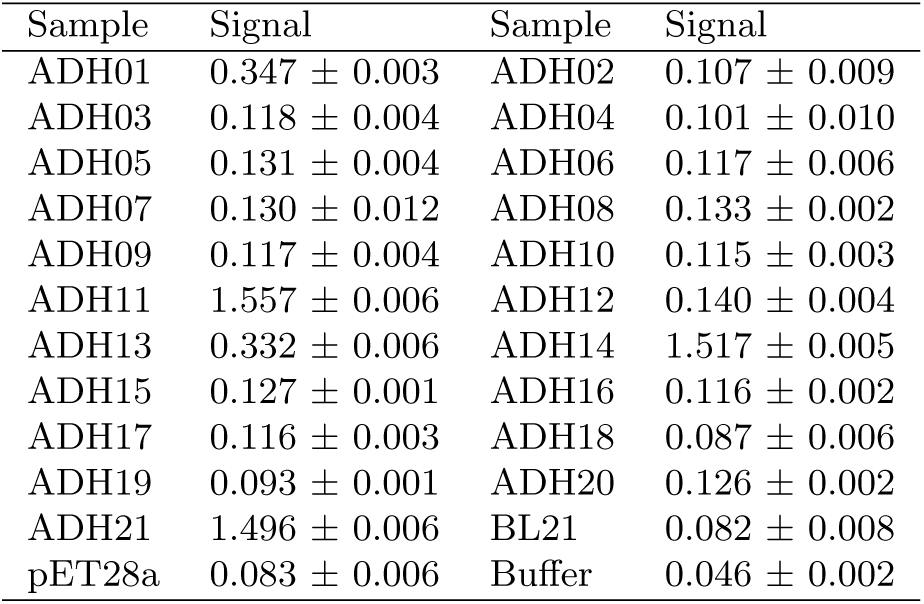
ADH crude-lysate activity screen with 10 mM ethanol. Values are mean absorbance signals at 450 nm *±* s.d. from three independent crude-lysate assays.

**Table 13.** MDH crude-lysate activity screen with 2 mM L-malate. Values are mean absorbance signals at 450 nm *±* s.d. from three independent crude-lysate assays.

| Sample | Signal | Sample | Signal |
| --- | --- | --- | --- |
| MDH01 | 0.360 ± 0.008 | MDH02 | 0.362 ± 0.006 |
| MDH03 | 0.367 ± 0.014 | MDH04 | 0.354 ± 0.002 |
| MDH05 | 0.385 ± 0.006 | MDH06 | 0.355 ± 0.005 |
| MDH07 | 0.379 ± 0.007 | MDH08 | 0.351 ± 0.010 |
| MDH09 | 0.350 ± 0.004 | MDH10 | 0.390 ± 0.007 |
| MDH11 | 0.300 ± 0.004 | MDH12 | 0.341 ± 0.009 |
| MDH13 | 0.347 ± 0.007 | MDH14 | 0.353 ± 0.013 |
| MDH15 | 0.347 ± 0.006 | MDH16 | 0.341 ± 0.002 |
| MDH17 | 0.355 ± 0.009 | MDH18 | 0.355 ± 0.007 |
| MDH19 | 0.413 ± 0.004 | MDH20 | 0.355 ± 0.003 |
| MDH21 | 0.456 ± 0.017 | MDH22 | 0.436 ± 0.021 |
| BL21 | 0.205 ± 0.011 | pET28a | 0.215 ± 0.007 |
| Buffer | 0.045 ± 0.001 | | |

### 7.2 MMseqs2 natural-sequence similarity statistics

At the sequence-entry level, 2,727 of 3,424 ZymCTRL sequences yield a top1 hit in UniRef50, corresponding to a hit rate of 79.64%, whereas 4,585 of 4,736 EnzymeArt sequences yield a top1 hit, corresponding to a hit rate of 96.81% (Table 15). The top1 sequence-identity distribution is likewise shifted upward for EnzymeArt, with a median pident of 59.7% compared with 48.0% for ZymCTRL and a mean of 61.31% compared with 52.86%. Similarly, the fraction of top1 matches with pident ≥ 50% is 71.10% for EnzymeArt versus 46.75% for ZymCTRL, and the fraction with pident ≥ 70% is 31.04% versus 22.08%. These results indicate that EnzymeArt sequences are, overall, positioned closer to natural sequence neighbourhoods than ZymCTRL sequences. This pattern, however, does not extend uniformly to all alignment statistics. Although EnzymeArt shows higher top1 identity, its median top1 bit score is lower than that of ZymCTRL (244 versus 286), and this difference is accompanied by a pronounced asymmetry in target coverage: the median qcov is similarly high for both methods (0.969 for EnzymeArt and 0.961 for ZymCTRL), whereas the median tcov is substantially lower for EnzymeArt (0.635 versus 0.917). These results suggest that many EnzymeArt hits align strongly to most of the generated query but only to a more limited fraction of a longer natural target. In other words, the higher apparent naturalness of EnzymeArt does not necessarily imply greater full-length similarity to natural proteins. Rather, it is often consistent with stronger local similarity within conserved regions, domain-like segments, or functionally constrained cores of longer natural proteins. By contrast, when ZymCTRL sequences recover natural neighbours, the alignments more often extend across a larger fraction of both the query and the target, consistent with broader full-length homology. The same pattern appears in the all-hit landscape: under the same –-max-seqs 300 cutoff, EnzymeArt yields 1,333,048 retained hits, compared with 689,055 for ZymCTRL, indicating that EnzymeArt sequences tend to occupy denser natural sequence neighbourhoods with more retrievable nearby homologues, whereas ZymCTRL sequences are distributed more broadly toward lower-similarity regions of sequence space. A technical caveat applies to the EnzymeArt query file: although enzymeart common ec.fasta contains 4,736 sequence entries, it includes only 926 unique FASTA headers, with all 926 headers duplicated and a maximum duplication count of 20, whereas all 3,424 headers in zymctrl common ec.fasta are unique. To avoid artefacts introduced by header-level aggregation, all formal comparisons are therefore performed at the sequence-entry level rather than the unique-header level.

**Table 14.** Lipase crude-lysate activity screens with tributyrin and triolein. Values are mean absorbance signals at 570 nm *±* s.d. from three independent crude-lysate assays.

| Sample | Tributyrin | Triolein |
| --- | --- | --- |
| LP01 | 0.321 ± 0.007 | 0.173 ± 0.003 |
| LP02 | 0.312 ± 0.005 | 0.163 ± 0.004 |
| LP03 | 0.321 ± 0.005 | 0.171 ± 0.004 |
| LP04 | 0.345 ± 0.004 | 0.173 ± 0.006 |
| LP05 | 0.324 ± 0.003 | 0.172 ± 0.001 |
| LP06 | 0.322 ± 0.005 | 0.175 ± 0.005 |
| LP07 | 0.345 ± 0.005 | 0.179 ± 0.005 |
| LP08 | 0.317 ± 0.008 | 0.158 ± 0.003 |
| LP09 | 0.346 ± 0.004 | 0.179 ± 0.003 |
| LP10 | 0.328 ± 0.006 | 0.193 ± 0.002 |
| LP11 | 0.345 ± 0.009 | 0.187 ± 0.002 |
| LP12 | 0.362 ± 0.008 | 0.182 ± 0.001 |
| LP13 | 0.342 ± 0.009 | 0.173 ± 0.001 |
| LP14 | 0.343 ± 0.007 | 0.181 ± 0.005 |
| LP15 | 0.457 ± 0.006 | 0.198 ± 0.007 |
| LP16 | 0.388 ± 0.014 | 0.170 ± 0.005 |
| LP17 | 0.354 ± 0.009 | 0.182 ± 0.006 |
| LP18 | 0.363 ± 0.007 | 0.177 ± 0.003 |
| LP19 | 0.379 ± 0.004 | 0.166 ± 0.003 |
| LP20 | 0.363 ± 0.010 | 0.161 ± 0.003 |
| P61872 | 0.415 ± 0.007 | 0.189 ± 0.005 |
| Commercial lipase | 0.458 ± 0.011 | 0.207 ± 0.005 |
| BL21 | 0.061 ± 0.003 | 0.057 ± 0.006 |
| pET28a | 0.092 ± 0.005 | 0.073 ± 0.004 |
| Blank | 0.047 ± 0.002 | 0.046 ± 0.002 |

**Table 15.** Top1 similarity statistics from MMseqs2 searches against UniRef50.

| Metric | ZymCTRL | EnzymeArt |
| --- | --- | --- |
| Total sequences | 3,424 | 4,736 |
| Top1 hits | 2,727 | 4,585 |
| No-hit sequences | 697 | 151 |
| Hit rate (%) | 79.64 | 96.81 |
| Top1 **pident**, mean (%) | 52.86 | 61.31 |
| Top1 **pident**, median (%) | 48.0 | 59.7 |
| Top1 **bits**, mean | 342.96 | 249.63 |
| Top1 **bits**, median | 286 | 244 |
| Top1 **qcov**, median | 0.961 | 0.969 |
| Top1 **tcov**, median | 0.917 | 0.635 |
| Top1 **pident** ≥ 50% (%) | 46.75 | 71.10 |
| Top1 **pident** ≥ 70% (%) | 22.08 | 31.04 |
| Top1 **pident** ≥ 90% (%) | 6.60 | 4.99 |
| All retained hits | 689,055 | 1,333,048 |

Overall, these analyses show that ZymCTRL sequences are globally less similar to natural proteins than EnzymeArt sequences, as reflected by their lower UniRef50 hit rate, lower top1 sequence identity, lower fraction of high-identity matches, and smaller all-hit neighbourhood under the same search threshold. EnzymeArt therefore appears closer to natural sequence space on average, but primarily through stronger local similarity to conserved regions of natural proteins rather than uniformly greater similarity across the full protein length. ZymCTRL, by contrast, explores a more divergent region of sequence space overall; however, when it does recover natural neighbours, it more often does so at the level of broader full-length similarity. Sequence identity alone is therefore insufficient to characterize proximity to natural sequence space, and needs to be interpreted jointly with bit score and alignment coverage.

### 7.3 Subclass-wise structural confidence statistics

**Table 16:**
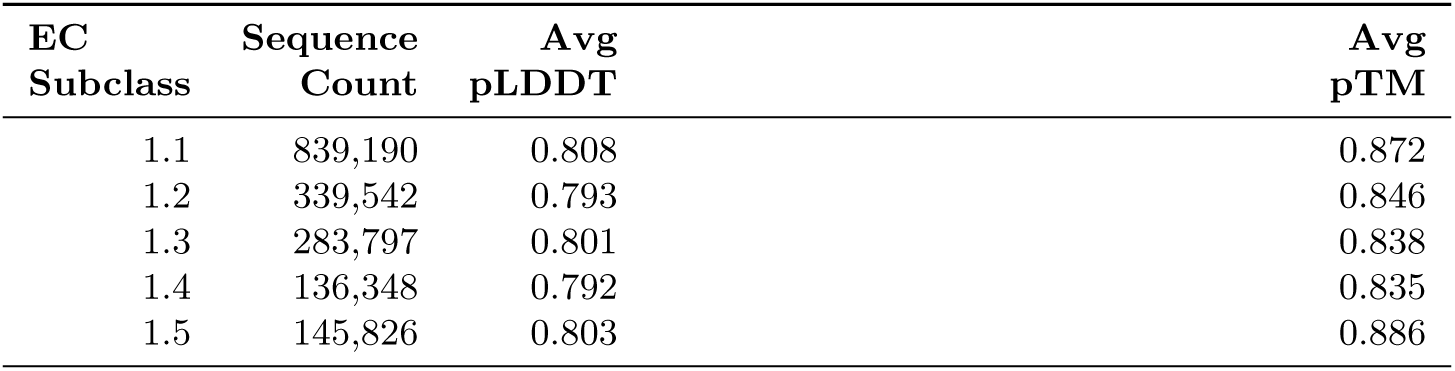
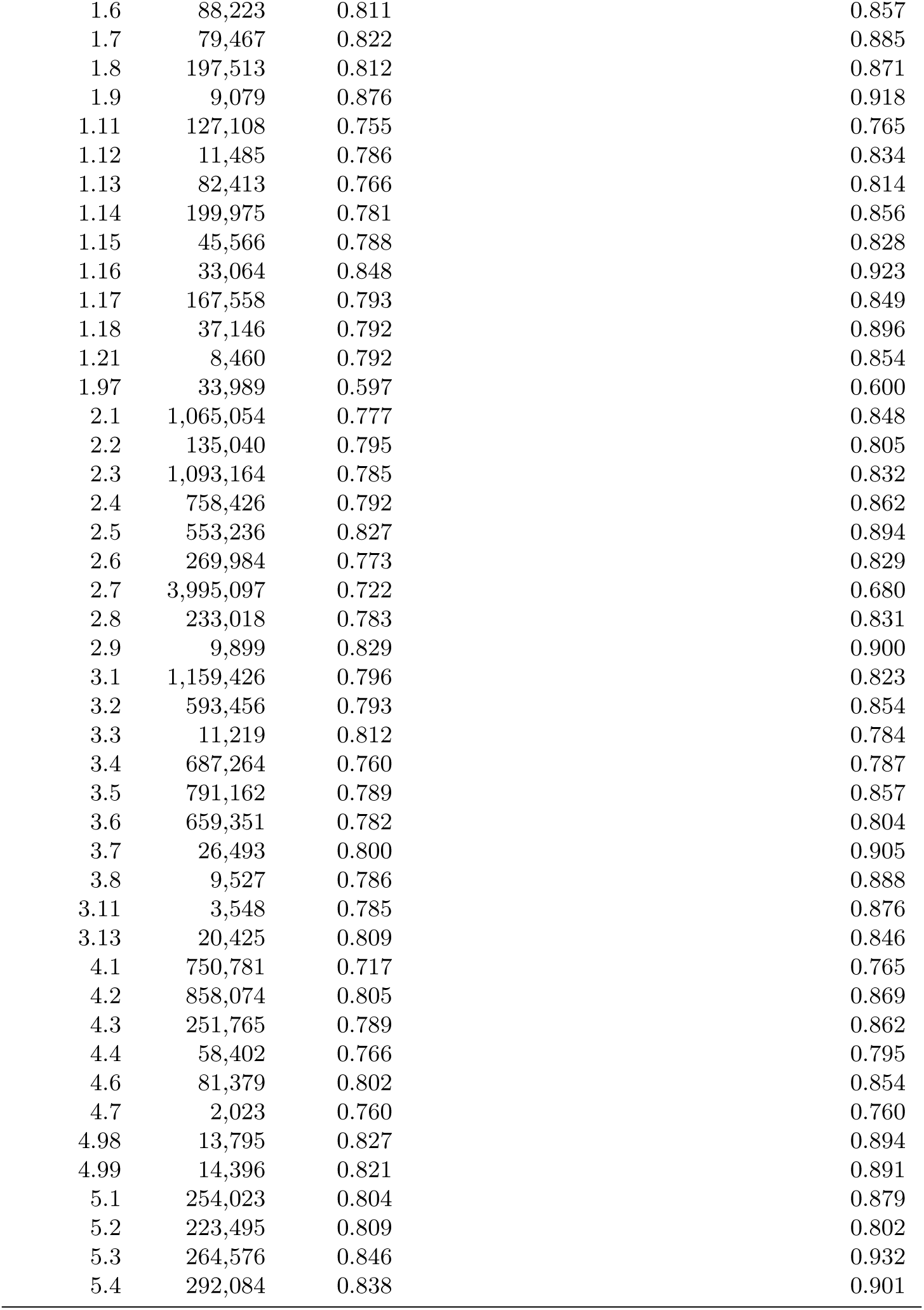
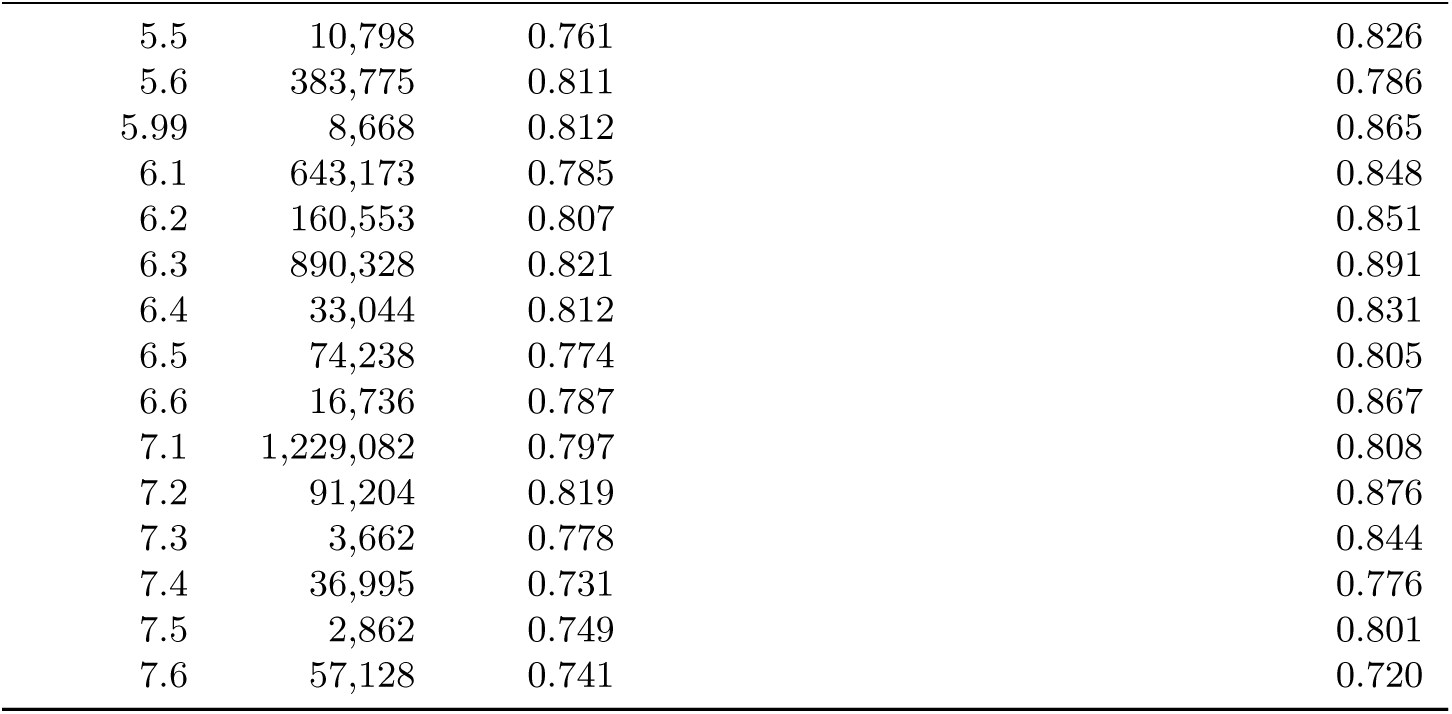
Subclass-wise statistics of predicted structural confidence.

### 7.4 Supplementary analysis of the prediction–evaluation–redesign loop

To quantify the effect of iterative refinement in EnzymeArt, we evaluate sequences across successive redesign stages using predicted structural confidence and sequence–function alignment metrics. For each model, we analyze three stages: the initial generated sequences, sequences after one redesign cycle, and sequences after a second redesign cycle (Tab. 17).

**Table 17.** Structural and semantic evaluation metrics across redesign loop for different models.

| Model | Stage | Sequences | pLDDT | PAE | pTM | ProTrek Score |
| --- | --- | --- | --- | --- | --- | --- |
| EnzymeArt | Initial | 20,000 | 0.749 | 8.847 | 0.787 | 0.329 |
|  | Round 1 | 20,000 | 0.766 | 8.463 | 0.796 | 0.326 |
|  | Round 2 | 20,000 | 0.772 | 8.368 | 0.798 | 0.324 |
| EnzymeArt-base | Initial | 20,000 | 0.688 | 11.117 | 0.656 | 0.343 |
|  | Round 1 | 20,000 | 0.721 | 10.168 | 0.687 | 0.338 |
|  | Round 2 | 20,000 | 0.737 | 9.714 | 0.701 | 0.335 |
| Pinal | Initial | 1,000 | 0.688 | 10.283 | 0.687 | 0.320 |
|  | Round 1 | 1,000 | 0.700 | 10.233 | 0.686 | 0.314 |
|  | Round 2 | 1,000 | 0.707 | 10.249 | 0.686 | 0.309 |
| zymCTRL | Initial | 2,800 | 0.732 | 9.234 | 0.772 | 0.309 |
|  | Round 1 | 2,800 | 0.746 | 8.891 | 0.781 | 0.307 |
|  | Round 2 | 2,800 | 0.754 | 8.723 | 0.785 | 0.305 |

#### Improvement of predicted structural confidence

Iterative redesign consistently increases predicted structural confidence. For EnzymeArt, mean pLDDT increases from 0.749 to 0.772 over two refinement rounds, whereas PAE decreases from 8.847 to 8.368 and pTM increases from 0.787 to 0.798. EnzymeArt-base shows a similar trend, with pLDDT increasing from 0.688 to 0.737 and PAE decreasing from 11.117 to 9.714.

Baseline generators exhibit comparable but smaller improvements. For Pinal, pLDDT increases modestly from 0.688 to 0.707, while PAE remains largely stable. For ZymCTRL, pLDDT increases from 0.732 to 0.754 and PAE decreases from 9.234 to 8.723. These results indicate that the redesign procedure consistently improves predicted structural confidence across diverse sequence generators.

#### Trade-off between structural confidence and functional alignment

In contrast, sequence–function alignment decreases slightly during redesign. For EnzymeArt, ProTrek similarity decreases from 0.329 to 0.324 after two refinement rounds. A similar gradual reduction is observed for EnzymeArt-base (0.343 to 0.335), Pinal (0.320 to 0.309), and ZymCTRL (0.309 to 0.305).

This trend reflects the objective of the redesign step, which prioritizes structural plausibility rather than explicitly optimizing functional prompt alignment. Mutations introduced to resolve low-confidence regions can therefore improve predicted foldability while slightly weakening the original conditioning signal.

Because sequence sets differ across models in conditioning inputs and sample sizes, the absolute metric values are not intended for direct cross-model comparison. Instead, these analyses show that the redesign module produces consistent directional effects, increasing structural confidence while introducing a modest reduction in function alignment.

### 7.5 Ablation: Necessity of EnzymeArt-base Pretraining

To evaluate the contribution of the EnzymeArt-base pretraining stage to enzyme-centric generation, we perform an ablation in which the model is trained without general protein–function pretraining and is instead pretrained only on enzyme sequences. We compare three models—EnzymeArt, a 500K-step enzyme-only variant and a 1M-step enzyme-only variant—using pLDDT, PAE and pTM, and ProTrek-score as a proxy for text–sequence alignment.

As shown in Fig. 9, EnzymeArt has the highest mean pLDDT (0.78) and pTM (0.82) and the lowest mean PAE (6.58). As enzyme-only pretraining increases from 500K to 1M steps, mean pLDDT decreases (0.77 to 0.75), mean PAE increases (7.25 to 8.72), and mean pTM decreases (0.80 to 0.75). ProTrek-score also decreases from 0.31 (EnzymeArt) to 0.28 (enzyme-only variants).

**Fig. 9.**
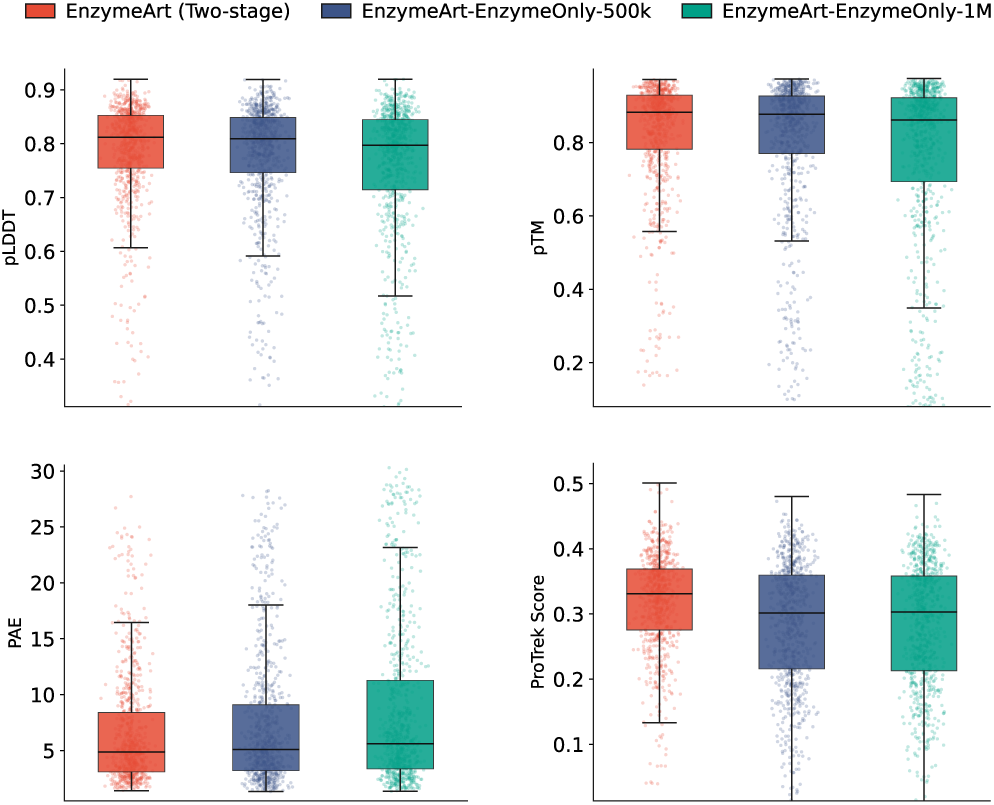
Comparison of two-stage and enzyme-only pretraining strategies.

The distributions show the same pattern. Median pLDDT is similar for EnzymeArt and the 500K enzyme-only model (0.81 vs 0.80), but variability increases (pLDDT s.d. 0.09 for EnzymeArt, 0.11 for 500K and 0.13 for 1M). Median PAE increases from 4.87 (EnzymeArt) to 5.10 (500K) and 5.62 (1M). Mean and median pTM also decrease in the enzyme-only variants. Together, these results are consistent with EnzymeArt-base pretraining contributing to higher structural confidence and better text-conditioned alignment in downstream enzyme-centric design.

**Table 18.** Unconditional generation benchmark across target sequence lengths. pLDDT and pTM are reported as predicted structural-confidence metrics and are used only for computational comparison.

| Length | EnzymeArt-base |  | RFdiffusion |  | Chroma |  |
| --- | --- | --- | --- | --- | --- | --- |
|  | pLDDT | pTM | pLDDT | pTM | pLDDT | pTM |
| 100 | 0.6867 | 0.6295 | 0.7591 | 0.8269 | 0.6500 | 0.6100 |
| 200 | 0.8083 | 0.7830 | 0.6927 | 0.7768 | 0.6100 | 0.6100 |
| 300 | 0.7901 | 0.7353 | 0.6186 | 0.6761 | 0.5900 | 0.5800 |
| 400 | 0.7934 | 0.7576 | 0.5520 | 0.5495 | 0.5700 | 0.5500 |
| 500 | 0.8082 | 0.7715 | 0.5147 | 0.4861 | 0.5400 | 0.4700 |
| 600 | 0.7994 | 0.7002 | 0.4519 | 0.3994 | 0.5200 | 0.4400 |
| 700 | 0.7778 | 0.6646 | 0.4254 | 0.3567 | 0.5300 | 0.4100 |
| 800 | 0.7854 | 0.6260 | 0.3857 | 0.3078 | 0.5400 | 0.4000 |
| 900 | 0.7780 | 0.6124 | 0.3625 | 0.2688 | 0.5500 | 0.3900 |
| 1000 | 0.7445 | 0.5275 | 0.3431 | 0.2463 | 0.5400 | 0.3700 |

### 7.6 Unconditional sequence-generation benchmark across target lengths

We evaluated unconditional sequence generation across target sequence lengths as a supplementary computational benchmark.

### 7.7 Comparison with function-guided protein design methods

**Table 19.**
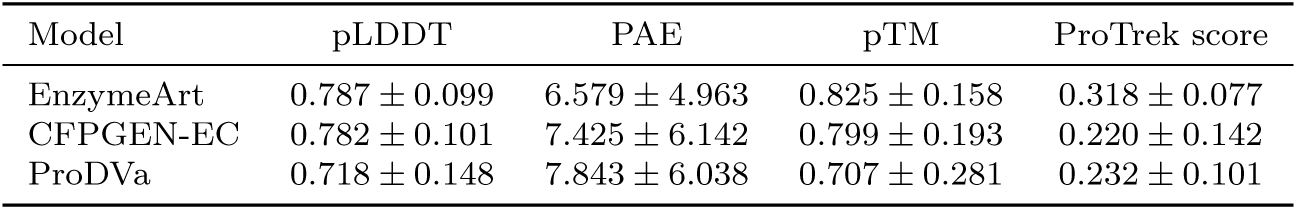
Structural and semantic evaluation metrics of different enzyme design models. Values are shown as mean *±* s.d.

| Model | pLDDT | PAE | pTM | ProTrek score |
| --- | --- | --- | --- | --- |
| EnzymeArt | 0.787 ± 0.099 | 6.579 ± 4.963 | 0.825 ± 0.158 | 0.318 ± 0.077 |
| CFPGEN-EC | 0.782 ± 0.101 | 7.425 ± 6.142 | 0.799 ± 0.193 | 0.220 ± 0.142 |
| ProDVa | 0.718 ± 0.148 | 7.843 ± 6.038 | 0.707 ± 0.281 | 0.232 ± 0.101 |

We compare three function-guided protein design methods (EnzymeArt, CFPGEN-EC [49] and ProDVa [50]). EnzymeArt shows the strongest overall performance, with higher structural confidence and fold consistency (mean pLDDT 0.78; mean pTM 0.82) and lower error (mean PAE 6.58), together with the highest ProTrek score (0.31).

**Fig. 10.**
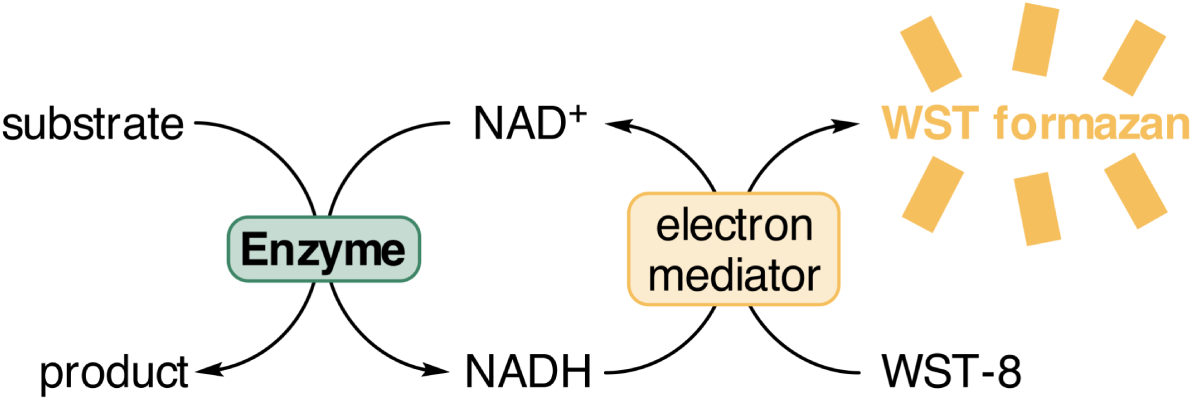
Principle of the NAD^+^-dependent enzyme activity assay using WST-8. NAD^+^-dependent enzymes catalyse the target reaction with concomitant reduction of NAD^+^ to NADH. The generated NADH then reduces WST-8 to orange formazan in the presence of an electron mediator. The resulting formazan product shows strong absorbance at approximately 450 nm. Therefore, the amount of formazan formed is directly proportional to NADH production and is used as a readout of enzymatic activity.

**Fig. 11.**
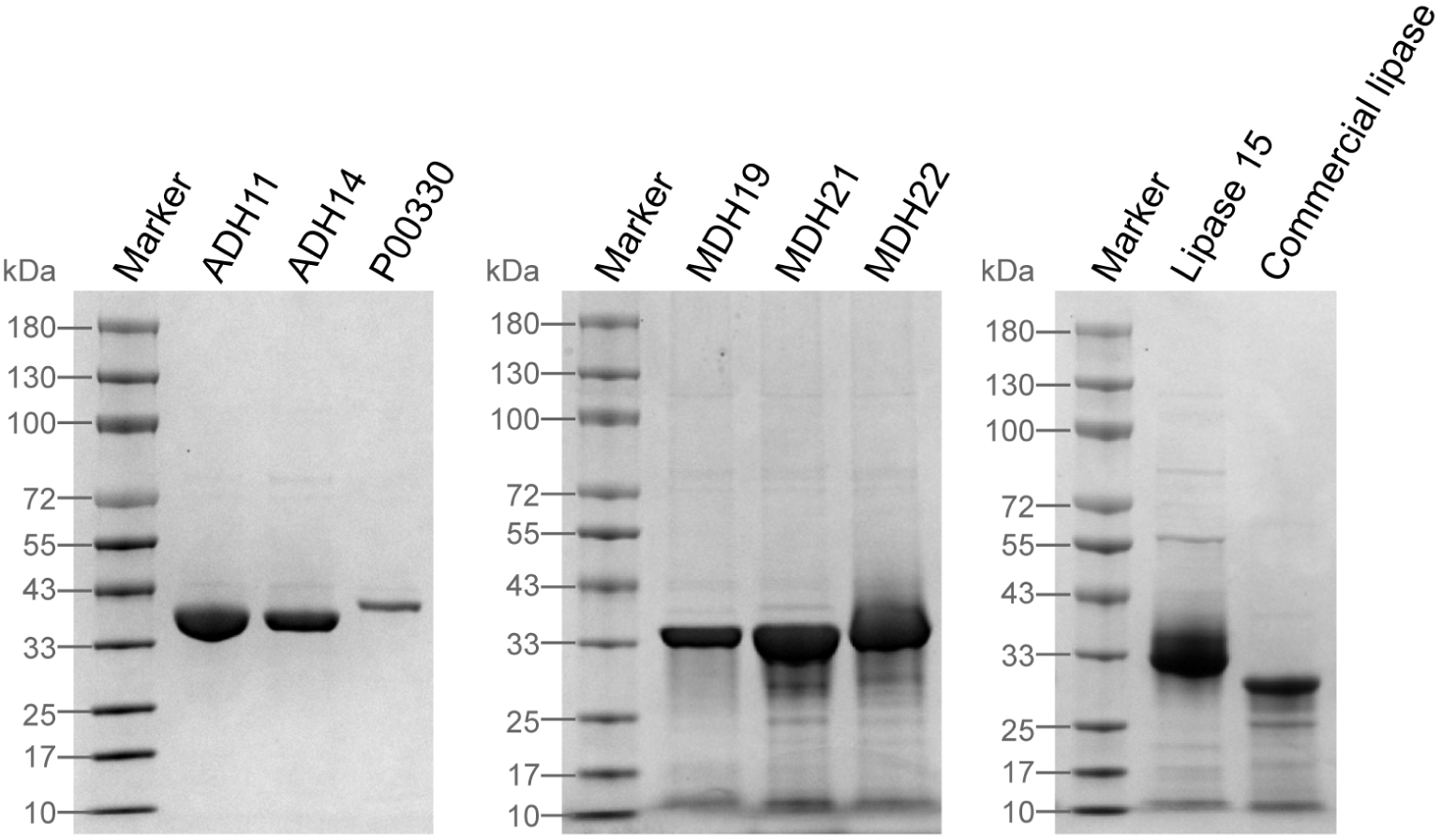
SDS gel electrophoresis of selected enzymes.

**Fig. 12.**
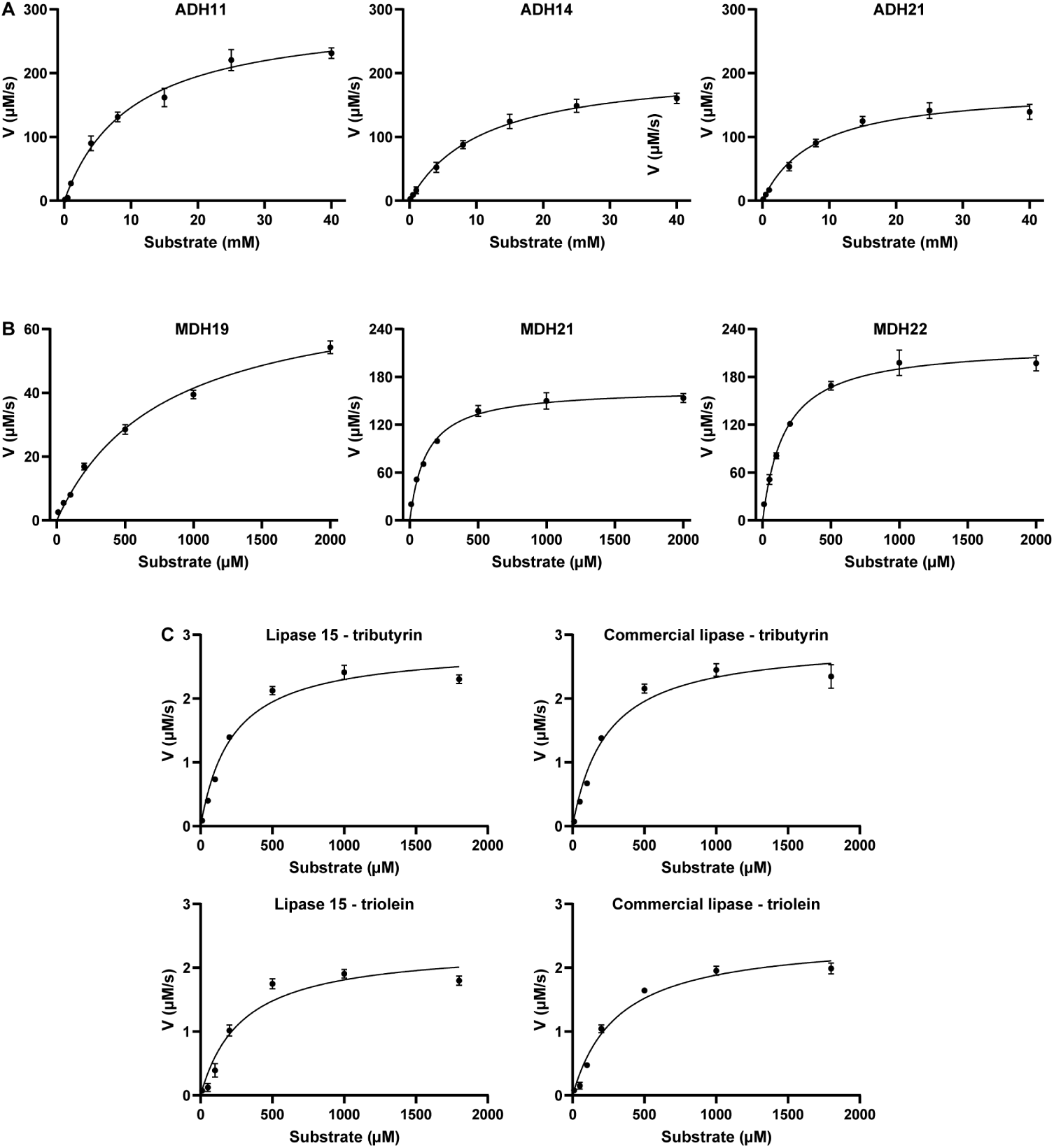
Michaelis–Menten kinetics of selected purified enzymes.

**Table 20.** Software tools and versions used in this study.

| Software | Version | Primary Use | URL |
| --- | --- | --- | --- |
| AlphaFold3 | v3.0.1 | Structure prediction | https://github.com/google-deepmind/alphafold3 |
| ESM C | – | Protein LM for perplexity evaluation | https://github.com/evolutionaryscale/esm |
| ESM-IF | – | Structure-informed sequence scoring | https://github.com/facebookresearch/esm |
| ESM-2 | – | Protein representation learning | https://github.com/facebookresearch/esm |
| ESMFold | – | Fast structure prediction from sequence | https://github.com/facebookresearch/esm |
| ProTrek | – | Sequence–function similarity score | https://github.com/westlake-repl/ProTrek |
| BLASTP | – | Sequence similarity search | https://blast.ncbi.nlm.nih.gov/Blast.cgi |
| InterPro | v104.0 | Domain annotation and EC assignment | https://www.ebi.ac.uk/interpro/ |
| PyMOL | v3.1.6.1 | Structural visualization | https://pymol.org/ |

### 7.8 Biochemical robustness of designed ADH enzymes

Because computational confidence scores do not directly establish biochemical stability, we experimentally examined the environmental robustness of representative ADH designs. Two EnzymeArt-generated alcohol dehydrogenases, ADH11 and ADH14, were compared with the natural reference enzyme ADH21 across pH, temperature and organic solvent perturbations.

**Table 21.** Summary of protein molecular weights.

| Protein | Molecular weight (kDa) |
| --- | --- |
| ADH11 | 38.1 |
| ADH14 | 38.1 |
| ADH21 | 39.0 |
| MDH19 | 38.3 |
| MDH21 | 38.8 |
| MDH22 | 38.7 |
| Lipase15 | 29.5 |
| Commercial lipase | 26.5 |

The three enzymes showed similar pH preferences, with maximal activity at pH 8 (Supplementary Fig. 13a). This indicates that the designed enzymes retain a native-like alkaline activity profile rather than showing a shifted or strongly compromised pH optimum. Temperature activity profiling showed an optimal reaction temperature of 35 ^◦^C for ADH11 and 40 ^◦^C for both ADH14 and ADH21 (Supplementary Fig. 13b). Thus, ADH14 exhibited a temperature activity profile comparable to the natural reference, whereas ADH11 remained highly active around moderate temperatures but had a slightly lower apparent temperature optimum.

Finally, organic solvent tolerance assays showed that the designed enzymes remained active under several solvent treatments (Supplementary Fig. 13c). Although all enzymes were strongly inhibited by 50% ethanol, ADH11 retained higher residual activity than ADH21 in 50% DMSO, and ADH14 retained higher residual activity than ADH21 in 80% toluene.

Together, these measurements show that the designed ADHs are not only computationally plausible and catalytically active, but also retain measurable activity under diverse environmental perturbations. These assays do not replace direct biophysical measurements of folding stability, such as melting temperature or unfolding free energy, but they provide experimental evidence that EnzymeArt designs can achieve native-like environmental robustness, with condition-dependent improvements under selected solvent conditions.

**Fig. 13.**
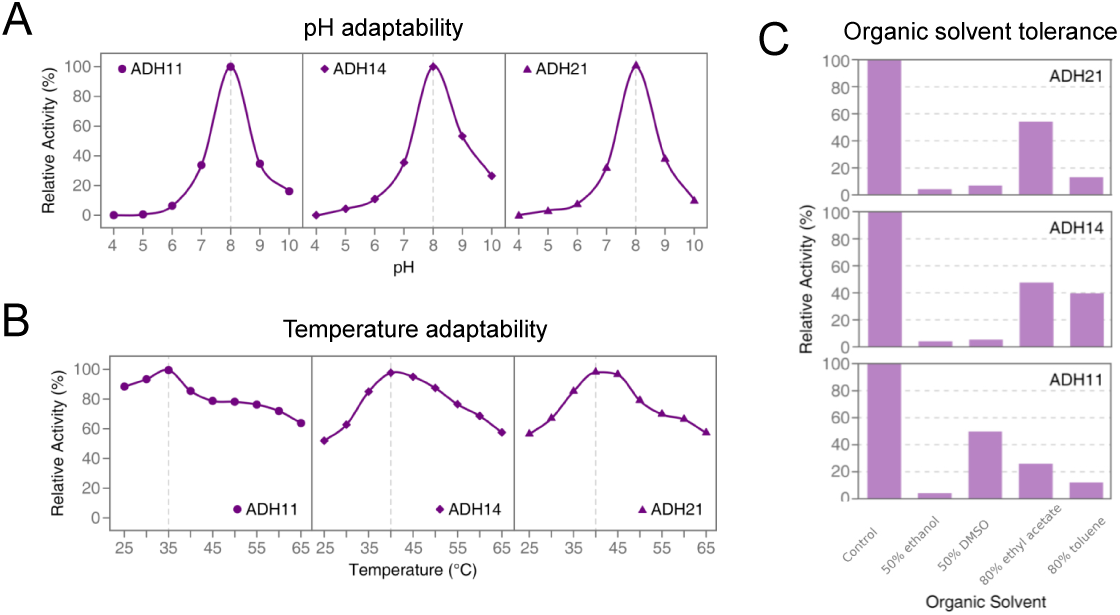
Activity profiles of designed ADHs under pH, temperature and organic solvent perturbations. **A**, pH activity profiles of ADH11, ADH14 and the natural reference ADH21, showing maximal activity at pH 8 for all three enzymes. **B**, Temperature activity profiles, showing optimal reaction temperatures of 35 *^◦^*C for ADH11 and 40 *^◦^*C for ADH14 and ADH21. **C**, Organic solvent tolerance profiles. ADH11 and ADH14 retained measurable activity under several organic solvent treatments and showed condition-dependent advantages relative to ADH21, including higher residual activity of ADH11 in 50% DMSO and higher residual activity of ADH14 in 80% toluene. Activities are shown relative to the corresponding control condition for each enzyme.

### 7.9 Substrate scope evaluation across seven representative alcohols

**Fig. 14.**
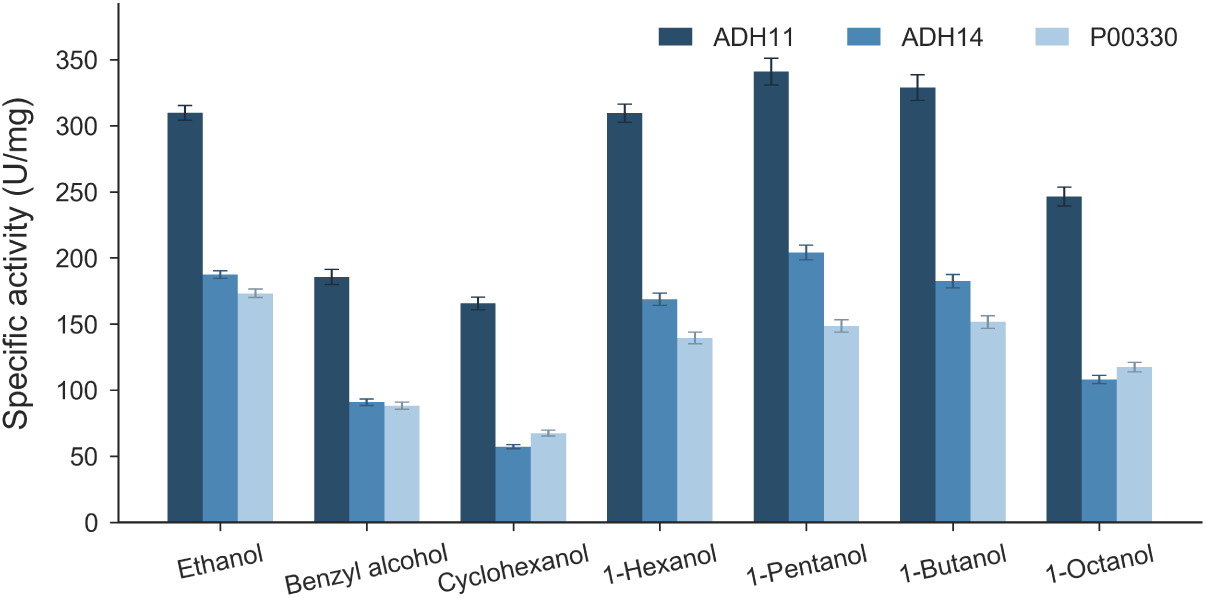
Substrate-specific activities of designed enzymes. Relative activities of ADH11 and ADH14 were measured across a panel of seven alcohol substrates and compared with the natural reference enzyme (P00330). The designed variants exhibit enhanced activity on several substrates, indicating improved catalytic performance and broader substrate tolerance.

### 7.10 Designed ADH sequences generated by EnzymeArt

**Table 22:**
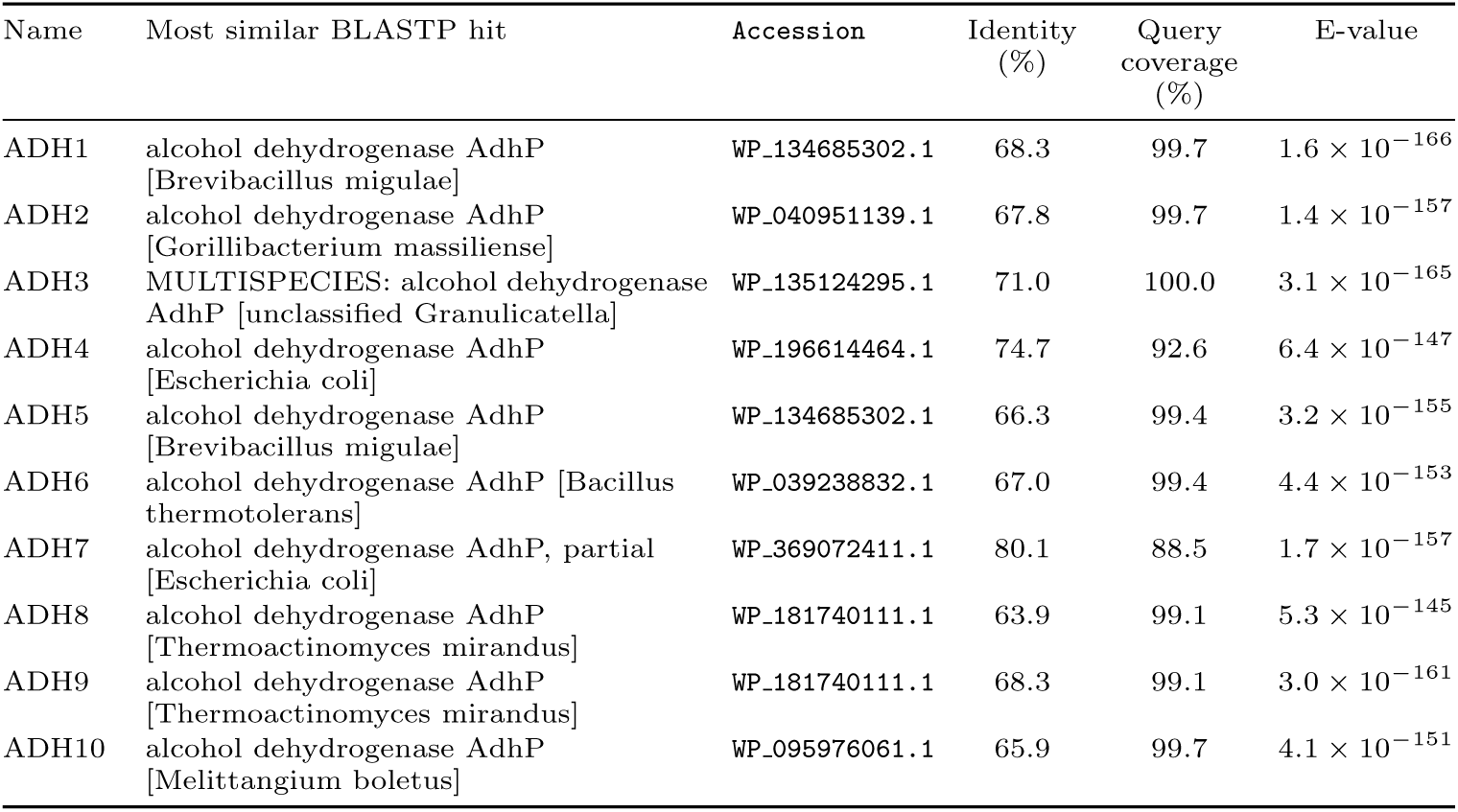
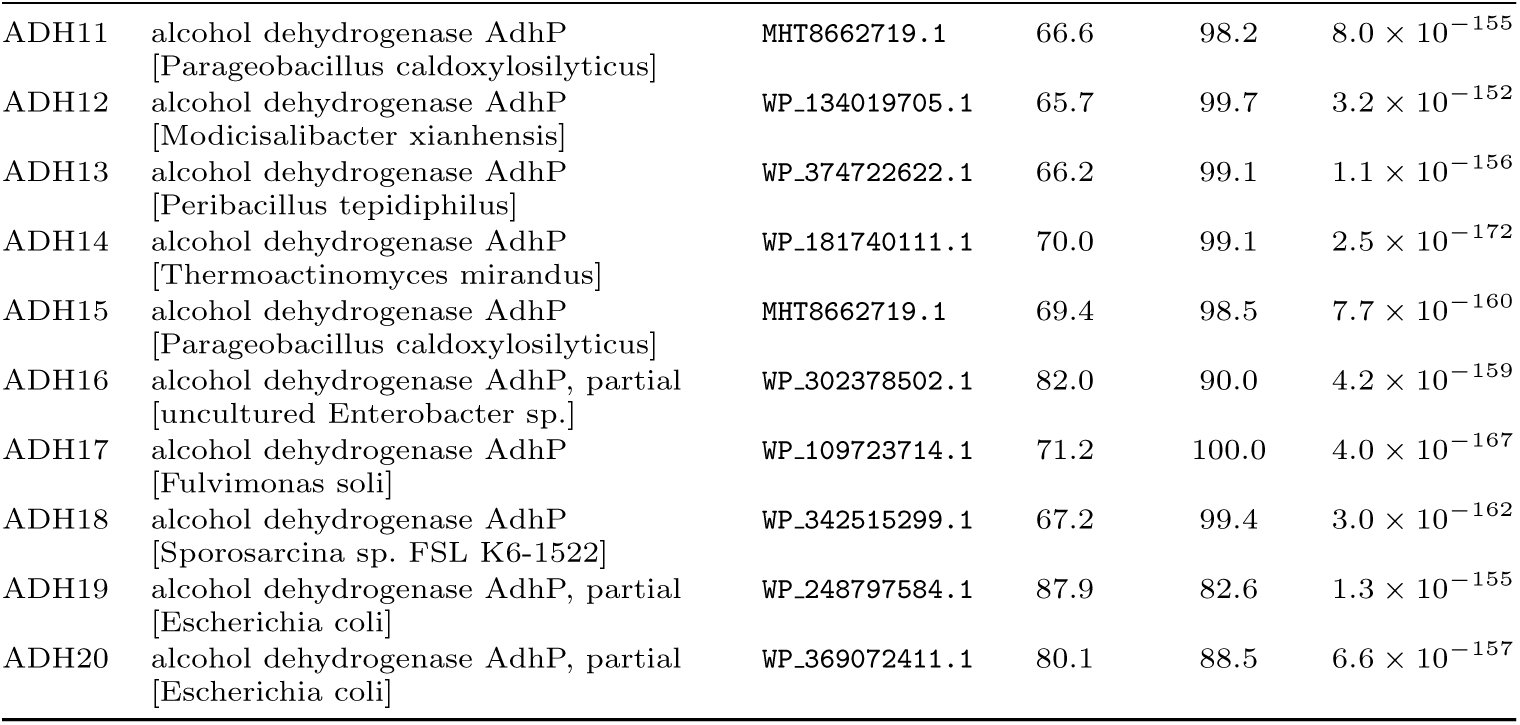
Most similar BLASTP hit for each EnzymeArt-designed alcohol dehydrogenase sequence, selected by maximum percent identity using NCBI web BLASTP results from 26 June 2026. Hit title, accession, identity, query coverage and E-value correspond to the selected hit.

| Name | Most similar BLASTP hit | Accession | Identity (%) | Query coverage (%) | E-value |
| --- | --- | --- | --- | --- | --- |
| ADH1 | alcohol dehydrogenase AdhP [Brevibacillus migulae] | WP_134685302.1 | 68.3 | 99.7 | $1.6 \times 10^{-166}$ |
| ADH2 | alcohol dehydrogenase AdhP [Gorillibacterium massiliense] | WP_040951139.1 | 67.8 | 99.7 | $1.4 \times 10^{-157}$ |
| ADH3 | MULTISPECIES: alcohol dehydrogenase AdhP [unclassified Granulicatella] | WP_135124295.1 | 71.0 | 100.0 | $3.1 \times 10^{-165}$ |
| ADH4 | alcohol dehydrogenase AdhP [Escherichia coli] | WP_196614464.1 | 74.7 | 92.6 | $6.4 \times 10^{-147}$ |
| ADH5 | alcohol dehydrogenase AdhP [Brevibacillus migulae] | WP_134685302.1 | 66.3 | 99.4 | $3.2 \times 10^{-155}$ |
| ADH6 | alcohol dehydrogenase AdhP [Bacillus thermotolerans] | WP_039238832.1 | 67.0 | 99.4 | $4.4 \times 10^{-153}$ |
| ADH7 | alcohol dehydrogenase AdhP, partial [Escherichia coli] | WP_369072411.1 | 80.1 | 88.5 | $1.7 \times 10^{-157}$ |
| ADH8 | alcohol dehydrogenase AdhP [Thermoactinomyces mirandus] | WP_181740111.1 | 63.9 | 99.1 | $5.3 \times 10^{-145}$ |
| ADH9 | alcohol dehydrogenase AdhP [Thermoactinomyces mirandus] | WP_181740111.1 | 68.3 | 99.1 | $3.0 \times 10^{-161}$ |
| ADH10 | alcohol dehydrogenase AdhP [Melittangium boletus] | WP_095976061.1 | 65.9 | 99.7 | $4.1 \times 10^{-151}$ |
| ADH11 | alcohol dehydrogenase AdhP<br>[Parageobacillus caldoxylosilyticus] | MHT8662719.1 | 66.6 | 98.2 | $8.0 \times 10^{-155}$ |
| ADH12 | alcohol dehydrogenase AdhP<br>[Modicisalibacter xianhensis] | WP_134019705.1 | 65.7 | 99.7 | $3.2 \times 10^{-152}$ |
| ADH13 | alcohol dehydrogenase AdhP<br>[Peribacillus tepidophilus] | WP_374722622.1 | 66.2 | 99.1 | $1.1 \times 10^{-156}$ |
| ADH14 | alcohol dehydrogenase AdhP<br>[Thermoactinomyces mirandus] | WP_181740111.1 | 70.0 | 99.1 | $2.5 \times 10^{-172}$ |
| ADH15 | alcohol dehydrogenase AdhP<br>[Parageobacillus caldoxylosilyticus] | MHT8662719.1 | 69.4 | 98.5 | $7.7 \times 10^{-160}$ |
| ADH16 | alcohol dehydrogenase AdhP, partial<br>[uncultured Enterobacter sp.] | WP_302378502.1 | 82.0 | 90.0 | $4.2 \times 10^{-159}$ |
| ADH17 | alcohol dehydrogenase AdhP<br>[Fulvimonas soli] | WP_109723714.1 | 71.2 | 100.0 | $4.0 \times 10^{-167}$ |
| ADH18 | alcohol dehydrogenase AdhP<br>[Sporosarcina sp. FSL K6-1522] | WP_342515299.1 | 67.2 | 99.4 | $3.0 \times 10^{-162}$ |
| ADH19 | alcohol dehydrogenase AdhP, partial<br>[Escherichia coli] | WP_248797584.1 | 87.9 | 82.6 | $1.3 \times 10^{-155}$ |
| ADH20 | alcohol dehydrogenase AdhP, partial<br>[Escherichia coli] | WP_369072411.1 | 80.1 | 88.5 | $6.6 \times 10^{-157}$ |

**Table 23:**
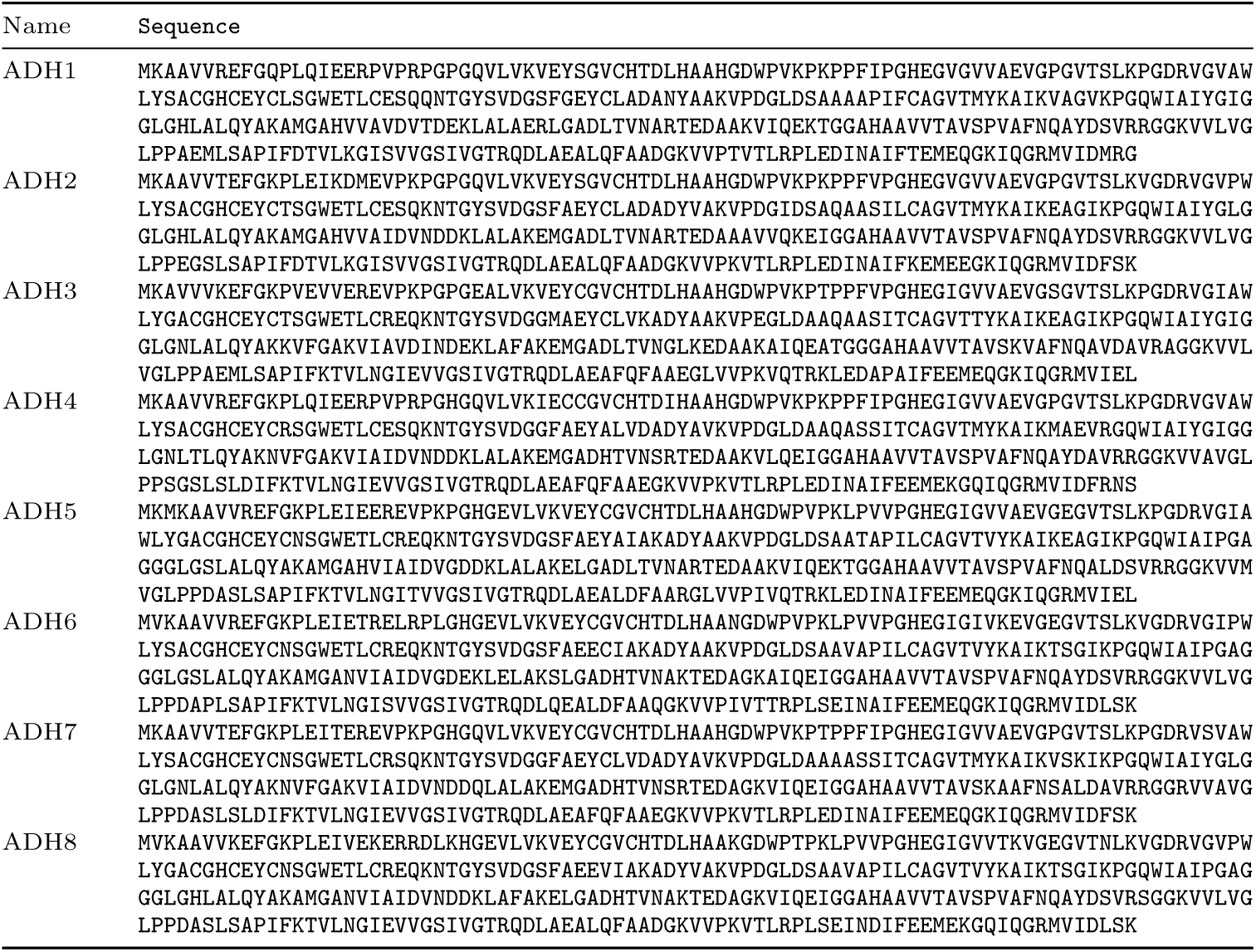
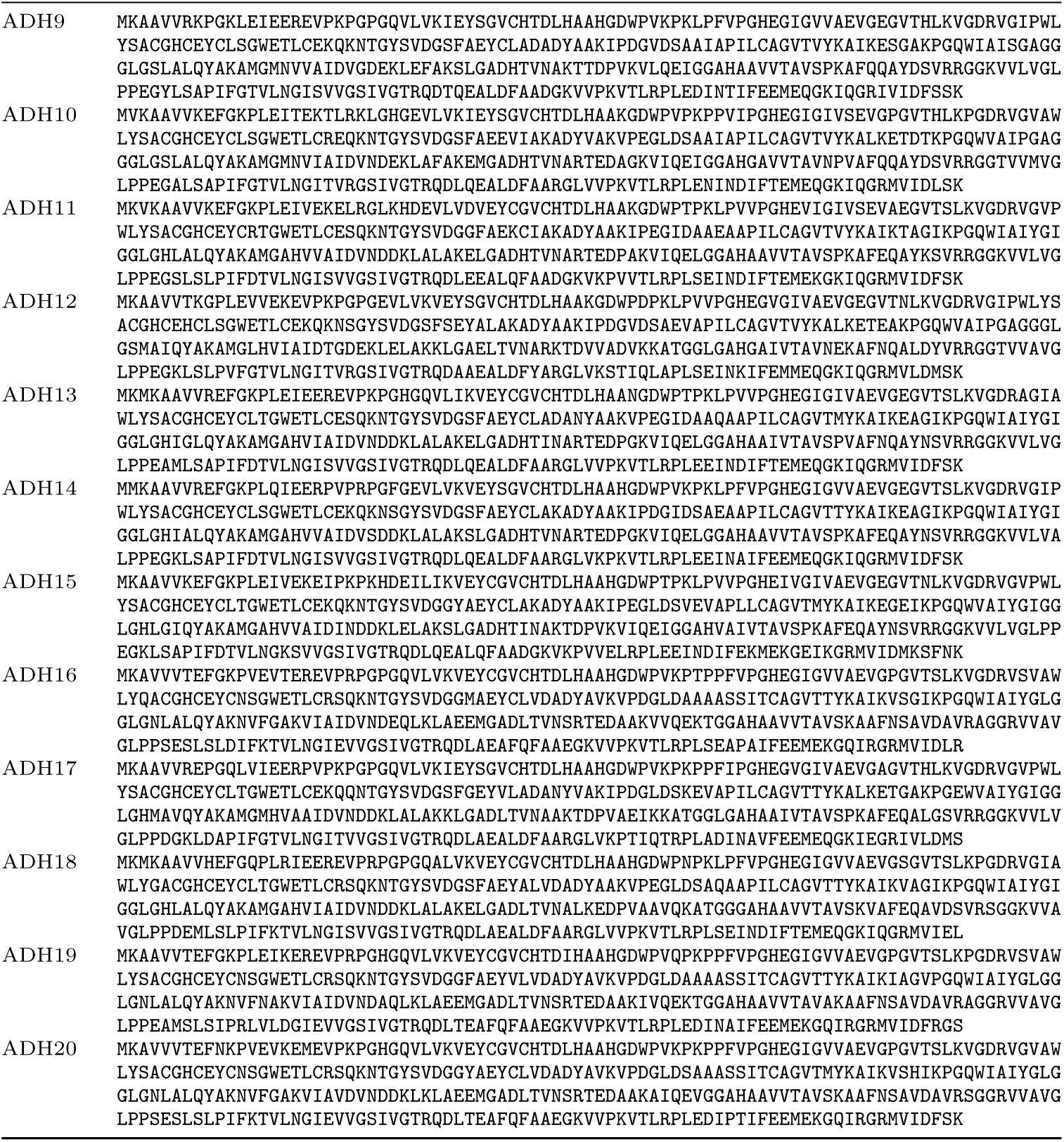
Sequences of EnzymeArt-designed alcohol dehydrogenases.

### 7.11 Designed MDH sequences generated by EnzymeArt

**Table 24:**
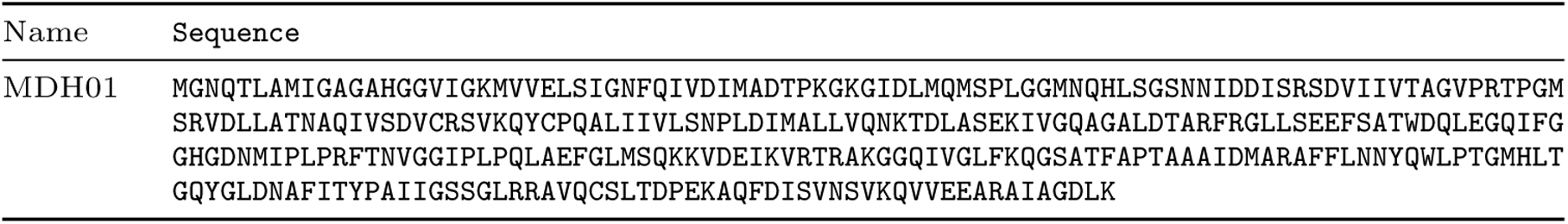
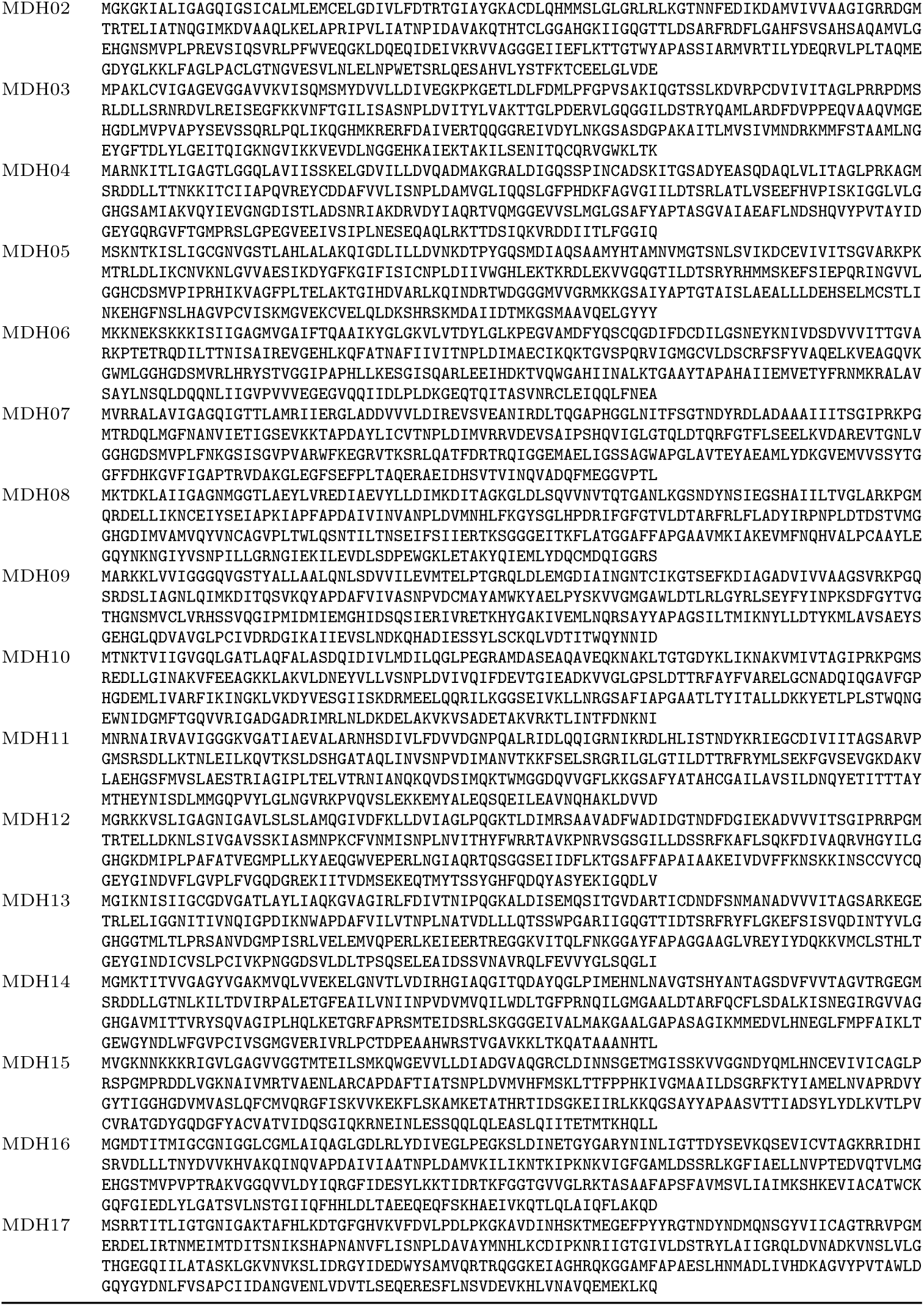
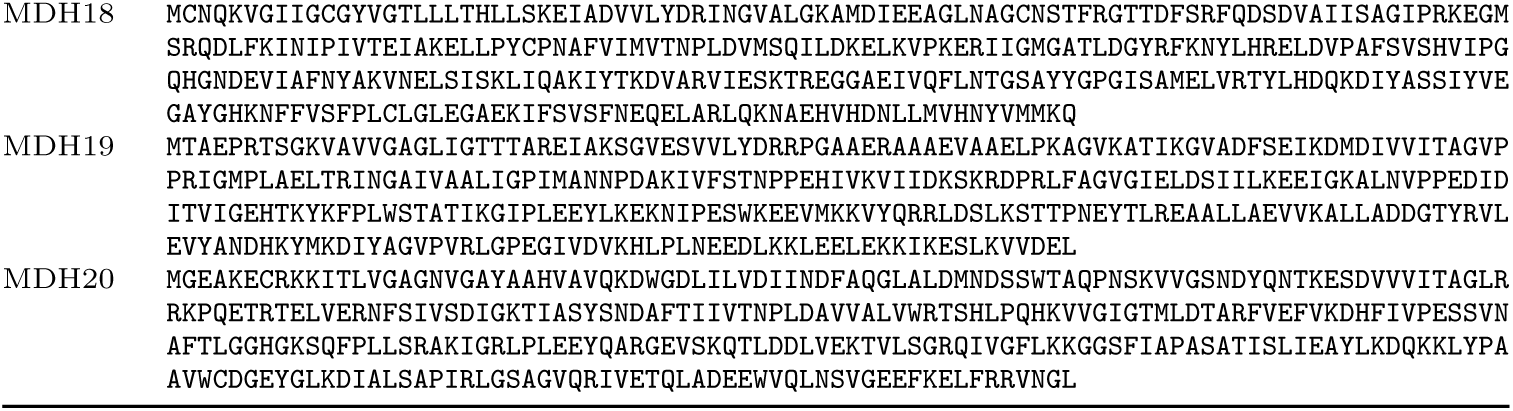
Sequences of EnzymeArt-designed malate dehydrogenases.

**Table 25:**
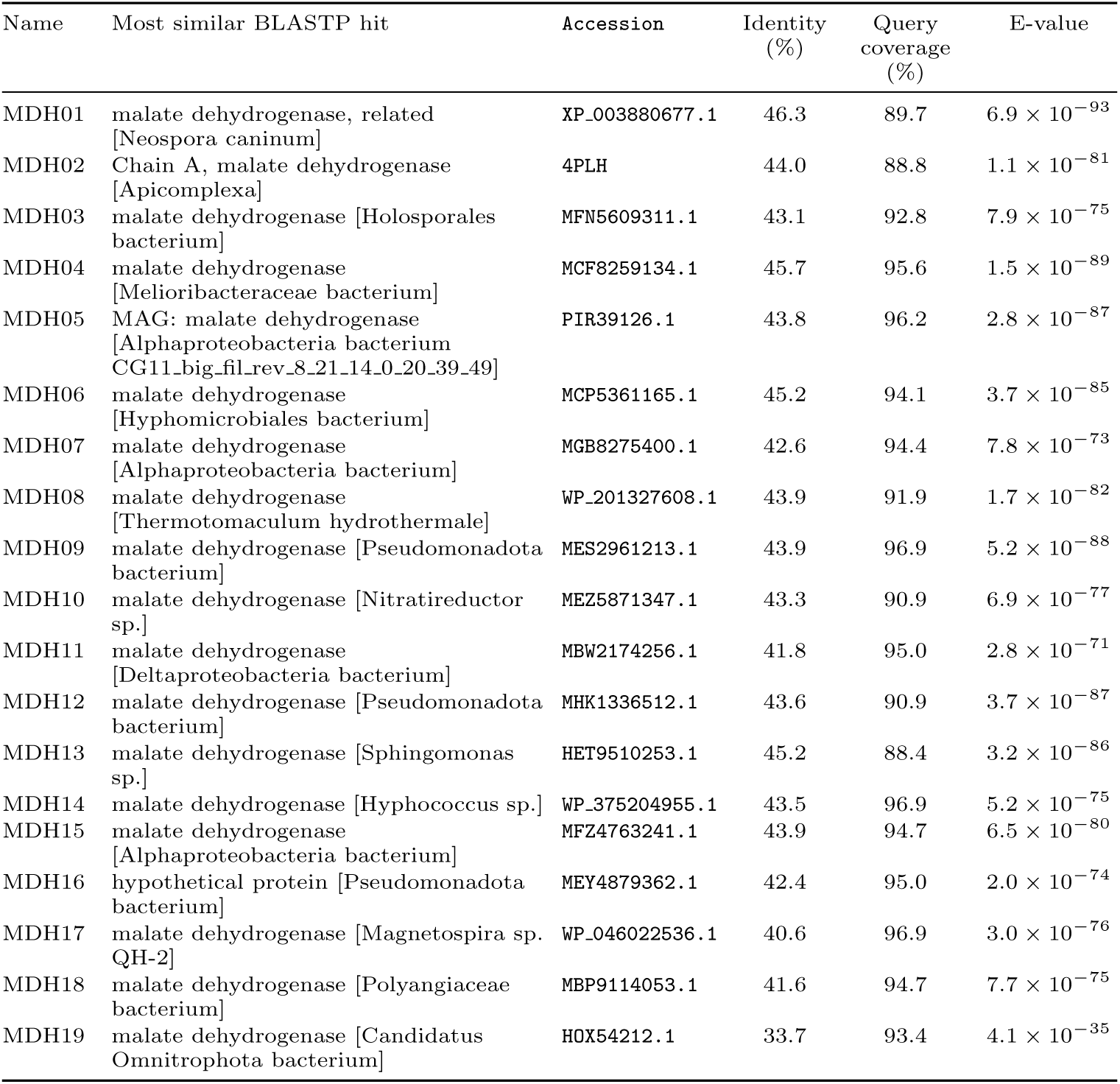
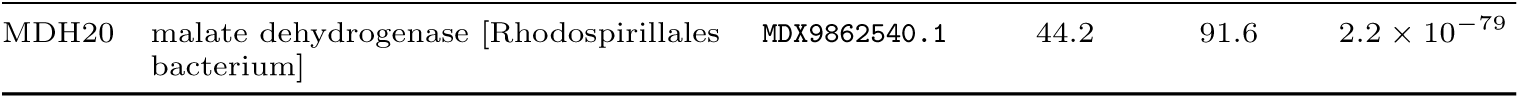
Most similar BLASTP hit for each EnzymeArt-designed malate dehydrogenase sequence, selected by maximum percent identity using NCBI web BLASTP results from 26 June 2026. Hit title, accession, identity, query coverage and E-value correspond to the selected hit.

### 7.12 Similarity of designed lipase sequences

**Table 26:**
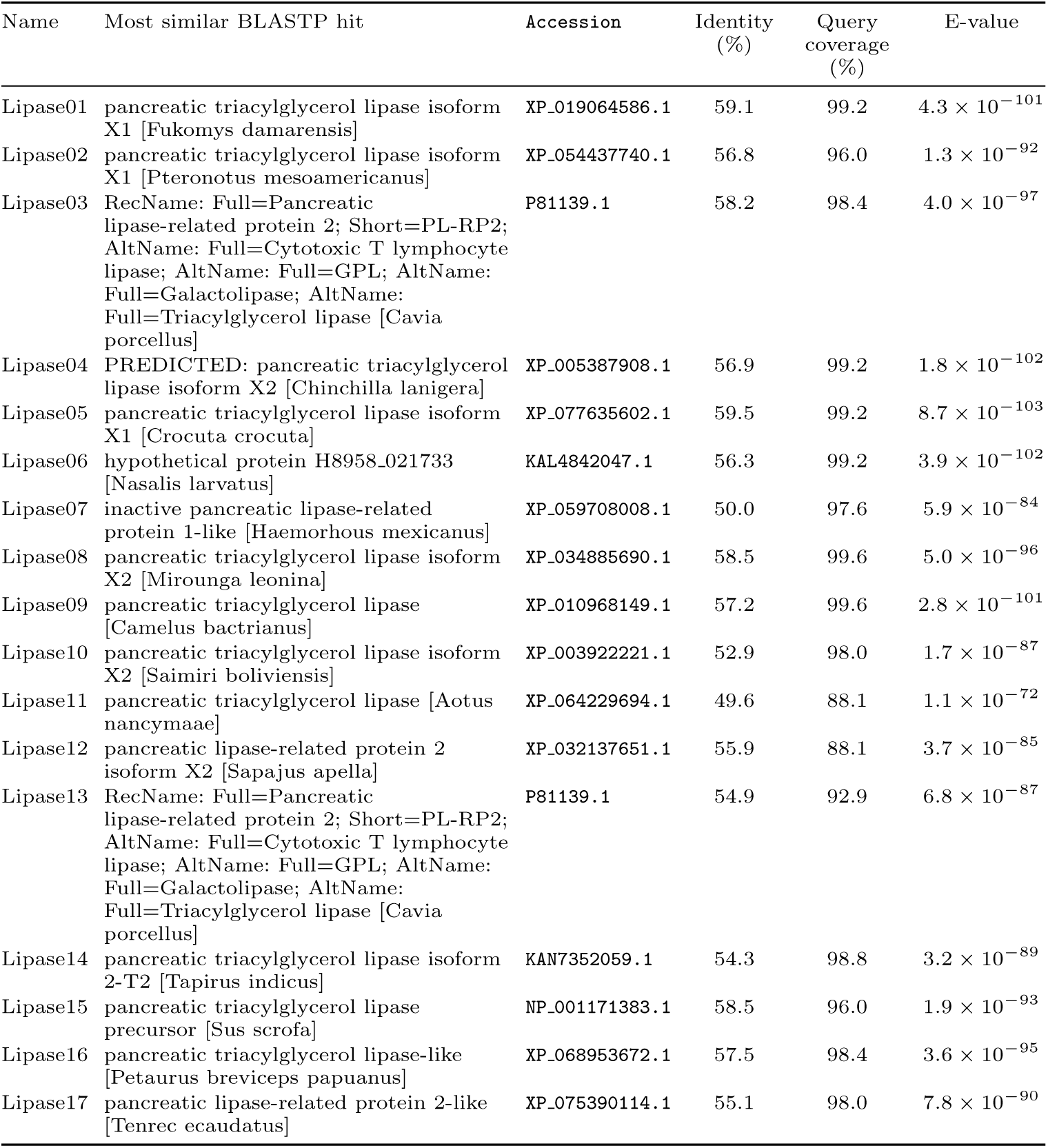
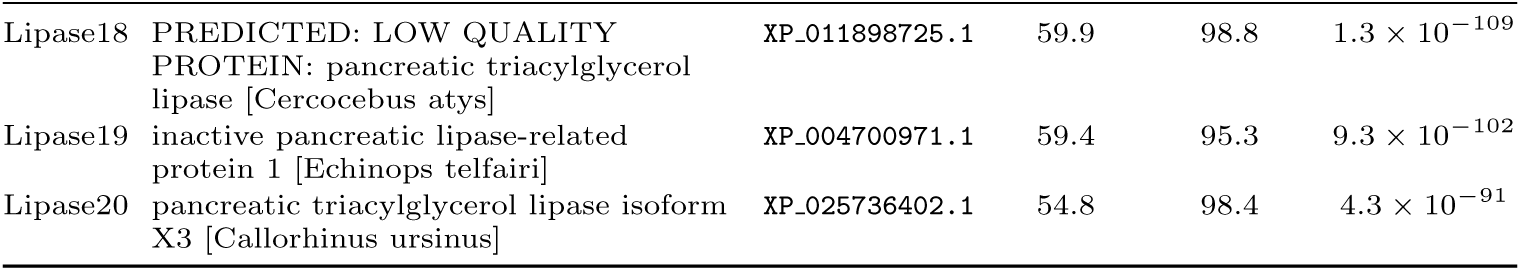
Most similar BLASTP hit for each EnzymeArt-designed lipase sequence, selected by maximum percent identity using NCBI web BLASTP results from 26 June 2026. Hit title, accession, identity, query coverage and E-value correspond to the selected hit.

### 7.13 Designed lipase sequences generated by EnzymeArt

**Table 27:**
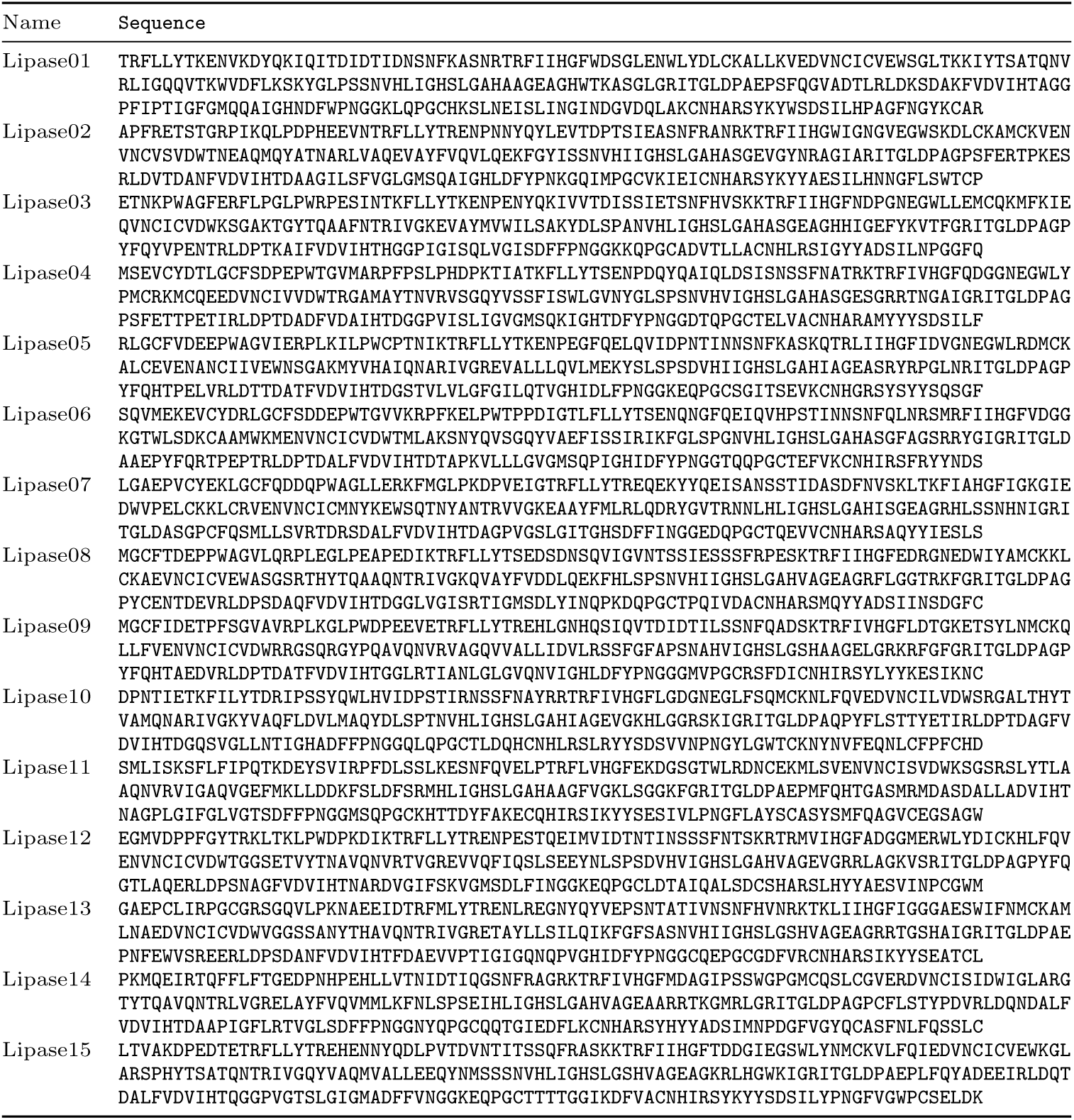
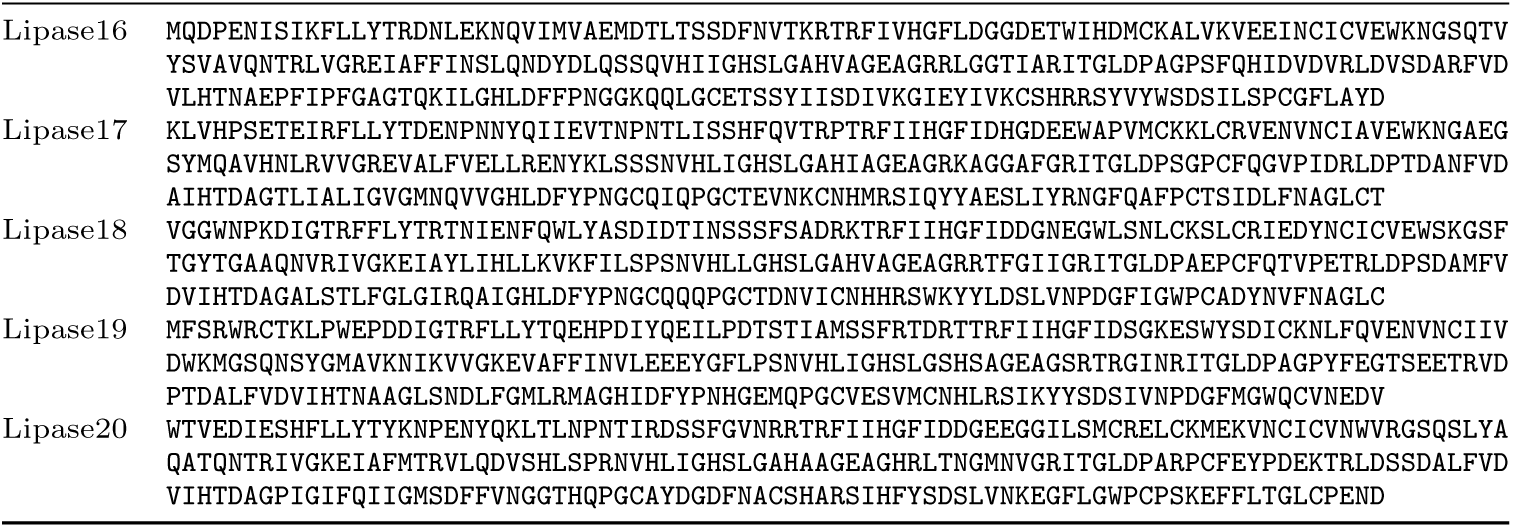
Sequences of EnzymeArt-designed lipases.

